# Multivalent DNA-encoded lectins on phage enable detecting compositional glycocalyx differences

**DOI:** 10.1101/2023.12.13.571601

**Authors:** Guilherme Meira Lima, Zeinab Jame Chenarboo, Mirat Sojitra, Susmita Sarkar, Eric J. Carpenter, Claire Yi-Ling Yang, Edward Schmidt, Justine Lai, Alexey Atrazhev, Danial Yazdan, Chuanhao Peng, Elizabeth Anne Volker, Ray Ho, Gisele Monteiro, Raymond Lai, Lara K. Mahal, Matthew S. Macauley, Ratmir Derda

## Abstract

Selective detection of disease-associated changes in the cellular glycocalyx is a foundation of modern targeted therapies. Detecting minor changes in the density and identity of glycans on the cell surface is a technological challenge exacerbated by lack of 1:1 correspondence between cellular DNA/RNA and glycan structures on cell surface. We demonstrate that multivalent displays of up to 300 lectins on DNA-barcoded M13 phage on a liquid lectin array (LiLA), detects subtle differences in composition and density of glycans on cells *ex vivo* and in immune cells or organs in animals. For example, constructs displaying 73 copies of diCBM40 lectin per 700×5 nm virion (φ-CBM73) exhibit non-linear ON/OFF-like recognition of sialoglycans on the surface of normal and cancer cells. In contrast, a high-valency φ-CBM290 display, or soluble diCBM40, exhibit canonical progressive scaling in binding with increased epitope density; these constructs cannot amplify the subtle differences detected by φ-CBM73. Similarly, multivalent displays of diCBM40 and Siglec-7 detect differences in the glycocalyx between stem-like and non-stem populations in cancer cells that are not detected with soluble lectins. Multivalent display of lectins on M13 scaffold with protected DNA inside the phage offer non-destructive detection of minor differences in glycocalyx in cells *in vitro* and *in vivo* not feasible to currently available technologies.

## Introduction

Identifying disease-associated changes in the cellular glycocalyx is necessary for the effective targeting of drugs to such cells and diagnosis of diseased cells. Modern cell-targeting paradigms include diverse clinical modalities such as antibody-drug conjugates (ACDs)^1^, chimeric antigen receptor (CAR) T cells^2^, and targeted radiopharmaceuticals^3^. An ideal disease-associated target epitope should exhibit “all-or-none expression”, i.e., high on disease-associated cell and zero on all healthy cells. A greater challenge in targeting arises if the epitopes are ubiquitous. For example, most cells in the human body present low-to-moderate levels of sialic acid in their glycocalyx, whereas tumors are often hypersialylated^4^. Although the distribution of sialic acids and many other cellular epitopes represent a continuous spectrum rather than the ideal “all-or-nothing expression”, the immune system has evolved the capacity to be triggered by expression levels above a non-zero threshold density of such epitopes^5^. Classical IgM-mediated responses, such as complement, are triggered in cells that express antigen above a certain non-zero threshold and not in cells with moderate and low expression^6^. Recognition of multivalent cell-surface epitopes is now firmly established to be the key requirement for achieving non-zero target thresholding. Several research groups have confirmed that multivalent interactions can indeed target cells that express high but not medium or low density of target receptors^7, 8^.

Glycans are ubiquitously present on the surface of cells across all kingdoms of life^9^. Both the binding of glycans to glycan-binding proteins (GBP) at cell interfaces and the interactions of cells with foreign organisms critically modulate cell signaling pathways^10^. Changes in the structure of glycans present in the glycocalyx of human cells have been linked to a variety of diseases^11, 12^. In cancer, tumor cells modify the amounts and compositions of glycans on their surface to escape recognition by the immune system and to promote metastasis^12^. Examples of aberrant glycosylation include increased sialylation and truncation of *O*-linked glycans on tumors^12^. Sequencing of cellular DNA or RNA cannot infer the composition of the glycocalyx because glycan synthesis is not DNA-encoded and the expression of glycosyltransferase genes only weakly correlates with the composition of glycocalyx^13, 14^. Decoding the composition of glycocalyx is traditionally performed by mass spectrometry^15, 16^ or printed lectin-array methods^17-19^; both require destructive pre-processing steps. Fluorescently labeled lectins^20, 21^ and lectin-functionalized liposomes^22, 23^ can detect the presence or absence of specific glycan motifs in the glycocalyx of live cells *in vitro* and *in vivo*; such technology is limited by the multiplexing capacity of fluorescently labeled lectins as the intensity of different fluorophores cannot be readily compared to one another. Covalent attachment of DNA to lectins^24-26^ could, in principle, increase the multiplexing capacity of lectins, however fundamental drawbacks remain: (i) DNA-tagged lectins are not *in vivo-* compatible due to the nuclease sensitivity of the DNA-tags; (ii) Soluble DNA-tagged lectins do not have the multivalent presentation of GBPs on the surface of cells; multivalent presentation of lectins can critically impact their ability to recognize glycans on the cell surface^27^. Here, we demonstrate that DNA-encoded multivalent display of lectins on a carrier that protects the DNA not only overcomes the above limitations but also reveals non-linear cooperative cellular recognition events that are possible to observe only by multivalent moieties.

Multivalent presentation is critical in the discovery of physiologically relevant lectin-glycan interactions^28^. Knowledge built on observations emanating from multivalent display of glycans and lectins on slide-based arrays serves as foundation of the entire field of glycobiology.^29^ Profiling of the glycocalyx on live cells is possible by individual (non-encoded) lectins and lectin conjugates. However, no technology today offers the capacity to combine multivalent presentation of lectins with DNA-encoding capacity for glycocalyx profiling *in vivo*. To address this need, we take advantage of the robust DNA-barcoded M13 bacteriophage and use the SpyCatcher-SpyTag technology pioneered by the Howarth group^30, 31^ to build *in vivo*-compatible platform for controlled multivalent display of covalently-attached lectins on the surface of bacteriophage that can be decoded by next-generation sequencing (NGS). Here, we show that the multivalent liquid lectin array (LiLA) platform can effectively profile the composition of the glycocalyx on cells *in vitro* and *in vivo* using mice as a model organism.

## Results

### Synthesis and validation of lectin-phage conjugates

We previously employed SpyTag technology to display tetrameric or PEGylated proteins on phage^32^. Here, we expand this approach to display roughly a thousand copies of the third generation SpyCatcher (Sc) protein^31^ on phage and use this construct to display lectins containing the 16 amino acid SpyTag (St) peptide.^33^ We used St-Sc approach to immobilize lectins on phage to ensure consistent glycan-binding performance of lectins immobilized in a uniform fashion^34^. Incubation of M13 phage with 0.1 mM *N*-hydroxysuccinimide-iodoacetamide (NHS-IA) acetylated 1200 out of 2700 pVIII phage proteins (Fig. 1a,b). Subsequent addition of the S49C mutant of SpyCatcher^31, 35^ at pH 8.3, consumed 60% of pVIII-IA as confirmed by MALDI-TOF; we capped the unreacted 40% of pVIII-IA with 5 mM cysteine. MALDI observations implied that ∼700 copies of the SpyCatcher protein have been ligated per phage (Fig. 1b,c). Incubation of this construct with a SpyTag-5-carboxytetrametrylrhodamine (TAMRA) dye and fluorescence measurements estimated 1300 TAMRA molecules per phage particle (Fig. 1d, Supplementary Fig. 1). Based on the combined MALDI and fluorescence results, we assumed that ∼1000 copies of Sc are displayed on a 700 nm x 5 nm virion and used φ-Sc_1000_ to describe this construct. Notably, 1000 copies approaches van der Walls packing density of the Sc protein on phage (Fig 1c). A one hour incubation of φ-Sc_1000_ with diCBM40 protein fused to SpyTag (St-diCBM40) ligated 290 copies of this lectin to phage yielding phage constructs which we referred to as φ-CBM_290_ (Fig. 1e, Extended Data Fig. 1, and Extended Data Fig. 2a,b). Soluble diCBM40 lectin robustly binds to α2-3 and less to α2-6-linked sialosides^36, 37^ and myeloid leukemia cell line U937 expresses these sialoglycans (Supplementary Fig. 2). A previously published phage binding assay^38^ confirmed that binding of φ-CBM_290_ to U937 cells was 1000-fold higher than binding of blank (unmodified) clones (Fig. 1f,g; Supplementary Fig. 3). A fold change (FC) difference between normalized read counts for DNA of phage binding to U937 cells vs. input phage measured by next-generation sequencing (NGS) yielded the same factor of 1000 between φ-CBM_290_ and blank phage (Fig. 1h).

**Fig. 1.**
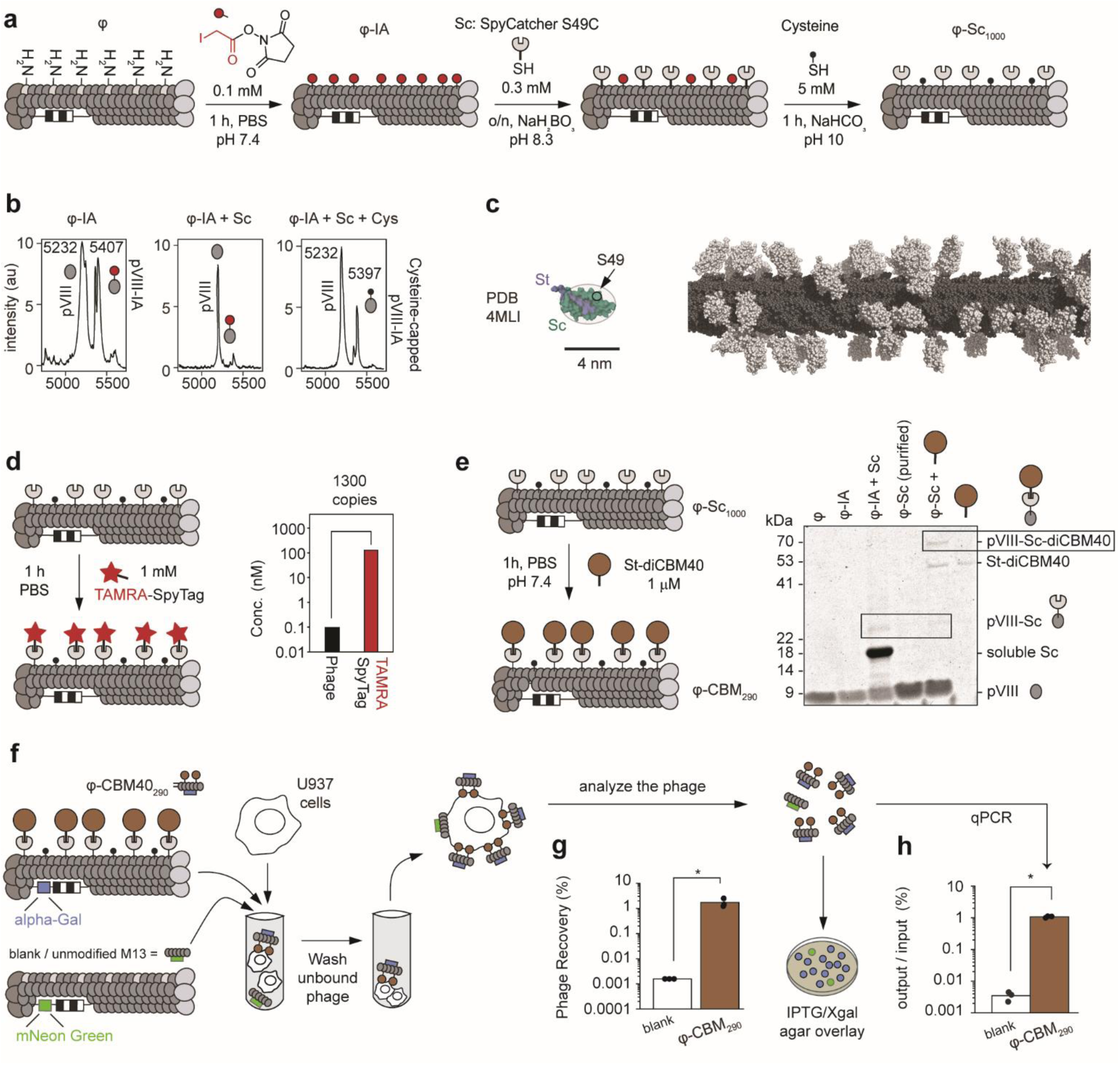
Synthesis and characterization of SpyCatcher-displaying phages (φ-Sc_1000_). **a,** Scheme of φ-Sc synthesis via chemical conjugation of SpyCatcher on phage. **b,** MALDI spectra of φ-Sc synthesis steps. **c,** Spacing of Sc (colored in white, PDB: 4MLI) on the M13 virion (colored in gray, PDB: 2MJZ). **d,** Quantification of φ-Sc fluorescence after reaction with TAMRA-SpyTag. **e,** SDS-PAGE image of synthesis of φ-CBM_290_ phages. **f,** Scheme of cell panning, colorimetric plaque assay and deep sequencing used for rapid validation of φ-CBM_290_ phages in the presence of blank phages. **g,** Recovery of φ-CBM_290_ and blank phages as measured by plaque assay. Values represent mean of *n* = 3, Mann-Whitney U Test. **h,** Recovery of φ-CBM_290_ and blank phages as measured by the ratio of DNA reads after (output) and before (input) cell panning. Values represent mean of *n* = 3. Mann Whitney U Test. In **g-h**, all respective n values are independent technical replicates. * *P* < 0.05.

Encouraged by these results, we conjugated sialic acid binding immunoglobulin-type lectin 7 (Siglec-7) and a R124A Siglec-7 mutant (Siglec-7R), yielding φ-Siglec7_20_ and φ-Siglec7R_30,_ respectively (Supplementary Fig. 4), and as a baseline reference we modified phage with maltose binding protein (MBP), yielding the φ-MBP_140_ construct (Extended Data Fig. 2a,b). Using the same clonal phage binding assay (Fig. 2a), we observed a 50-fold increase in binding of φ-Siglec7_20_ to carbohydrate sulfotransferase 1 overexpressing cells (CHST1-U937) when compared to WT U937, confirming the known role of 6-*O*-Gal-sulfation on an underlying galactose in enhancing binding to Siglec-7. Disruption of the essential Arg124 residue in Siglec-7R mutant ablated binding of φ-Siglec-7R_30_ construct to a level statistically indistinguishable from the baseline (Fig. 2b). Binding of both φ-CBM_290_ and φ-Siglec7_20_ to cytidine monophosphate *N-*acetylneuraminic acid synthase (CMAS) deficient (CMAS^KO^) U937 cells was decreased (Fig. 2b) because CMAS^KO^ cannot produce sialic acid; binding of φ-CBM_290_ to CMAS^KO^ was minor (< 5% of binding of φ-CBM_290_ to WT) and reminiscent of published observations of binding of sialic acid binding lectins to CMAS^KO^ cells^39^. In all these experiments, we observed no differences in binding between φ-MBP_140_ and blank phages (Fig. 2b), which was expected because mammalian cell glycocalyx does not contain maltose. Cell binding capacities of individual lectin-modified phage clones were closely reproduced when all the DNA-barcoded phages and controls were mixed (Fig. 2c) and analyzed by NGS (Fig. 2d). As previously (Fig 1g and 1h), FC enrichment measured by NGS correlated closely with absolute recoveries measured by clonal phage-binding assays (compare Fig. 2b and 2e, FC = CPM_out_/CPM_in_). NGS confirmed equivalence of blank and φ-MBP_140_ phage and re-confirmed that MBP is a suitable benchmark for future mammalian cell binding assays. In contrast, phages with Sc not ligated to any protein consistently showed a 100-fold increase binding over the baseline. This non-specific binding of Sc to cells has been reported^40^ and we observed that the risk of non-specific binding is reduced as long as all Sc are capped by an inactive protein (here MBP). Agreement of binding measured for individual phage clones and within a complex mixture established the robustness of NGS as a readout for the LiLA technology.

**Fig. 2.**
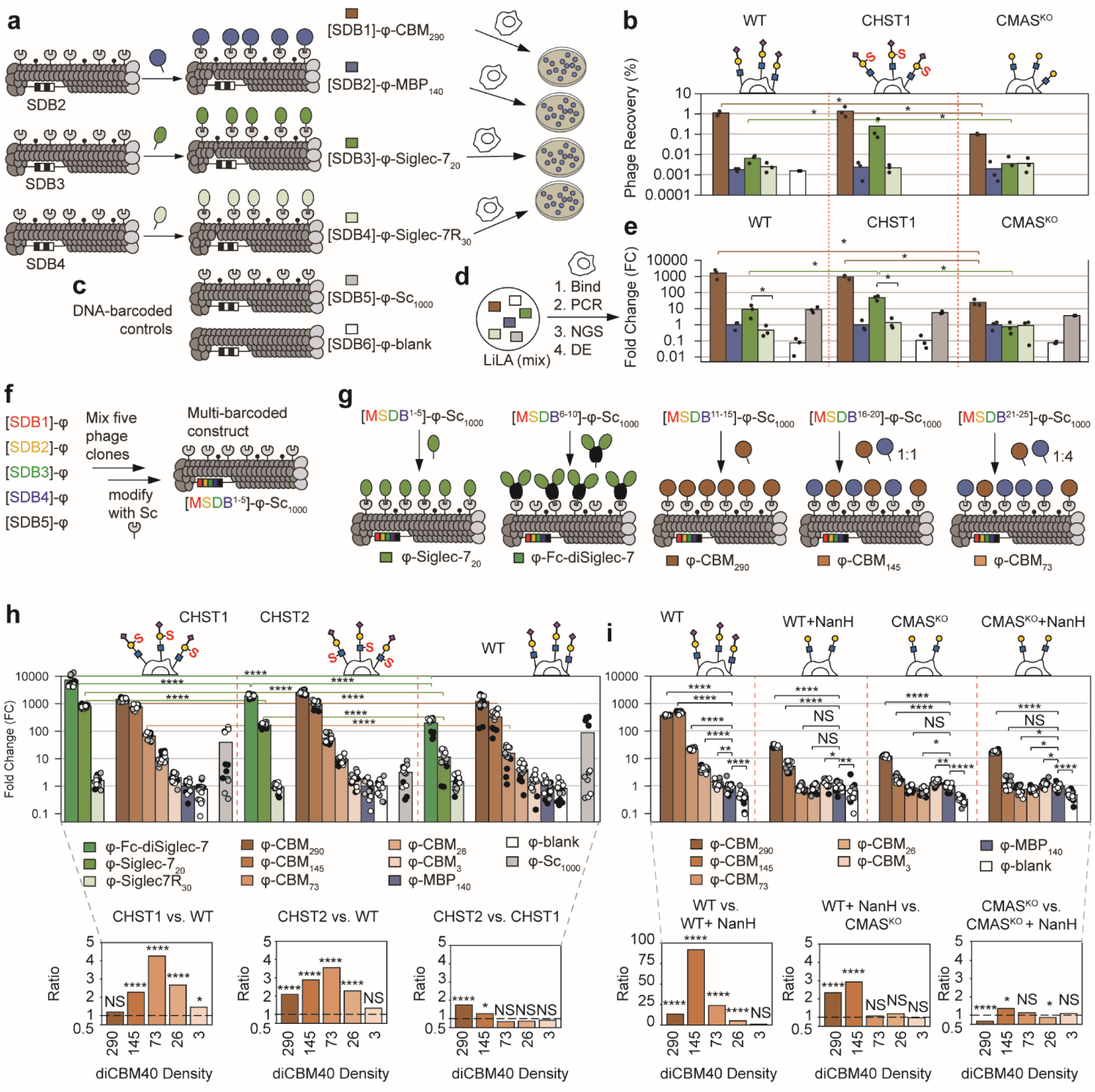
Characterization of the LiLA. **a,** Clonal binding assay characterizes binding of individual phage constructs to various U937 cells (**b**). Values represent mean of *n* = 3 technical replicates, Mann-Whitney U Test **c,** Incorporation of DNA-barcoded controls in the LiLA. Each component of the library is associated with a specific DNA barcode. **d,** Phage constructs are probed as a mixture against a given cell line. DNA barcodes associated with each phage construct are decoded by deep sequencing and used to quantify binding of each component of the library. **e,** Binding profile of LiLA on WT, CHST1 and CMAS^KO^ U937 cells as measured by deep sequencing. FC of φ-MBP_140_ clones was set to one. Values represent mean of *n* = 3 technical replicates, Mann-Whitney U test. **f,** A defined set of five DNA barcodes are associated with φ-Sc_1000._ **g,** φ-Sc_1000_ phages containing a mixture of DNA barcodes are mixed with a lectin of interest to result in a library containing all constructs associated with five different DNA barcodes each. **h,** Binding profile of expanded LiLA on WT, CHST1 and CHST2 cells as measured by deep sequencing. FC of φ-MBP_140_ clones was set to one. Values represent mean of *n* = 3 (technical replicates) x 5 (DNA barcodes per construct), Mann-Whitney U Test. Bottom plot represents ratio of FC values between the different cell lines. **i,** Binding of φ-CBM/φ-MBP phages displayed at various densities to U937 cells treated or not treated with neuraminidase as measured by deep sequencing. FC of φ-MBP_140_ clones was set to one. Values represent mean of *n* = 3 (technical replicates) x 5 (DNA barcodes per construct), Mann-Whitney U Test. Bottom plot represents ratio of FC values between the different cell lines treated or not with neuraminidase. In **h-i**, each color of scatter plot represents one independent experiment, and each technical replicate contains 5 DNA barcodes associated with each phage. NS, *P* > 0.05; * *P* < 0.05, ** *P* < 0.01, **** *P* < 0.0001.

### Specific density of lectins on phage allows sensitive discrimination of differences of sialoepitopes in glycocalyx

In nature, glycan-GBP interactions are modulated by avidity and the multivalent presentation of glycans and GBPs^41^. Immune cells that survey sialo-epitope density on tumor cells can turn off in response to increase to modest changes in density of sialic acids^42^. This multivalent interaction depends both on sialo-epitopes density on cancer cells and density of sialic acid binding proteins on immune cells^42^. To probe these phenomena, we expanded LiLA to include different multivalent presentations of densities of diCBM40 lectin and tested whether different displays can yield a more precise discrimination in density of sialic acid epitopes on cells when compared to soluble diCBM40 lectin. We produced a DNA-encoded “CBM density gradient” with distinct copy numbers of diCBM40 lectin on phage; each density was encoded by multiple DNA barcodes (MSDB Fig. 2g) to increase the confidence of NGS analysis. Starting from a mixture of 5 clonal phage modified by Sc, we conjugated a defined copy number of diCBM40 by controlling concentration of St-diCBM40 and capping unreacted Sc by St-MBP or St. Increasing the concentration of St-MBP proportionally decreased copy number of diCBM40 displayed on phage (Extended Data Fig. 3a, b) giving rise to φ-CBM_N_ with N=290, 145, 73, 26, or 3 copies of diCBM40 lectin on phage. NGS analysis of binding of CBM-gradient (φ-CBM_N_) to U937 cells showed an exponential surge in binding with increase of N (Fig. 2h). Binding of individual phage clones to WT U937 re-confirmed that binding of φ-CBM_N_ phage clones increased exponentially as N increased (Extended Data Fig. 3c). We observed the same pattern for φ-Siglec-7_N_ phages displaying three different densities of Siglec-7 (Extended Data Fig. 3d). We further observed that phages that display dimeric Siglec-7-Fc chimera (φ-Fc-diSiglec-7) exhibits a consistent increase in binding when compared to φ-Siglec-7 displaying monomeric protein. Binding of φ-Siglec-7 was the greatest to CHST1 cells, followed by CHST2, and WT U937 cells (Fig. 2h) corroborating reported preference of Siglec-7 for 6-*O*-Gal sulfation and 6-*O*-GlcNAc sulfation installed by CHST1 and CHST2, respectively^43^. Binding of φ-Fc-diSiglec-7 to the CHST1, CHST2, and WT cells mirrored these trends (Fig. 2h) but shows consistent 10-fold increase in binding of φ-Fc-diSiglec-7 (i.e. multivalent display of Siglec-7 dimers on phage) when compared to φ-Siglec-7 (i.e., multivalent display of Siglec-7 monomers on phage). Comparison of the ability of each multivalent construct to discriminate WT and CHST1/2 cells, remarkably uncovered that binding of φ-CBM_73_ was 5 times higher to CHST1 U937 than WT U937 cells, but the difference between CHST1-U937 and WT U937 was indistinguishable when measured with high density φ-CBM_290_, low density φ-CBM_3_ and soluble diCBM40 (Fig. 2h). Discrimination of CHST1-U937 and WT U937 by φ-CBM_N_ constructs is bimodal with clear maximum at N = 73 and progressively tapering out both as density decreases (N = 3, 26) and increases (N = 145, 290). Similarly, φ-CBM_73_ was the most effective in discriminating CHST2 from WT and similar bimodal trend was observed for all φ-CBM_N_ constructs. A plausible biochemical rationale for subtle difference between CHST1 and WT is that 6-*O*-Gal sulfation could prevent ST6-β-galactoside-α-2,6-sialyltranferase-1 (ST6GAL1) from installing α-2,3 sialylation on glycans, resulting in more α2-3-linked sialosides on cells. The observation that only “medium” density display of diCBM40 can detect the differences between these cells highlight a critical interplay between the lectin density and their ability to differentiate subtle differences in glycocalyx on the surfaces of cells.

The binding of CBM-gradient to cells plummeted non-linearly, across orders of magnitude as sialo-epitopes were progressively removed. Binding of φ-CBM_N_ to WT U937 cells spanned 3 orders of magnitude: from FC=400 for high-density φ-CBM_290_ construct to FC=2 for low-density φ-CBM_3_ construct and FC=1 for φ-MBP control (Fig. 2i). Removal of α2-3, 2-6, and 2-8 sialic acid from U937 cell surface by the neuraminidase NanH enzyme ablated binding of φ-CBM_73_, φ-CBM_26_, and φ-CBM_3_ to the levels indistinguishable from φ-MBP control. Binding of φ-CBM_145_ to NanH-treated U937 decreased by factor of 100 when compared to WT U937 but it was detectable above the background (Fig. 2i). The binding of φ-CBM_145_ was fully ablated in CMAS^KO^ U937, suggesting that the concentration of sialic acid on NanH-treated WT U937 cells is not zero and it is detectable by φ-CBM_145_ and φ-CBM_290._ Interestingly, the highest density φ-CBM_290_ exhibited a significant FC=10 binding even in CMAS^KO^ U937 cells and further treatment of CMAS KO U937 cells with NanH to remove any potential exogenously introduced sialic acid from cell surface did not reduce this binding (Fig. 2i). Note that FC=10 represents 2.5

% of the signal for WT U937 cells (FC=400) and it is robustly detectable owing to a high dynamic range of the NGS-powered LiLA assay. Such low signal might evade detection in an assay with narrower dynamic range (e.g., staining of cells by soluble diCBM40 lectin). DNA-encoded constructs displaying different valency of diCBM40 per phage, thus, serve different purposes as probes of sialic acid epitopes in a glycocalyx. The high density φ-CBM_290_ construct offers wide dynamic range for detecting broad changes in sialylation density. Whereas binding of φ-CBM_73_ is strongly attenuated and turned off (Fig. 2i) in response to modest decreases in the sialo-epitope density; such non-linear attenuation is a hallmark of cooperative interactions. To rule out NGS or PCR artefacts in these experiments, all observations in Fig. 2g and 2h were supported by MSDBs constructs (i.e. 5 DNA barcode sequences per construct). To explore further this interplay of sensitivity and cooperativity in LiLA and CBM-gradient, we applied this platform to a diverse set of cell types *in vitro*, *ex vivo,* and *in vivo*.

### Analysis of glycocalyx in heterogenous cell populations by LiLA+FACS

Phage-displayed technologies based on M13 phage are uniquely positioned for detection of specific interactions in complex heterogeneous mixtures of cells using a simple 3-step protocol: (i) incubate M13-displayed library with complex mixtures of cells; (ii) use fluorescence-activated cell sorting (FACS) to isolate the desired population of cells; and (iii) analyze the associated ligands/barcodes by NGS. For example, we analyzed the binding profile of LiLA on human peripheral blood mononuclear cells (PBMCs) obtained from the blood of a healthy human volunteer. White blood cells were incubated with LiLA and a cocktail of anti-CD3, CD19, CD56, and CD14 antibodies, and sorted by FACS to isolate populations of T-cells (CD3^+^), B-cells (CD19^+^), NK cells (CD56^+^), and monocytes (CD14^+^) (Fig. 3a, Supplementary Table 3, and Supplementary Fig. 5). Sequencing of DNA barcodes associated with each cell type revealed a gradient of enrichment of diCBM40 constructs on the four selected PBMCs: T-cells have the highest binding to diCBM40, followed by monocytes, NK-cells, and B-cells (Fig. 3b). On the other hand, binding of φ-CBM_73_ and φ-Siglec-7_20_ was not detectable and it was indistinguishable from φ-CBM_26_, φ-CBM_3_, and controls φ-MBP_140_, φ-Sc_1000,_ and blank φ phages (Fig. 3b). Staining of mouse splenocytes *ex vivo* using FACS→LILA showed greatest enrichment of φ-CBM_290_ and φ-Siglec-7_20_ to T-cells relative to B-cells, similar to what was observed with human leukocytes (Fig. 3d). LiLA can also associate with already sorted PBMC cells (FACS→LiLA) (Supplementary Fig. 6a). The results were similar but the quality of signal deteriorated (Supplementary Fig. 6b) because sequential FACS→LILA introduced more stress (washing and cell handling) and yields fewer cells for analysis (Supplementary Table 3). The staining intensity of PBMC obtained using soluble Siglec-7-Fc proteins correlated both with φ-CBM_290_ binding and φ-Fc-diSiglec-7 binding (Extended Data Fig. 4) and similar to reported binding of tetrameric Siglec-7-Fc-streptactin to immune cells^44^. The decision to apply LiLA before or after sorting should thus consider whether cells are stable after isolation. For more delicate cells, a streamlined LiLA+FACS is preferred.

**Fig. 3.**
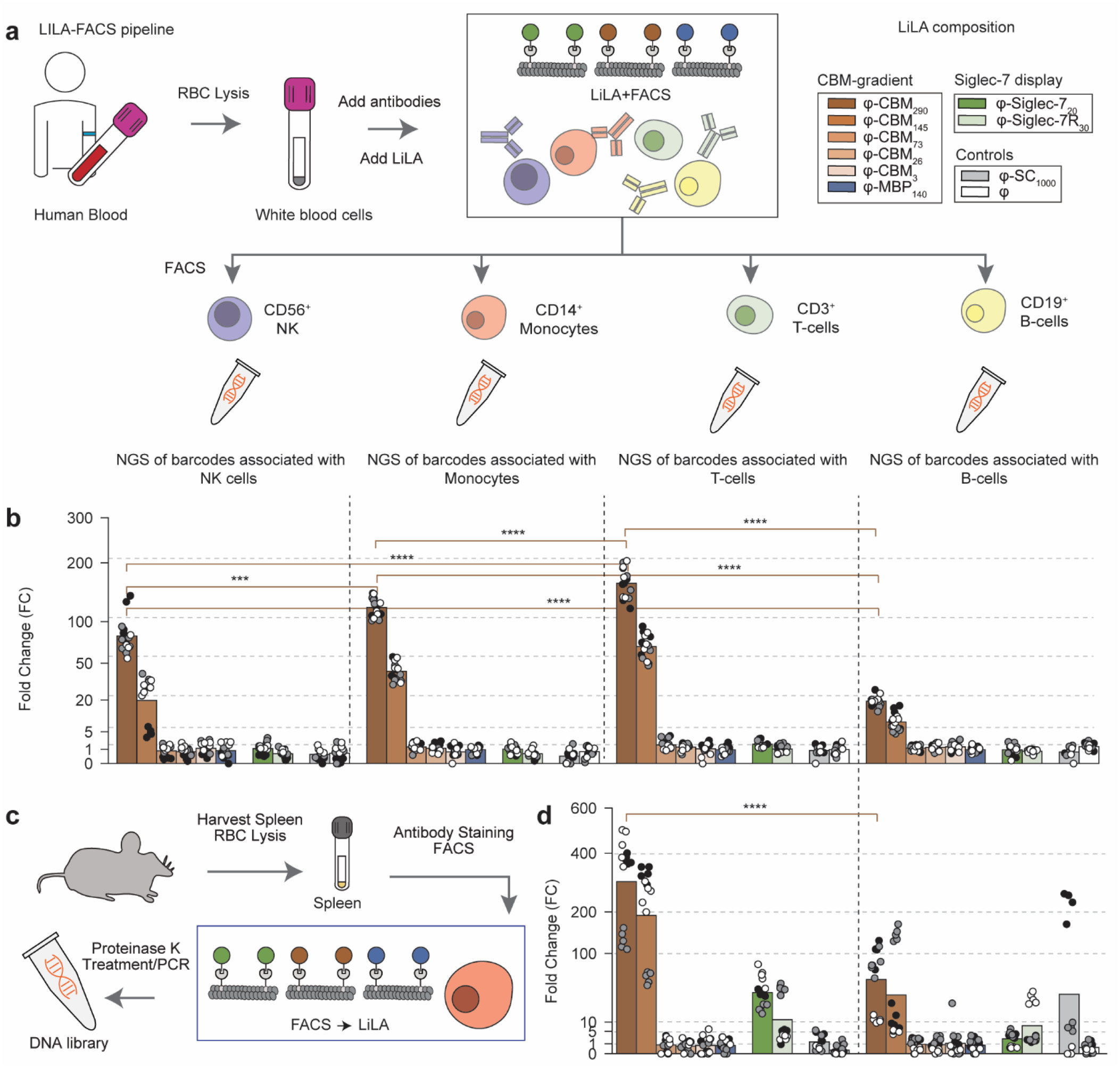
Glycome profiling of immune cells by LiLA-FACS. **a,** Scheme of the LiLA+FACS approach using human PBMCs. White blood cells were obtained by drawing human blood followed by RBC lysis treatment. Desired immune cells were stained with antibodies, mixed with LiLA, and submitted for FACS. **b,** Binding profile of human NK-cells, monocytes, T-cells, and B-cells to LiLA. FC of φ-MBP_140_ clones was set to one. Values represent mean of *n* = 3 (technical replicates) x 5 (DNA barcodes per construct). PBMCs were collected from a single volunteer. **c,** Scheme of the FACS→LiLA approach using murine splenocytes. White blood cells were obtained by homogenization of the murine spleen followed by RBC lysis treatment. Desired immune cells were stained with antibodies and submitted for FACS. Isolated cells (i.e. T-cells/B-cells) were individually mixed with LiLA. **d,** Binding profile of murine T and B cells to LiLA. FC of φ-MBP_140_ clones was set to one. Values represent mean of *n* = 3 (biological replicates) x 5 (DNA barcodes per construct). *** *P* < 0.001, **** *P* < 0.0001.

Comparison of LiLA results to staining of cells by soluble lectins highlights not only similarities (Extended Data Fig. 4), but also abrupt changes in the binding of lectins to cells in response to their changes in multivalent presentation. Phages displaying monomeric Siglec-7 exhibit nearly undetectable binding to PBMCs, but phage displaying the same Siglec-7 as a dimer (Fc-diSiglec-7) exhibits a non-linear surge in binding (Extended Data Fig. 4). Further comparison of concentration regimes and dose-response of multivalent lectin display on phage, components of LiLA, and soluble lectins reinforces their fundamental difference in interaction with cells. Binding of soluble fluorescent diCBM40 to U937 cells at 100 pM concentration or below is indistinguishable from a control (Supplementary Fig. 2) but binding of φ-CBM_290_ to U937 cells at 58 pM “per protein” concentration yields a 1000-fold enrichment compared to control constructs (Fig. 2b). In flow cytometry, across its practical range, a 2-fold increase in concentration of soluble diCBM40 leads to a predictable 2-fold increase in staining intensity (Supplementary Fig. 2). Constructs φ-CBM_290_ contains twice the protein of φ-CBM_145_; but this apparent 2x increase translated to either no change in binding to highly sialylated U937 cells, or a 10x increase in binding to cells with low sialic acid density (NanH-treated U937 or CMAS^KO^ U937 cells, Fig. 2i). Expanding this analysis to all tested PBMCs and variants of U937 cells unveils both linear and non-linear (cooperative) change in response to displayed sialic acid density. Binding of the φ-CBM_145_ construct decreased gradually as the density of sialic acid on cells decreased, in a manner expected from staining with soluble diCBM40 lectin (Fig. 4a). In contrast, binding of φ-CBM_73_ to cells ceased abruptly once density of sialic acids on cells passed a certain threshold (Fig. 4b). Intermediate density display of diCBM40 on phage offers “non-linear” amplified detection of subtle changes in density of sialylation on the surface of cells. This very strong positive cooperativity with respect to copy number of lectins per phage makes it possible for multivalent lectin constructs, components of LiLA, to amplify subtle change in glycan display density to a non-linear, abrupt ON/OFF binding response.

**Fig. 4.**
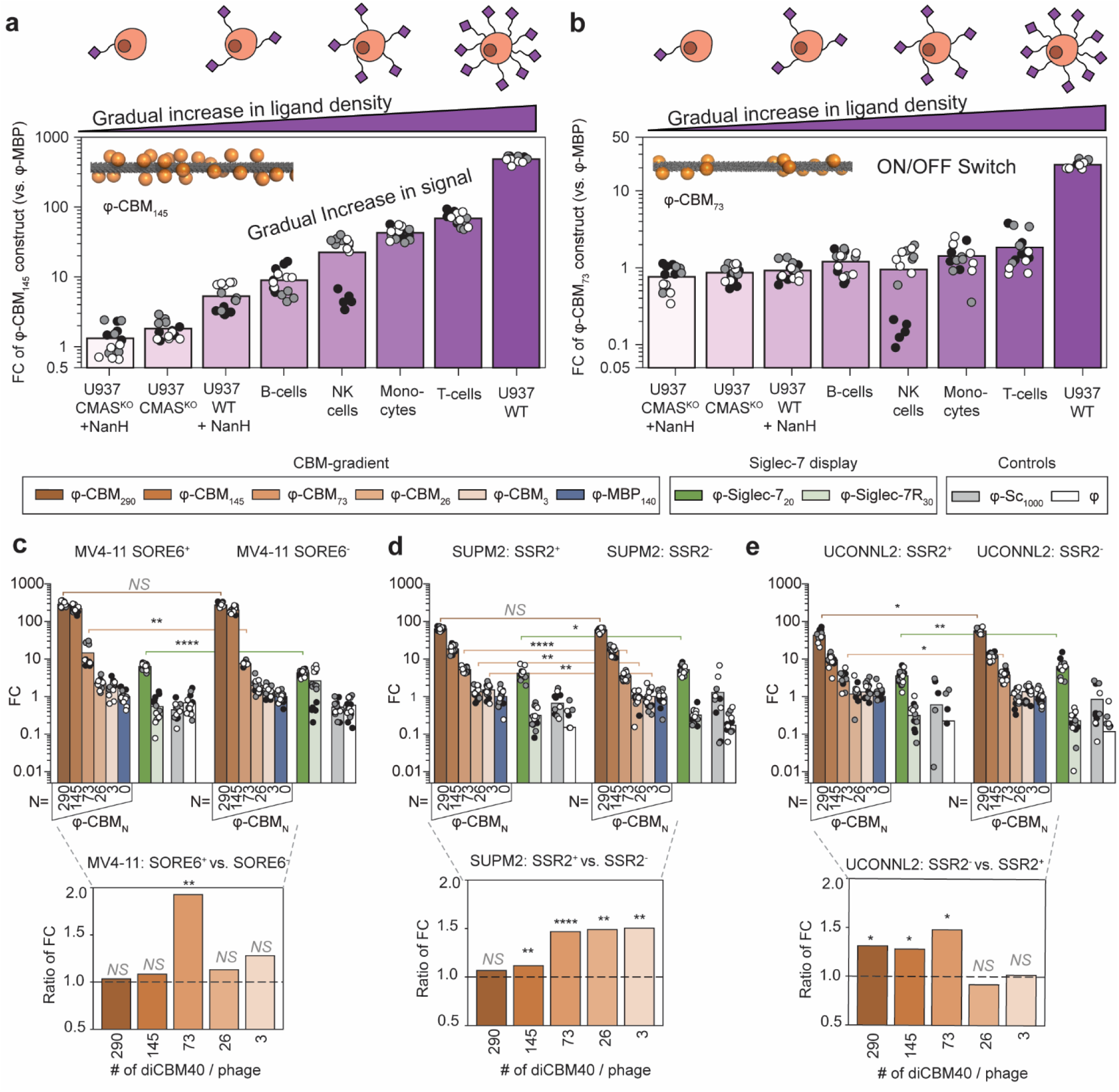
Enrichment of φ-CBM_145_ and φ-CBM_73_ on various cell lines and *in vitro* test *of* LiLA on CSC. **a,** Linear binding increase of φ-CBM_145_ as a function of sialic acid density on the surface of various cells. **b,** Non-linear response of φ-CBM_73_ when binding responses were plotted against the same cell lines. FC values were extracted from binding of LiLA to U937 cells (Fig. 2i) and human PBMCs (Fig. 3b). **c,** Binding of LiLA on MV4-11 SORE6^+^ and SORE6^-^ cells as measured by deep sequencing. FC of φ-MBP_140_ clones was set to one. Values represent mean of *n* = 3 (technical replicates) × 5 (DNA barcodes per construct), Mann-Whitney U Test. **d,** Binding of LiLA on SUPM2 T-cell lymphoma cell lines. **e,** Bindinf of LiLA on UCONNL2 cells. FC of φ-MBP_140_ clones was set to one. ‘+’ = CSC and ‘-’ = NSCC. Values represent mean of *n* = 3 (technical replicates) × 5 (DNA barcodes per construct), Mann-Whitney U Test. In **c-e** bottom plots represent ratio of FC values between the different cell lines NS, *P* > 0.05; * *P* < 0.05, ** *P* < 0.01, **** *P* < 0.0001.

### Analysis of glycocalyx on cancer stem cells

To explore whether LiLA can detect subtle changes in glycocalyx composition of closely related cell types, we profiled cancer stem-like cells (CSC-like) and closely related non-stem cancer cells (NSCC) with LiLA. We focused on CSC/NSCC-like populations derived from the human acute myeloid leukemia (AML) cell line (MV4-11) and two anaplastic large cell lymphoma (ALCL) cell lines (UCONN-L2 and SUPM2). CSC-like cells were identified using the SOX2/OCT4 response element (SORE6) reporter^45, 46^ or SOX2 regulatory region-2 (SRR2) reporter^47, 48^; In all these cells, GFP^+^ cells with increased SOX2/OCT4 transcriptional activity mark CSC-like cells that exhibit increased tumor-forming capacity *in vitro* and *in vivo* when compared to GFP^-^ NSCC^45, 47, 48^. Panning of LiLA by GFP^+^/SORE6^+^ and GFP^-^/SORE6^-^ populations from MV4-11 cells (Supplementary Fig. 7a, b) detected a 1.5-fold increase of binding of φ-Siglec-7_20_ and ∼1.9-fold increase in binding of φ-CBM_73_ phages to SORE6^+^ MV4-11 cells when compared to SORE6^-^ counterparts (Fig. 4c). Unlike intermediate density φ-CBM_73_ construct, other φ-CBM_N_ constructs—high density N=290, and 145 and low-density N=25 and 3—exhibited no significant differences in binding (Fig. 4c). Results were corroborated by binding of individual lectin-decorated phage clones (Supplementary Fig. 7c). In CSC/NSCC population from ALCL SUPM2 cells binding of φ-CBM_73_, φ-CBM_26_ and φ-CBM_3_ (but not higher densities) was higher to CSC (SSR2^+^) when compared to NSCC (SSR2^-^) cells from the same population (Fig. 4d). UCONN-L2 populations were also discriminated by CBM-gradient, and binding of φ-CBM_73_ was the lowest to CSC compared to NSCC (Fig. 4e). Binding of LiLA to MV4-11 cells treated with antineoplastic drugs revealed synergistic effects of treatment on glycosylation. Binding of φ-CBM_N,_ where N=290, 145, 73, 26, and φ-Siglec-7_20_ to MV4-11 cells were ablated upon treatment with Azacitidine and Venetoclax whereas treatment of cells with any of the drugs alone produced less pronounced effects on glycosylation, as detected by LiLA (Supplementary Fig. 8). Combined observation highlights the capacity of LiLA to amplify gradual changes in sialylation density which can be a powerful detector in differentiating CSC from NSCC tumor cells and response of these cells to drug treatments.

### *In vivo* panning of LiLA

As with other M13-based platform technologies^38, 49, 50^, LiLA is compatible with *in vivo* panning. We injected a library of barcoded lectin-phage conjugates in C57BL/6J mice to test whether LiLA can detect differences in glycocalyx composition on the surface of cells *in vivo*. Similarly to previous reports, we tracked DNA-barcoded phage conjugates in the plasma, organs, and immune cells isolated from the spleen^32^, using PCR and NGS as previously described^51^. Unlike previous report, we did not amplify organ-isolated phage in bacteria and instead we isolated phage DNA cell from organ-homogenates directly using a recently published updated protocol^52^ (Fig. 5a). As described in previous sections (Fig 2f), *in vivo* LiLA experiments maximized the robustness of NGS analysis by encoding every phage conjugate by 5 distinct DNA barcodes. More than 90% of phages remained in circulation one hour after injection, 5% were distributed to the liver, and ∼0.1% to the spleen. Phage recovery from the heart, lungs, and kidneys was 0.001 – 0.01% but the isolated DNA from phage particles was still amenable to NGS (Fig. 5b). DNA barcodes associated with φ-CBM_290_ were significantly depleted from plasma and enriched in the lungs, heart, kidneys, spleen, and red blood cells (RBCs) (Fig. 5c). More than 90% of Fc-diSiglec-7 constructs were cleared from the plasma, and concomitantly enriched in the heart, lungs, kidneys, and RBCs (Fig. 5d). In contrast, phages that do not bind to sialic acid (φ-MBP_140_ and φ-Siglec-7R_30_) were not depleted from the plasma. Analysis of global enrichment in the organs suggests that these “non-binding” constructs are cleared from the plasma to the liver (Fig. 5c,d). Isolation of immune cells from splenocytes by FACS showed that φ-CBM_290_ and φ-Fc-diSiglec-7 phages enriched in T-cells relative to B-cells in line with *ex vivo* staining of human PBMCs (Fig. 3c) and mouse splenocytes (Fig. 3d). Aligned results from *in vivo* and *ex vivo* (Fig. 3d) analysis of mouse splenocytes confirms the previously unexplored possibility to study the precise composition of cellular glycocalyx on the surface of live cells inside live organism in a “non-destructive” fashion using multivalent liquid lectin array.

**Fig. 5.**
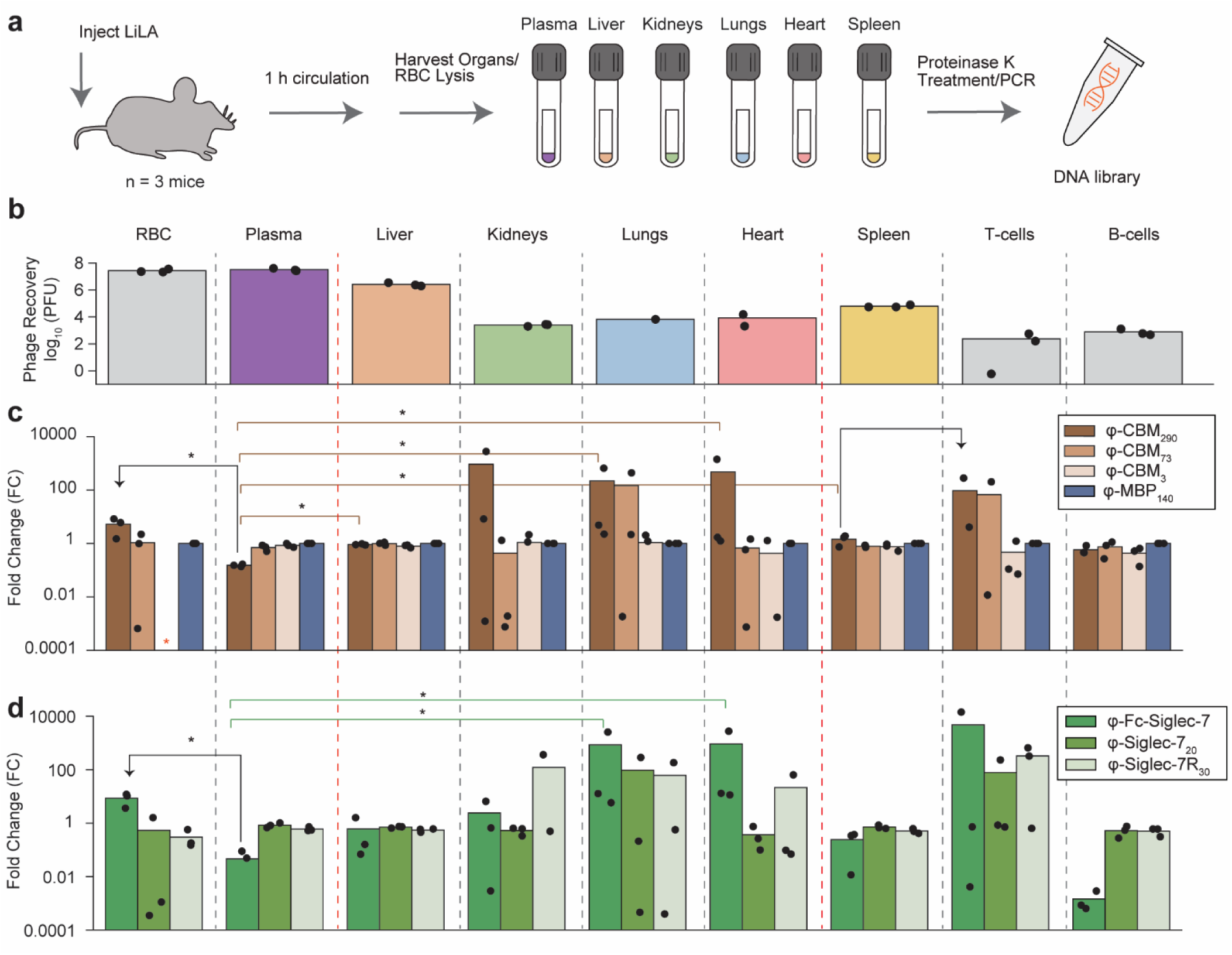
Injection of LiLA into mice. **a,** Scheme of *in vivo* panning procedure (*n* = 3 animals). **b,** Number of phage particles recovered from cells and organs as measured by qPCR **c,** Phage-enrichment of φ-CBM_290_, φ-CBM_73,_ φ-CBM_3,_ and φ-MBP_140_ on cells (left) and organs (right) relative to input sample as measured by deep sequencing. **d,** Phage-enrichment of φ-di-Siglec-7-Fc, φ-Siglec-7_20_, φ-Siglec-7R_30_ on cells (left) and organs (right) relative to input sample as measured by deep sequencing. Asterisk indicates no DNA reads associated with phage construct were found in the sample. In **c-d**, number of reads of all SDBs were averaged and normalized to averaged number of reads of SDBs associated with φ-MBP_140_. SDBs that do not appear in any replicate were excluded. Mann-Whitney U Test. * *P* < 0.05.

## Discussion

LiLA, like other versions of lectin arrays, fluorescent lectins and mass cytometry, offers the powerful capacity to understand the composition of glycocalyx. Differences on glycocalyx composition between normal and cancer cells as well as invading pathogens have been studied by slide-based lectin arrays^53-56^. LiLA adds a previously untapped ability to measure the composition of glycocalyx on intact cells in complex cellular mixtures, and in an animal model. Such measurements would not be possible by simply attaching “exposed” DNA to lectin proteins due to the rapid degradation of DNA barcodes in such environments. DNA degradation is prevented in LiLA as the DNA barcode is protected inside the phage. LiLA also shows that soluble diCBM40 has fundamentally different glycan recognition from near van-der-Walls (vdW)-packed multivalent displays of the same lectin and they, in turn, are different from non-linear recognition by mid-multivalent constructs. These observations are missed entirely in studies that employ soluble lectins tagged by DNA or fluorophores and they represent a yet-unexplored dimension in regulatory mechanisms in biology. Mass-spectrometry (MS) is a golden standard for analysis of structures and copy number of 100’s of diverse glycan epitopes on the surface of cells^57^. In turn, the non-destructive nature of LiLA measurements will make it possible to test how analysis inferred from MS of lysed cell populations and frozen and stained tissue sections translates to recognition of copy number and structure of those glycans on intact cells inside live organisms under the influence of their unique microenvironment.

Exploiting differences in the composition of cell surfaces is the foundation of modern breakthrough therapies: fourteen FDA-approved antibody-drug conjugates^1^, six CAR T-cells^58^, and two approved peptide-based targeted radiopharmaceuticals (Lutathera and Pluvicto)^59^ - all these examples exploit an exclusive expression of a unique, often protein-based, antigen on cancer cells. Glycan-protein interactions enable liver-specific delivery of nucleic acid therapies (GIVLAARI and 45 modalities in Phase 1-3 clinical trials^60^) and, again, they build on the unique overexpression of the GalNAc-binding receptor in liver cells. However, it is not trivial to identify a single molecular target uniquely expressed only on cancer cells and not expressed anywhere else in normal tissues in genetically diverse populations. For many pathologies, it might not be possible to identify such an exclusive target. Many cellular receptors exhibit high vs. low density on cancer vs. normal cells and residual low density of receptors poses a liability for traditional targeting and leads to toxicity. Our report focuses on sialic acid epitopes in cellular glycocalyx that are ubiquitously present on all cell types. Nevertheless, immune cells have evolved the ability to recognize the subtle differences in sialo-epitopes on cells^61^. LiLA technology demonstrates that precision in control of valency of sialic acid binding proteins offered a sharp discrimination between cells that have subtle differences in density of sialo-epitopes in their glycocalyx. A non-linear response evoked by multivalent φ-CBM_73_ (Fig. 4b) is a sharp contrast to a traditional linear response to epitope density exhibited by traditional targeting agents. φ-CBM_73_ shows a remarkable non-linear amplification of small differences in composition to nearly digital YES/NO binding. In contrast, near-vdW packed φ-CBM_290_ exhibit a traditional “linear” conversion of incremental differences in composition to incremental differences in binding to cells (Fig. 4a). The fundamental importance of cooperative ON-OFF genetic switches produced by the binding of oligomeric transcription factors to DNA have been studied since 1970’s^62^. Our data suggests that when the density of sialic acid binding receptors is turned to a specific level (e.g. 70-140 copies per 700×5-nm cylindrical scaffold, or 4.5-9.0×10^-3^ receptors/nm²), cooperativity in sialic acid recognition in cellular glycocalyx also produces ON/OFF switch in response to the density of sialic acid. This observation is particularly important because the regulation of sialic acid density is one of the fundamental hallmarks of cancer cells.

Multivalency today is often employed as a mechanism to convert the weak affinity of monovalent ligands to the strong avidity of polyvalent analogs^63^. Our manuscript investigates a lesser-explored fundamental property of multivalent scaffolds: they amplify, non-linearly, small differences in epitope density on cell surfaces and offers a different paradigm for targeting. Unlike traditional targeting agents, multivalent agents of specific valency can non-linearly diminish the liabilities associated with low expression of the same epitope on “normal” cell types. Conceptually similar non-linearity and cooperativity of several binding events govern genetic ON/OFF switches and other examples in biology that convert subtle change to a sharp response. φ-CBM_73_ construct binds to tumorigenic hypersialylated U937 and MV4-11 cells and even more significantly to CSC populations, but not healthy cells. Recurrent observations that a specific multivalent presentation of lectin can optimally detect changes in CSC/NSCC populations may have fundamental importance in CSC biology and practical outcome for identification and targeting or monitoring of CSC isolated population in heterogeneous tumors using tuned multivalent constructs (e.g. polymers or liposomes displaying a defined spacing of receptors). φ-CBM_73_ does not recognize sialic acids on the cell surface of B/T/NK cells in mice or humans, because these cells have high levels of α2-6-linked sialosides and the density of α2-3 is below a critical threshold. It is important to stress that sialic acid density on PBMCs is far from zero; soluble diCBM40 or high valency φ-CBM_290_ readily binds to sialic acids on PBMCs. Such binding is a classical demonstration of weak, “off-target liability” of low-density epitopes; however, medium-valency φ-CBM_73_ avoids this liability.

Mobile receptors can reorganize into clusters on cell surface. The static, immobile display in LiLA is different from cell surface receptors because it makes it possible to produce and test series of constructs with a well-defined spacing between the lectins starting from chemically-modified phages. LiLA platform technology displaying a defined, DNA-encoded protein densities ranging across several orders of magnitude is truly enabled by chemical modification of phage: we attempted biosynthetic display of the Sc protein on phage following best-in-class recipes for “classical” hybrid pVIII display of proteins^64^, but all cloning attempts in our hands yielded fewer than ten copies per phage (Supplementary Fig. 9). In monodisperse M13 virion, 700×5 nm rod, there is a precise 7 nm distance between each of the 2700 pVIII proteins; a stochastic ligation of proteins to a subset of pVIII proteins yields a distribution of distances, but at 300 lectins per phage, the packing of lectins on phage approaches the Van der Waals limit with more uniform spacing between proteins (Extended Data Fig. 1). Similar packing density may occur in tightly packed clusters of lectins in the lipid rafts. Precise determination of not only copy numbers and overall structure of and spacing of multivalent lectin-phage constructs is an open problem that can be tackled in the future with the help of advanced structural biology instrumentation (e.g., cryo-EM^65, 66^, TEM^67^, and AFM^68^).

Supply of lectins is central for manufacturing of lectin arrays. Both slide- and DNA-encoded lectin arrays are mostly built on plant-isolated lectins. Recombinant production of lectins with SpyTag offers convenience and control over immobilization but it requires individual expression of every lectin. Future versions of LiLA can build on other immobilization technologies to expand the scope; for example, we are exploring manufacturing of LiLA from sortase-tagged GBPs and biotinylated lectins.

Coupling of glycocalyx analysis and NGS opens new dimensions in biology. For example, Kiessling and co-workers elegantly coupled NGS to FACS of the microbiome using fluorescently labeled lectins to unveil previously unknown population dynamics in microbial communities^69^. Our manuscript shows multiple examples of LiLA-FACS pipeline analyzing the glycocalyx of live cells in complex environments, in heterogeneous cell populations, and *in vivo*. Tateno and coworkers demonstrated an exciting possibility of combining single cell RNAseq to cell staining with DNA-tagged lectins^25^. The universality of DNA-NGS technologies makes such integration possible in LiLA as well and we are currently investigating the M13 vectors with built-in DNA adapters for integration to RNAseq pipeline. The protected DNA message of LiLA will allow it to access applications not possible for simple lectins tagged with exposed DNA. The multivalent cooperative responses evoked by LiLA critically separates it from all prior technologies that employ soluble lectins. The ability of LiLA to unveil previously unexplored phenomena regulated by changes in multivalent lectin architectures will make it an important upgrade to previous platform technologies that bridge NGS, genomics and analysis of cellular glycocalyx.

## Methods

### Materials and general information

Chemical reagents were purchased from Sigma–Aldrich and Thermo Fisher Scientific unless noted otherwise. NanoDrop^TM^ (Thermo Fisher) was used to measure the absorbance of protein and DNA solutions. Sanger sequencing and next-generation sequencing was performed at the Molecular Biology Service Unit (University of Alberta) using an Illumina NextSeq500 system. All DNA primers were ordered from Integrated DNA Technologies. The U937 cells were obtained as described by Jung et al^43^. The MV4-11, SUPM2 and UCONN L2 cells were obtained as described by Li et al^46^. Biochemical reagents were purchased from Thermo Fisher Scientific unless noted otherwise. HEPES buffer contains 20 mM HEPES, 150 mM NaCl, 2 mM CaCl_2_, pH 7.4. PBS buffer contains 137 mM NaCl, 10 mM Na_2_HPO_4_, 2.7 mM KCl, pH 7.4.

### SDB clone isolation and amplification

The M13-SDB-SVEKY library described in a previous report^38^ was used to isolate the individual phage clones with built-in silent double barcodes. This is a short, readily PCR amplified region of the phage genome that varies between clones and codes for the same amino acid sequence.

### Analysis of phage modification by MALDI-TOF mass spectrometry

Sinapinic acid matrix was formed by deposition of two layers: layer 1 (10 mg/mL sinapinic acid in acetone:methanol, 4:1) and layer 2 (10 mg/mL sinapinic acid in acetonitrile:water, 1:1, with 0.1% TFA). In a typical sample preparation, 2 μL of phage solution in PBS was combined with 4 μL layer 2, and then a 1:2 mixture of layer 1:layer 2 with phage was deposited onto the MALDI inlet plate, ensuring that layer 1 was completely dry before adding layer 2 with phage. Spots were washed with 10 μL water containing 0.1% TFA. MS-MALDI-TOF spectra were recorded on the AB Sciex Voyager Elite MALDI mass spectrometer equipped with a MALDI-TOF pulsed nitrogen laser (337 nm) (3 ns pulse up to 300 μJ per pulse) operating in full-scan MS in positive ionization mode. To estimate the ratio of modified to unmodified pVIII, we implemented an automated pipeline for processing of raw MALDI *.txt files to images and integration data. This task was performed by plotMALDI.m MatLab script. The data of MALDI plots used in this work is available in Supplementary Data.

### Construction of S49C SpyCatcher variant by site-directed mutagenesis

SpyCatcher003_pDEST (Addgene, #133447) vector was mutated using primers P1 and P2 (Supplementary Table 2) to produce S49C_SpyCatcher vector. Site-directed mutagenesis was performed using QuikChange Lightning Site-Directed Mutagenesis Kit (Agilent Technologies). 25 ng of dsDNA template vector was mixed with 0.5 µL dNTP mix, 62.5 ng of each primer, 0.5 μL QuikChange Lightning Enzyme, 0.75 μL QuickSolution reagent in 1X reaction buffer in a total volume of 25 µL. The temperature cycling protocol was performed as follows: (1) 95 °C for 2 min, (2) 95 °C for 20 s, (3) 60 °C for 10 s, (4) 68 °C for 2.5 min, (5) repeat 2-4 for 18 cycles and 6) 68 °C for 5 min. PCR-amplified vector was treated with restriction enzyme *DpnI* to digest the parental dsDNA template vector. Vector was transformed into NEB® Stable chemically competent *E. coli* cells for amplification. Cells were rapidly resuspended in 1 mL of LB medium and incubated in a shaker for 60 min at 250 rpm at 30 °C. Cells were plated on carbenicillin plate (50 µg/mL) and then incubated overnight at 37 °C. Individual bacterial colonies formed in the following day were inoculated in LB medium and DNA was miniprepped and submitted for Sanger sequencing.

### Construction of phages that transduce the SpyCatcher domain

We produced filamentous phage vectors that contain the gene for the SpyCatcher003 variant cloned upstream the recombinant pVIII chain. SpyCatcher gene fragments were PCR amplified using primers P12 and P13 to produce fd-Sc1 vector (SC003_SfiI fragment) or using primers P14 and P15 to produce fd-Sc2 vector (SC003_BsaI fragment) (Supplementary Fig. 9). p88-SDB phage genome was PCR amplified using primers P16 and P17 to produce fd-Sc2 vector (p88_BsaI fragment) (Supplementary Fig. 9). PCR was performed using dsDNA template vector with 200 μM dNTPs, 0.5 μM each primer, 0.25 μL Platinum™ SuperFi™ II DNA polymerase in 1X SuperFi™ II buffer in a total volume of 25 μL. The temperature cycling protocol was performed as follows: (1) 98 °C for 30 s, (2) 98 °C for 10 s, (3) 60 °C for 10 s (for the SpyCatcher003 gene) or 55 °C for 10 s (for the p88-SDB vector), (4) 72 °C for 15 s (for the SpyCatcher003 gene) or 4 min and 15 s (for the p88-SDB vector), (5) repeat 2-4 for 30 cycles and 6) 72 °C for 5 min.

PCR amplified fragments were purified using Illustra^TM^ GFX PCR DNA and Gel Band purification kit (Cytiva). SC003_BsaI and p88_BsaI fragments were treated with restriction enzyme *BsaI*-HF®v2 (NEB #R3733S) while SC003_SfiI fragment and dsDNA p88-SDB vector were treated with restriction enzyme *SfiI* (NEB #R0123S) and then gel purified. DNA ligation was then carried out and resulting DNA was transformed into *E. coli* DH5α electrocompetent cells for amplification. Cells were rapidly resuspended in 1 mL of LB medium and incubated in a shaker for 60 min at 200 rpm at 37 °C. Cells were plated on LB-tetracycline plate (10 µg/mL) and then incubated overnight at 37 °C. Individual bacterial colonies formed in the following day were inoculated in LB medium and DNA was miniprepped and later submitted for sequencing. Purified dsDNA vectors (pVIII-Sc-1 and pVIII-Sc-2) (Supplementary Fig. 9) were transformed into One Shot™ TOP10F’ chemically competent cells for phage propagation. Cells were rapidly resuspended in 1 mL of LB medium and incubated in a shaker for 45 min at 200 rpm at 37 °C. 25 µL of transformed cells were mixed with 25 mL of 100-fold diluted overnight *E. coli* DH5α (F’) cells for phage amplification.

### Cloning of lectin constructs

SpyTag-MBP vector was purchased from Addgene (plasmid # 133450). SpyTag-diCBM40 vector was produced by restriction cloning where SpyTag was fused to the N-terminus of diCBM40. SpyTag gene was amplified from SpyTag003-MBP vectors using primers P3 and P4 (Supplementary Table 2) and cloned into diCBM40_pET-45b (+) vector by restriction enzyme digest to generate the SpyTag-diCBM40 vector. Vector was transformed into E. coli 10G electrocompetent cells for amplification.

The protocol for cloning of SpyTag-Siglec-7 vector was adapted from a published protocol^44^. To add a SpyTag to our previously developed Siglec-Fc construct, two successive PCR reactions were performed. The first PCR reaction was performed with primers P5 and P6 using, Fc-pcDNA5 from our previous work as a template. The PCR product size was validated by electrophoretic pattern in a 1% agarose gel and then the product was purified using a GeneJET Gel Extraction Kit. The second PCR reaction used the first PCR product as a template and used primers P6 and P7 to add the remaining portion of the SpyTag to the Fc. These primers also added a 5’ *AgeI* and 3’ *XmaI* restriction sites to the product of the first PCR reaction. The second PCR product was isolated as described above and was digested with *XmaI* and *AgeI* and purified using a GeneJET PCR Purification Kit. pcDNA5 was digested with *XmaI* and *AgeI* and purified as described above. The digested PCR product was ligated into the digested pcDNA5 vector. The ligation product was then transformed into chemically competent *E. coli* DH5α and miniprepped to yield pcDNA5 with SpyTag-Fc. The sequence of the construct was validated by Sanger sequencing. Amino acids 1-345 of human Siglec-7 was PCR amplified with a 5’ *NheI* and 3’ *AgeI* restriction sites and ligated into Spy-Fc-pcDNA5 vector cut with the same two restriction enzymes. The successful incorporation of the Siglec-7 into SpyTag-Fc-pcDNA5 was validated by restriction enzyme digestion and Sanger sequencing.

### Stable transfection

Chinese hamster ovary (CHO) cells were prepared for transfection by seeding at 100,000 cells/well in a 12-wells plate and cultured in growth medium (DMEM F12, 10% v/v Fetal Bovine Serum (FBS) with 100 U/ml Penicillin, and 100 μg/ml Streptomycin (Gibco)), 5% CO_2_, 37 °C. After 24 h, 3.6 µg of pOG44, 0.4 µg of Siglec-7-Spy in pcDNA5, 4.7 µL of Lipofectamine Plus Reagent (Thermo Fisher) was added to 0.5 mL of Opti-MEM (Thermo Fisher). The mixture was incubated for 15 min at room temperature. After 15 min, 15.7 µl of Lipofectamine LTX (Thermo Fisher) was added to the mixture and incubated at room temperature for 30 min. During the incubation, the growth media was removed from the seeded cells by washing with Opti-MEM. After the incubation, the mixture was added to the cells and placed into the incubator. After 24 h, 1 mL of the growth medium was added to the cells. After 48 h from the addition of the DNA mixture to the cells, the cells were expanded to a T-25 flask in growth medium with 0.5 mg/mL Hygromycin B (Thermo Fisher). The selection process continued by replacing the growth media with Hygromycin B. The Hygromycin B concentration was increased by 0.25 mg/mL every 48 h up to 1 mg/mL. The selection process was continued until non-transfected control cells were no longer viable.

### Expression and purification of S49C SpyCatcher, MBP and diCBM40

S49C_SpyCatcher, SpyTag-MBP and SpyTag-diCBM40 vectors were transformed into electrocompetent *E. coli* BL21 (DE3) cells. S49C_SpyCatcher cells were cultured overnight in 100 mL LB medium with 100 μg/mL of ampicillin, diluted to O.D 0.2 into 3 x 400 mL of LB medium with 100 μg/mL ampicillin and incubated at 37 °C at 200 rpm to OD 0.5. Expression was induced with a final concentration of 1 mM of IPTG for 3 hours at 37°C at 200 rpm. SpyTag-MBP and SpyTag-diCBM40 cells were cultured overnight in 20 mL LB medium with 50 μg/mL kanamycin and 100 μg/mL ampicillin, respectively. Cells were diluted to O.D. 0.2 into 200 mL of LB medium containing the appropriate antibiotic and incubated at 37 °C at 200 rpm to O.D. 0.5. Expression was induced with a final concentration of 1 mM of IPTG for 3 hours at 37°C at 200 rpm. Cells were harvested at 3,200 g for 10 min at 4° C. Pellet was resuspended in binding buffer (20 mM potassium phosphate, 50 mM imidazole and 500 mM NaCl, pH 7.4) and sonicated for 10 min using constant pulse ON for 15 seg and pulse OFF for 45 seg on ice. Cell lysate was harvested at 4,000 g for 20 min and supernatant was submitted to immobilized metal affinity chromatography. Sample was loaded on a 1 mL HisPur^TM^ Ni-NTA gravity-flow column and non-specific bound proteins were washed with 5 mL binding buffer. SpyTag-proteins were eluted with elution buffer (20 mM potassium phosphate, 500 mM imidazole and 500 mM NaCl, pH 7.4). The proteins were collected in 1 mL fractions and buffer exchanged to PBS buffer using a PD-10 desalting column (Cytiva).

### Expression and purification of Siglecs

Cells were expanded to ten T-175 flasks with 50 mL growth media into and placed into 37 °C incubators with 5% CO_2_. The media was harvested 7 days after confluency and filtered using a 0.22 µm filter. Purification was performed at 4 °C and initiated by equilibrating a His-Trap Excel (GE LifeSciences) column with 20 column volumes (CV) of equilibrium buffer (20 mM sodium phosphate, 0.5 M NaCl, pH 7.4). The supernatant was loaded onto the column at a rate of 1 CV/min. Non-specific protein binding was reduced by washing the column with 20 CV of washing buffer (20 mM sodium phosphate, 0.5 M NaCl, 50 mM imidazole, pH 7.4). The protein was eluted from the column using 20 CV of elution buffer (20 mM sodium phosphate, 0.5 M NaCl, 500 mM imidazole, pH 7.4), and the Siglec-7-SpyTag-TEV-Fc was collected in 1 mL fractions. The amount of protein in each fraction was measured by A_280_ via NanoDrop (Thermo Fisher). The fractions containing the protein were combined and diluted 5-fold into buffer W (100 mM Tris-HCl, 150 mM NaCl, 1 mM EDTA, pH 8). A Strep-Tactin column (IBA LifeSciences) was equilibrated with 15 CV of buffer W and the diluted protein was added onto the column at a rate of 1 CV/min. The column then was washed with 10 CV of buffer W and the protein was eluted with 10 CV of buffer E (100 mM Tris-HCl, 150 mM NaCl, 1 mM EDTA, pH 8, 5 mM Desthiobiotin). The protein was collected into 1 ml fractions. The amount of protein in each fraction was measured as described above and fractions containing protein were combined. The protein was then dialyzed against PBS overnight at 4 °C. The following day, the Siglec-7-Spy Tag-TEV-Fc was concentrated using an AMICON (Molecular Weight Cut Off 30 kDa – Merck Millipore) to approximately 0.15 mg/ml measured via NanoDrop as described above. An SDS-PAGE was used to confirm the presence and the purity of the protein.

### Generation of Siglec-7-SpyTag fragment

To remove the Fc fragment, the concentrated protein was incubated with 10 fold molar excess of TEV (Tobacco Etch Virus) protease enzyme at room temperature, overnight. The following day, the solution mixture was passed over a His-Trap Excel (GE LifeSciences) column preequilibrated with 20 column volume (CV) of equilibrium buffer (20 mM sodium phosphate, 0.5 M NaCl, pH 7.4) at RT to remove the TEV enzyme and Fc portion from the solution mixture. The flow-through containing the Siglec-7-SpyTag was collected. The amount of protein was quantified with NanoDrop, and an SDS-PAGE was used to confirm no Siglec-Spy Tag-TEV-Fc remained. The protein was dialyzed against PBS overnight at 4 °C. The following day, the protein was aliquoted into approximately 0.15 mg/mL and stored at -80 °C.

### SDS-PAGE and Densitometry Analysis

Conjugation experiments were analyzed on 10% Bis-Tris SDS-PAGE gel with MOPS electrophoretic buffer. Resolving gel was made using 10% (w/v) acrylamide, 0.27% (w/v) bis-acrylamide, 0.36 M bis-tris pH 6.6, 0.06% (w/v) APS and 0.27% (v/v) TEMED. Stacking gel was prepared with 5% (w/v) acrylamide, 0.13% (w/v) bis-acrylamide, 0.36 M bis-tris pH 6.6, 0.08% (w/v) APS and 0.41% (v/v) TEMED. For MOPS buffer, 50 mM MOPS, 50 mM Tris, 1 mM EDTA, 0.1% (w/v) SDS, and 5 mM sodium bisulfite were used. Samples were run at 35 mA. Gels were stained with 0.3% (w/v) Coomassie R-250, 10% (v/v) methanol, and 10% (v/v) acetic acid and destained with 10% (v/v) methanol / 10% (v/v) acetic acid solution.

Band intensities were calculated using the ImageJ 1.53c program (National Institutes of Health). Wand (tracing) tool was used to obtain the area under the curve of each band after plotting each gel lane. As band intensities are given as relative numbers, a gel densitometry calibration curve was not needed. The copy number was estimated by dividing the area under the curve of the product band by the area under the curve of their respective protein band at a known concentration.

### Chemical modification of phage clones with S49C SpyCatcher to build LiLA components

A solution of SDB phage (10^13^ PFU/ml in PBS) was combined with SIA (TCI America) (5 mM in DMSO) to afford a 0.1 mM concentration of SIA in the reaction mixture, which typically yields 40-50% of pVIII modification after 1h of incubation (pVIII-IA). After conjugation of SIA, phages were purified on a Zeba^TM^ Spin desalting column (7K MWCO, 0.5 mL) following the manufacturer instructions, while buffer was exchanged to 40 mM Na_2_H_2_BO_3_, pH 8.3. Solution of TCEP-reduced S49C SpyCatcher (0.8 mM in Na_2_H_2_BO_3_) was added to the solution of pVIII-IA phages (3.10^12^ PFU/ml in Na_2_H_2_BO_3_) to afford a 0.3 mM concentration of S49C SpyCatcher. The solution was further incubated overnight at room temperature. All conjugates were exposed to 5 mM L-cysteine in 100 mM sodium carbonate buffer, pH 10, to cap any unreacted iodine group. All chemical reactions were verified and quantified by MALDI-TOF MS as described in previous sections. After capping, 1/5 volume of 20% PEG 8,000/2.5 M NaCl solution was added and the solution was kept for 1 h on ice. The solution was then centrifuged at 16,000 g for 10 min at RT and the pellet was resuspended in PBS buffer.

### Conjugation of lectin to phage using SpyTag/SpyCatcher system

All conjugations were performed at room temperature for 1 hour. 0.5 – 1.10^12^ PFU/mL solution of phage and low micromolar concentrations of lectin were mixed in PBS buffer. Protein and phage concentrations were estimated spectrophotometrically at 280 nm and 269/320 nm, respectively.

### Preparation of the LiLA from lectin-phage clones

In a typical protocol, the LiLA was prepared by mixing 10^8^ PFU of desired lectin-phage conjugate in a single tube. The mixture was characterized by deep sequencing and N×10^7^ PFU (N = lectin-phage conjugates) was used for a typical cell-binding experiment. Each unique LiLA mixture was assigned a two-letter identifier (e.g., “GA”) and a “dictionary” (e.g., GA.xlsx description of the correspondence between the DNA barcodes and the lectins in the LiLA mixture is available in Supplementary Table 1). These dictionaries were used to translate from DNA in the deep-sequencing files to the corresponding lectin and their density. The composition of each naive library was used as the normalization factor for each experiment. Dictionaries for LiLA mixtures relevant to this manuscript are available in the Supplementary Data. Naïve compositions of these mixtures and references to the deep-sequencing data are available on https://48hd.cloud/ with dataset name listed in Supplementary Table 1. To access the data, enter the dataset name at https://48hd.cloud home page search bar. Locations of “silent barcode” regions SB1 and SB2 in the M13 genome are illustrated in a previously published work^38^.

### Binding of LiLA components to glycan on the cell surface

Cells were aliquoted into FACS tubes (Corning, 352054), which afforded 1 million cells per FACS tube for replicate samples. Cells were centrifuged at 300 *g* for 5 min and cell pellets were resuspended in incubation buffer (1% BSA in HEPES buffer) at 10^7^ cells per ml. Thereafter, phage was added to each FACS tube at 10^7^ PFU and incubated on ice for 1 h. After incubation, the cells were gently vortexed, and wash buffer (3 mL, 0.1% BSA in HEPES buffer) was added to each FACS tube using a serological pipette. The solution was centrifuged at 300 *g* for 5 min at 4 °C in a swinging bucket rotor. The supernatant was decanted by inverting the FACS tubes and blotting on a Kimwipe. Two additional washes were performed: during each wash, the tube was filled with 3 mL of wash buffer, gently vortexed, centrifuged, and inverted to discard the supernatant in the same manner as described above. After the last wash, the pellet was resuspended in 100 μL of wash buffer. An aliquot of this solution (20 μL) was sampled and combined with LB medium for titering. The remaining solution was subjected to proteinase treatment and used as the template for PCR reaction following the protocol described the in section “*Extraction of phage DNA from cell pellet*”.

### Treatment of cells with neuraminidase (NanH)

10^6^ cells were washed with 3 mL of cold PBS and resuspended in 50 µL solution containing 100 µg/mL neuraminidase from *Clostridium perfringens* (Roche, #11585886001). Cells were incubated for 1 h at 37 °C and washed twice with PBS before incubation with LiLA.

### Extraction of phage DNA from cell pellet

Cell pellet from panning experiment containing ∼10^6^ cells was resuspended with 50 μL 0.1 mg/mL RNase A in 1x Phusion® HF reaction buffer (NEB, Cat. B0518S). Cells were vortexed at maximum speed for 1 min. The suspension was centrifuged at 500 *g* for 3 min and the supernatant was transferred to a clean Eppendorf ^TM^ tube. 1 μL of 1% Proteinase K solution was added and the solution was incubated at 56 °C for 10 min. Proteinase K was inactivated after incubation at 97 °C for 3 min. Solution was centrifuged at 16,000 *g* for 1 min and the supernatant was used as a template for qPCR.

### Isolation of immune cells from peripheral human blood

Human blood sample collection was approved by the human research ethics board (HREB) biomedical panel at the University of Alberta. Human blood was diluted 1:10 with RBC lysis buffer (150 mM NH_4_Cl, 10 mM NaHCO_3_, 1 mM EDTA disodium salt) and incubated at RT for 7 min. Cells were harvested at 300 *g* for 5 min at 4 °C. Lysis procedure was repeated twice. The final pellet containing white blood cells was resuspended in FACS buffer (1X PBS, 2% FBS v/v) and stained with Fc block agent (1:100 v/v) on ice for 10 min followed by staining with antibodies. The following antibody cocktail was used: hCD3 (BV650, clone OKT3, Biolegend Cat No.317324, 1:100), hCD19 (APC-Cyanine7, clone SJ25C1, Biolegend Cat No. 363010, 1:100), hCD56 (BV510, clone NCAM16.2, BD Bioscience Cat No. 563041, 1:100) and hCD14 (PE-Cyanine7, clone 61D3, Invitrogen Cat. No. 25-0149-41, 1:100). Cells were washed twice with FACS buffer and resuspended in FACS buffer containing propidium iodide (PI) (1:100 v/v). Fluorescence-activated cell sorting (FACS) was performed on a Sony MA900 cell sorter (Faculty of Medicine & Dentistry Flow Cytometry Facility, University of Alberta) and sorted cells were collected in FACS tubes containing 500 µL of 1:1 PBS:FBS. LiLA was added at 10^7^ PFU either together with antibodies before sorting (LiLA+FACS) or after FACS (FACS→LILA).

### Flow cytometry

Flow cytometry measurements were taken on a Accuri^TM^ C6 Flow Cytometer (BD Biosciences) for all experiments except for experiment with soluble Siglec-7-Fc constructs in which a 5-laser Fortessa X-20 (BD Biosciences) was used instead.

### Panning of LiLA in mice

All the procedures and experiments with mice were approved (AUP00002467) by the UAlberta Animal Care and Use Committees (ACUC). The protocol was approved as per the Canadian Council on Animal Care (CCAC) guidelines. All mice were maintained in pathogen-free conditions at the University of Alberta breeding facility. Animals were injected with LiLA in the tail vein (0.2 mL, 3 x 10^7^ PFU/mL in PBS). We used 10-12 weeks old male C57BL/6J mice for this study. One-hour post-injection mice were bled (0.1 mL) and immediately afterwards they were euthanized with CO_2_. Internal organs (heart, liver, kidney, lungs, and spleen) were collected and stored in cold PBS. Tissues were homogenized using a 45 μm cell strainer and washed with 5 mL cold PBS. Homogenized tissues of each organ were centrifuged at 300 *g* for 5 min and the pellet was submitted to proteinase K treatment following the protocol described in the section “*Extraction of phage DNA from cell pellet*”. 100-fold diluted plasma samples and phage DNA isolated from homogenized tissues were amplified by PCR using the protocol described in the “*PCR protocol section*”.

### Isolation of immune cells from murine spleen

Mice **s**pleen was cut into small pieces (∼2 mm^2^), homogenized using a 45 μm cell strainer, and washed with 5 mL cold PBS. The homogenized spleen was centrifuged at 300 *g* for 5 min and resuspended in 10 mL cold RBC lysis buffer for 5 min. 20 mL cold PBS was added to stop the lysis process and cells were harvested at 300 *g* for 5 min. The pellet was washed with FACS buffer and stained with an antibody cocktail on ice for 30 min. The following antibody cocktail was used: mCD3 (FITC, clone 17A2, Invitrogen Cat No.11-0032-80, 1:100) and mCD19 (APC, clone eBio1D3, Invitrogen Cat No. 17-0193-80, 1:100). Cells were sorted as described in *“Isolation of immune cells from peripheral human blood*”. LiLA was either injected into mice as described in “*Panning of LiLA in mice*” (pre-harvesting incubation) or added after FACS (post-sorting incubation).

### PCR protocol and Illumina sequencing

2 μl volume of DNA template solution after the binding procedure was amplified in a total volume of 20 μl with 1× qPCR Master Mix (MBSU, University of Alberta), 0.5 μM forward primer (P8) and 0.5 μM reverse primer (P9). For amplifying clonal phage and naive libraries, the template (phage solution) was diluted 100-fold in water. The reaction was run in Bio-Rad C1000 Thermal cycler CFX96 Real Time System using the following thermocycler settings: (1) 95 °C for 3 min, (2) 95 °C for 10 s, (3) 60 °C for 20 s, (4) 72 °C for 10 s (with picture), (5) repeat steps 2– 4 for 39 cycles, and (6) melting point step 65-95°C (with picture). The product from the qPCR step was used in the next PCR amplification step to prepare samples for Illumina sequencing (Supplementary Fig. 10). The protocol was adapted from Sojitra et al^38^ with minor modifications. 1 μl volume of DNA template from first-step PCR was amplified in a total volume of 25 μl with 1× Phusion buffer, 200 μM of each dNTP, 0.4 μM forward barcoded primer (P10), 0.4 μM reverse barcoded primer (P11), 5% (v/v) DMSO and 0.5 U Phusion High-Fidelity DNA polymerase (NEB, M0530S). Cycling was performed using the following thermocycler settings: (1) 95 °C for 3 min, (2) 95 °C for 10 s, (3) 58 °C for 30 s, (4) 72 °C for 20 s, (5) repeat steps 2–4 for ten cycles and (6) 72 °C for 5 min.

PCR products were quantified with a 2% (w/v) agarose gel in Tris–borate–EDTA buffer at 100 V for ∼35 min using a 1 Kb Plus DNA ladder. PCR products that contained different indexing barcodes were pooled, allowing 40 ng of each product in the mixture. The mixture was purified using Nucleospin® Gel and PCR clean-up kit (Macherey-Nagel # 740609), quantified by Qubit (Thermo Fisher), and sequenced using the Illumina NextSeq paired-end 500/550 High Output kit version 2.5 (2 × 75 cycles). Data were automatically uploaded to the BaseSpace Sequence Hub. The processing of data is described in the section “*Processing of Illumina data*”.

### General data processing methods

In NGS sequencing, the reads with length above or below 96 base pairs, were systematically excluded from all analyses. NGS Reads that could not be mapped to specific DNA-barcodes or their neighbors (Hamming distance = 1) were excluded from all of the the analyses. Data analysis was performed in Python and testing differences for significance in LiLA data was performed as Mann-Whitney U test analyses of phage-displayed libraries based on fold changes (FC) enrichment analysis. The FC-analysis assumed that the group of φ-MBP_140_ phages does not bind to test samples. All other lectin FC ratios were normalized to MBP. Core processing scripts are available as part of the Supplementary information or on GitHub. To assess the significance of a lectin binding in a specific experiment the FC analysis of the levels of the DNA barcode associated with that lectin in “test” sets of DNA reads was compared to the levels of the same barcode in naïve library or “input” data set. Reads in the input dataset were averaged when more than one input sample was sequenced. Means of input were not used to calculate the statistics of the data. Prior to FC-analysis, “test” and “input” data sets were retrieved from the http://48hd.cloud/ server as tables of DNA sequences, and raw sequencing counts (Supplementary Table 1). DNA reads that could not be mapped to any entries in the LiLA dictionary were discarded.

### Processing of Illumina data

The processing of FASTQ files downloaded from BaseSpace™ Sequence Hub to reads and frequencies of lectins were performed as previously described^38, 70^. A LiLA-specific lookup table was used to convert the identified SDB to lectin and display density. Translated files with raw DNA reads, raw counts, and mapped lectins were uploaded to the https://48hd.cloud/. server. All LiLA sequencing data is publicly available at this location. Each experiment has a unique alphanumeric name (e.g., 20220527-1163LILAAuaaYX-GL, see Supplementary Table 1).

## Data availability

All raw deep-sequencing data are publicly available at 48hd.cloud with data-specific URLs listed in Supplementary Table 1. Source data are provided with this paper.

## Code availability

MATLAB and Python scripts were deposited at https://github.com/derdalab/lila

## Acknowledgements

We thank the staff at the University of Alberta mass spectrometry facility (Chemistry Department) for help with MALDI analysis and S.Dang at the molecular biology service unit for assistance with Illumina sequencing. We thank Kelli A. McCord (University of Alberta) for assistance with blood drawing. Cell sorting was performed at University of Alberta, Faculty of Medicine and Dentistry Flow Cytometry Facility. We acknowledge funding from NSERC (RGPIN-2022-04484 to R.D.), CIHR (#180445 to R.D.), Glyconet (CR-29 and TP-22 to R.D.), Alberta Innovates Strategic Research Projects (to R.D.), Alberta Innovates Graduate Student Scholarship (to G.M.L.), and the Canada Excellence Research Chair Program (L.K.M.). G.M. received grant from São Paulo Research Foundation (FAPESP - grant number 2022/02456-0) and a Productivity Fellowship from the Brazilian National Counsel of Technological and Scientific Development (CNPq 306060/2022-1).

## Author Contributions

G.M.L. constructed and characterized all LiLA phages and libraries and performed statistical analysis. E.A.V. and R.H. isolated phage clones. S.S. performed animal experiments. E.J.C. performed statistical analysis and computational modeling of proteins on phage. G.M.L., C.Y.Y., and M.S. performed cell-based assays. E.S. performed flow cytometry with Siglec-Fc constructs. Z.J.C. cloned and expressed Siglec-Fc constructs. Z.J.C. and J.L. contributed mammalian cell lines. D.Y. and C.P. synthesized TAMRA-SpyTag peptide. A.A. and L.K.M. contributed plasmids. G.M.L and R.D. wrote the manuscript. R.D., G.M., R.L., L.K.M., and M.S.M edited the final manuscript and contributed intellectual and strategic input. All authors approved the final manuscript.

## Extended Data Figures

**Extended Data Fig. 1.**
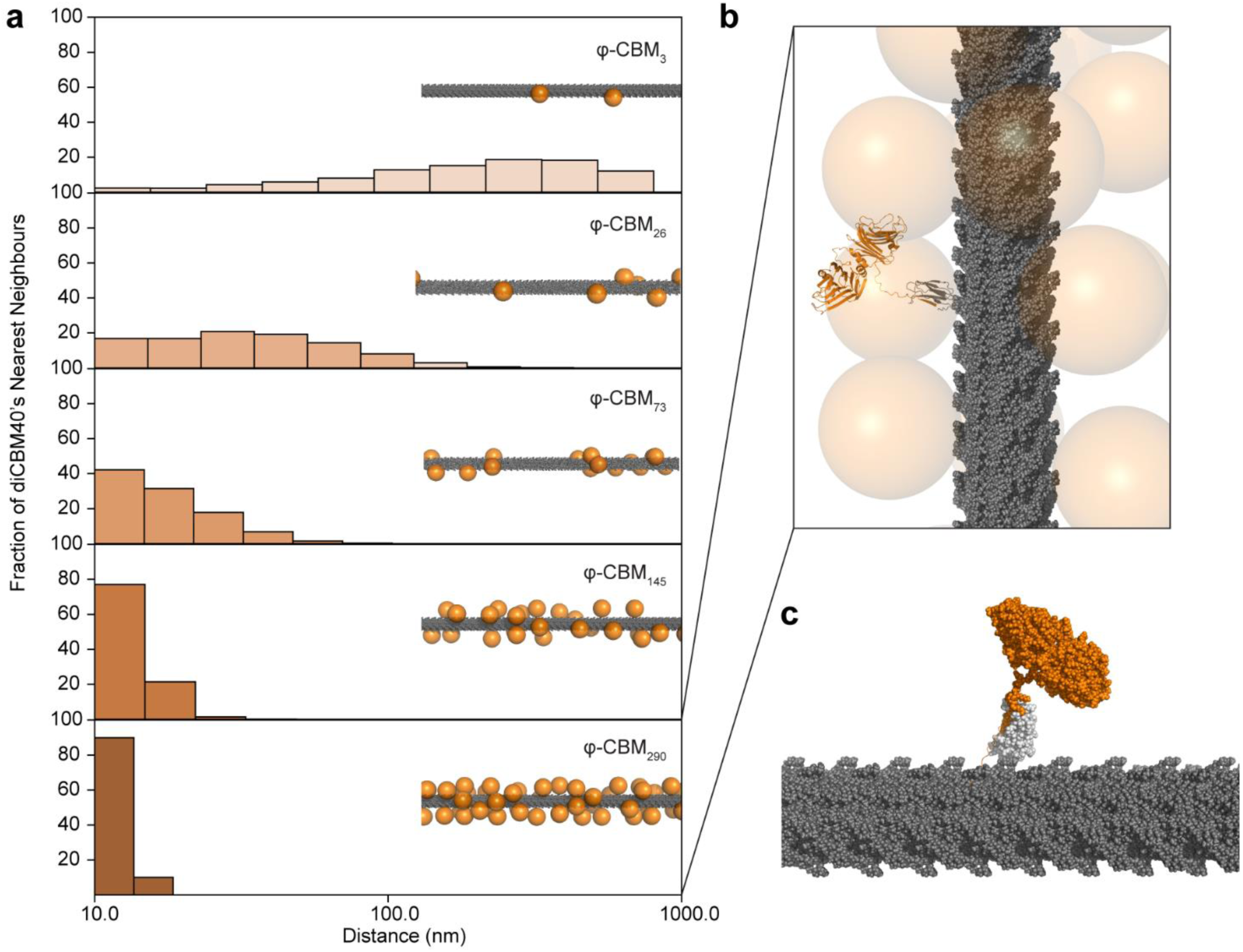
Nearest-neighbour spacing between diCBM40 at different modification densities. **a,** Histograms of the nearest-neighbour spacing at different densities. Orange spheres on phage diagram denote diCBM40-modified N-termini of pVIII proteins. diCBM40 distributions were modeled using the N-termini of the pVIII viral capsid protein based on the capsid geometry found in Model 1 of PDB entry 2MJZ^71^. **b,** Front view of the viral capsid structure at diCBM40 density equivalent to 290 copies per virion showing a cartoon representation of diCBM40 fused with SpyCatcher. **c,** Model of diCBM40 fused with SpyCatcher on phage using a sphere representation.

**Extended Data Fig. 2.**
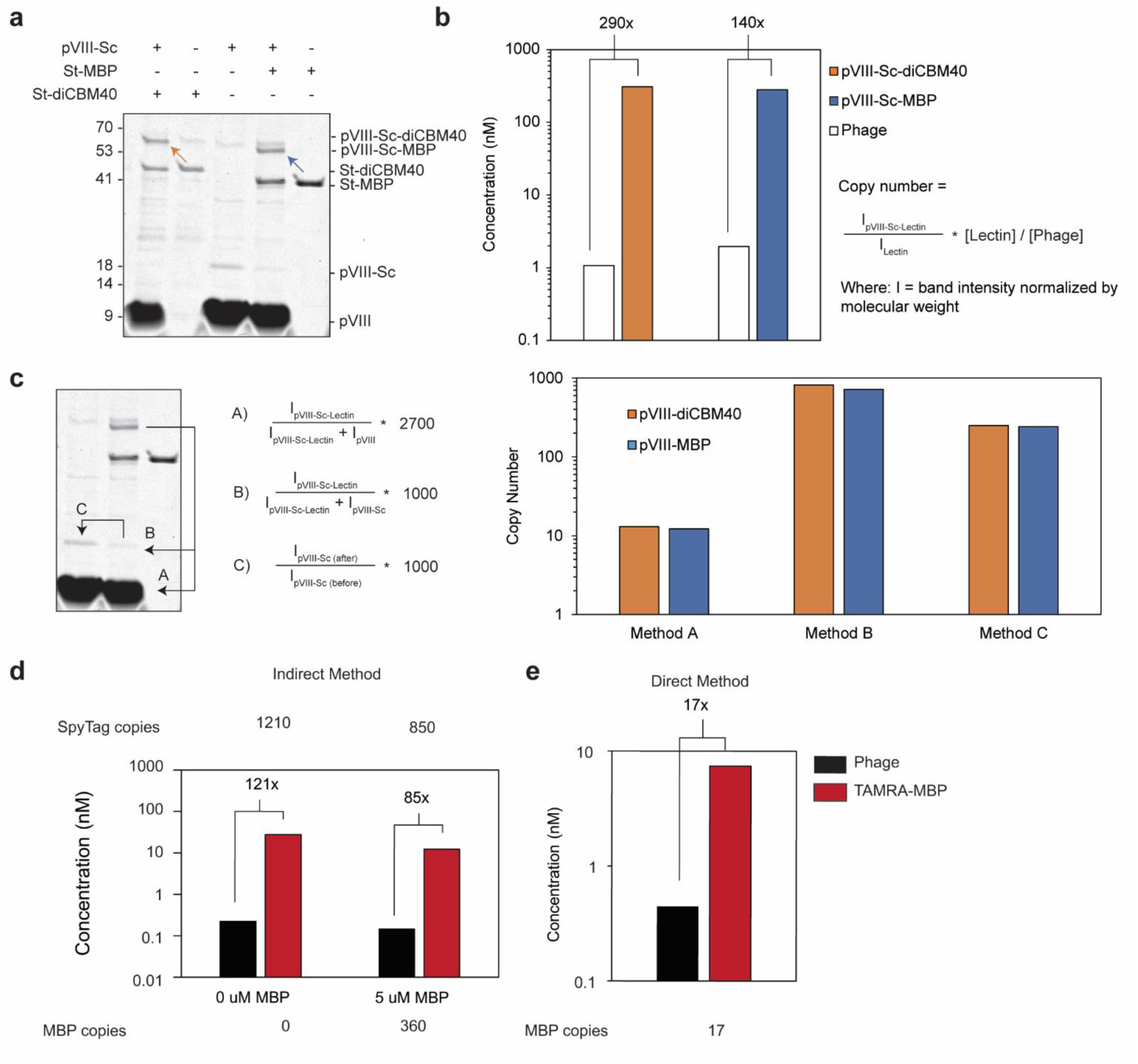
Copy number estimation of proteins on phage by various methods. **a,** SDS-PAGE of φ-Sc_1000_ phages conjugated to St-diCBM40 and St-MBP. Orange and blue arrows indicate pVIII-Sc protein fused with St-diCBM40 and St-MBP, respectively. **b,** Concentration of phages and pVIII-Sc-lectins (pVIII-Sc-diCBM40 and pVIII-Sc-MBP) as measured by qPCR and SDS-PAGE densitometry, respectively. The copy number is estimated by dividing the intensity of the pVIII-Sc-lectin band (product) by the intensity of the initial lectin band (starting material). Multiplying this ratio by the concentration of starting material results in the concentration of the product. Dividing the molar concentration of the product by the molar concentration of phage gives the copy number. **c,** Alternative copy number estimations consider the intensity of the pVIII and pVIII-Sc bands. 2700 is the average number of pVIII proteins on a phage virion; 1000 is the estimated number of pVIII-Sc proteins as measured by MALDI. **d,** We compared the fluorescence of phages conjugated to TAMRA-SpyTag before and after reaction with St-MBP. The difference in fluorescence results in the copy number of MBP molecules per phage (indirect method). Phages reacted with a solution containing 9:1 SpyTag: TAMRA-SpyTag. **e,** Reacting fluorescently labeled MBP gives a direct method for quantifying the copy number of MBP per phage. For all measurements, *n* = 1 experiment.

**Extended Data Fig.3.**
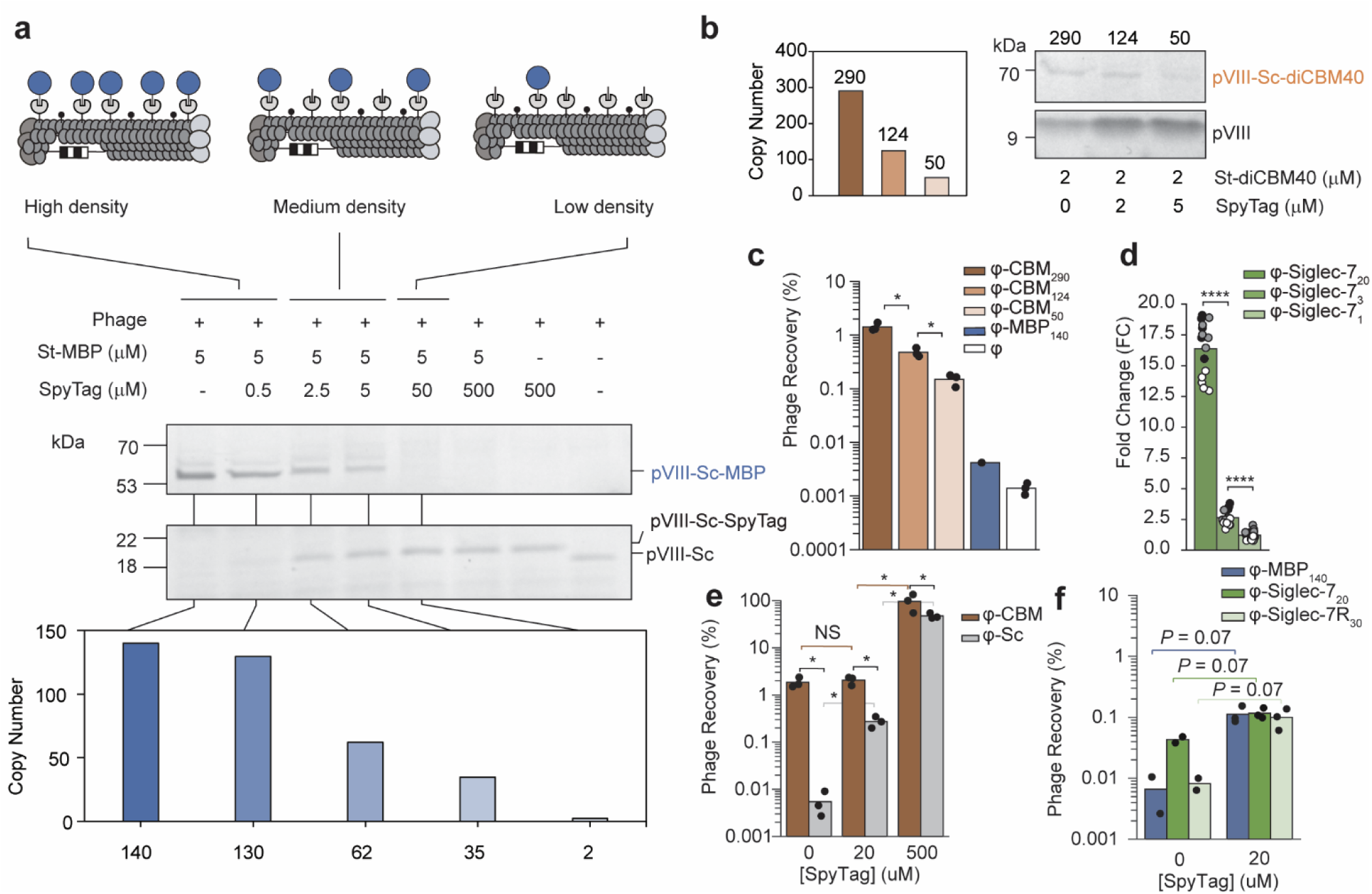
Density control of φ-lectins using SpyTag peptide. **a,** SDS-PAGE image of synthesis of multiple densities of φ-MBP phages and relative copy number estimation based on the highest φ-MBP density. This observation allowed us to extrapolate that copy number of proteins on phage scales with relative concentration of SpyTag-containing constructs in solution. **b,** Quantification of copy number of diCBM40 displayed on phage by densitometry relative to the highest diCBM40 density. *n* = 1 experiment. **c,** Binding of high, mid and low diCBM40 densities on WT U937 cells as measured by phage titering. φ-MBP_140_ and blank φ phages were used as controls. Valu es represent mean of *n* = 3 for all constructs except φ-MBP_140_ (*n* = 1), Mann-Whitney U Test**. d,** Binding of multiple densities of φ-Siglec-7 phages on WT U937 cells as measured by deep sequencing. Values represent mean of *n* = 3 (technical replicates) x 5 (DNA barcodes per construct), Mann-Whitney U Test. **e,** Effect of various SpyTag concentrations on the binding of φ-CBM and φ-Sc phages on WT U937 cells. Values represent mean of *n* = 3, Mann-Whitney U Test**. f,** Effect of SpyTag concentration on binding of φ-lectins on WT U937 cells. Values represent mean of *n* = 2 (0 µM SpyTag) and *n* = 3 (20 µM SpyTag). In **e-f,** all respective *n* values are technical replicates. NS, *P* > 0.05; * *P* < 0.05; **** *P* < 0.0001.

**Extended Data Fig. 4.**
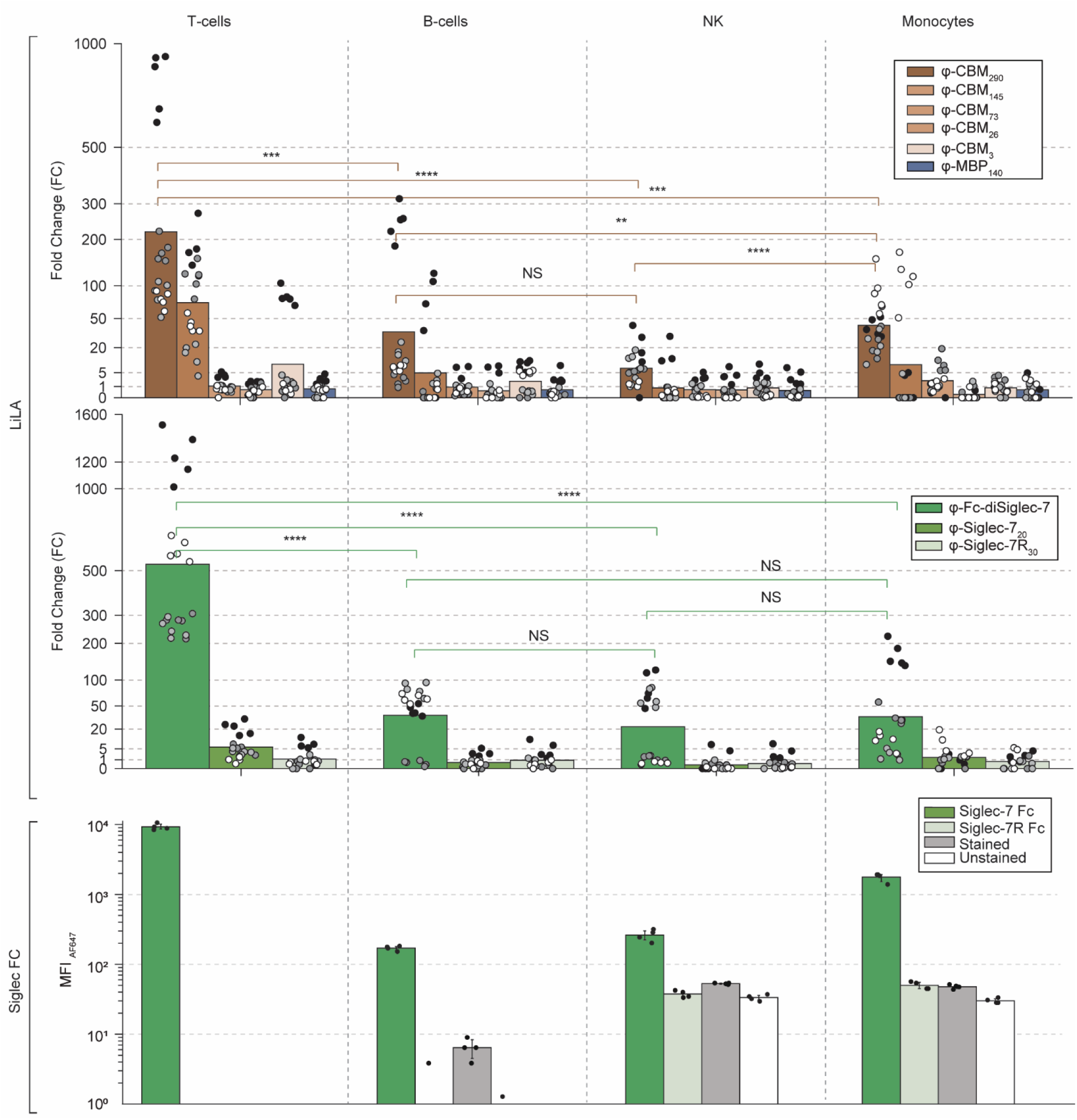
Comparison of binding of φ-CBM and φ-Siglec-7 phages to PBMCs by LiLA obtained using the FACS+LiLA approach and measured by deep sequencing vs Siglec-Fc constructs using flow cytometry. FC of φ-MBP_140_ clones was set to one. Values represent mean of *n* = 4 (technical replicates) x 5 (DNA barcodes per construct) for LiLA measurements, and *n* = 4 technical experiments for flow cytometry measurements. NS, *P* > 0.05; ** *P* < 0.01; *** *P* < 0.001; **** *P* < 0.0001.

