## Supplementary material for "Multivalent DNA-encoded lectins on phage enable detecting compositional glycocalyx differences": GL_IV_151_cbm_validation.py

```
import pandas as pd
import matplotlib.pyplot as plt
import os
import numpy as np
import seaborn as sns

plt.rcParams.update({'font.size': 7, 'font.family': 'arial', 'pdf.fonttype': 42})
pd.set_option('display.max_rows', None)

file_name = 'GL_IV_151_cbm_validation_master_list.csv'
file_path = os.path.join(os.getcwd(), file_name)
dictionary = 'GA.xlsx'
number_sdbs = pd.read_excel(os.path.join(os.getcwd(), dictionary)).shape[0]
replicates = 3
targets = 2
figure = 'Fig2a.xlsx'

### STEP 1

# 1 - Creates a master list of all SDBs and associated lectins
if not os.path.exists(file_path):

    master_list = pd.read_excel(os.path.join(os.getcwd(), dictionary)).iloc[:,[0,2,5]].set_index('SDB')

    # 2 - Opens the folder containing all sequencing files
    folder = os.path.join(os.getcwd(), 'Sequencing Files', file_name[:-16])
    files = os.listdir(folder)
    for file in files:

        # 3 - Creates panda dataframe from illumina file
        data = pd.read_csv(f'{folder}/{file}', sep=' ', header=9)
        primers = list(data.columns.values)[6:]

        # 4 - Parses through each primer combination (one replicate)
        for reads in primers:

            total_reads = int(data[reads][0])

            # 5 - Combine reads with same mindex and delete all mindex != 0
            y = 0
            for x in data['mindex']:
                if x != 0:
                    data.loc[data.index == x, reads] += data[reads][y]
                y += 1
            mindex_0 = data[data['mindex'] == 0]


            # 6 - Creates a list of desirable SDBs and corresponding reads in ppm
            df = []
            for x in master_list['Sequence']:

                for y in mindex_0['Nuc']:
                    if x.upper() == y:
                        raw = (mindex_0.loc[mindex_0['Nuc'] == y][reads].tolist()[0])
                        ppm = ((mindex_0.loc[mindex_0['Nuc'] == y][reads].tolist()[0])/total_reads)*1000000
                        sequence = master_list.loc[master_list['Sequence'] == x].index[0]

                        df.append([raw, ppm, sequence])
            df = pd.DataFrame(df, columns=[reads[:-7]+'_Raw', reads[:-7]+'_CPM', 'SDB']).set_index('SDB')

            # 7 - Export data to master list


            master_list= pd.concat([master_list, df], axis=1)

            print(f'File {file}, read {reads} completed')
            print(master_list)

    raw_cpm = [0,1] + list(range(2, master_list.shape[1], 2)) + list(range(3, master_list.shape[1], 2))
    master_list.iloc[:,raw_cpm].to_csv(file_path)


# PART 2

# 1 - Import data from step 1, fill Nan with 0 and convert to numpy array
data = pd.read_csv(file_path).fillna(0).to_numpy()

igreen_test = np.mean(data[0,15:18]).reshape(1,1)
igreen_control = np.mean(data[0,21:24]).reshape(1,1)
iblue_test = np.mean(data[1,15:18]).reshape(1,1)
iblue_control = np.mean(data[1,21:24]).reshape(1,1)


# 4 - Obtain Fold changes
green_test, green_control, blue_test, blue_control = data[0,18:21], \
                       data[0,24:], \
                       data[1,18:21], \
                       data[1,24:],

fc = np.concatenate(((green_control / igreen_control).reshape(3, 1),
                     (blue_control / iblue_control).reshape(3, 1),
                     (green_test / igreen_test).reshape(3, 1),
                     (blue_test / iblue_test).reshape(3, 1),
                     igreen_control,
                     igreen_test,
                     iblue_control,
                     iblue_test
                     ))

cells = np.array(['WT_control']*number_sdbs*replicates + ['WT_test']*number_sdbs*replicates +
                 ['Input_control']*number_sdbs + ['Input_test']*number_sdbs).\
    reshape((number_sdbs*replicates*targets + (number_sdbs*2), 1))

phage = np.array(['Green']*3 + ['Blue']*3 + ['Green']*3 + ['Blue']*3 +
                 ['Green']*2 + ['Blue']*2).reshape(16, 1)


df = pd.DataFrame(np.concatenate((cells, phage, fc), axis=1), columns=['cells', 'phage', 'fold'])
df_cells = df.loc[(df['cells'] != 'Input_control') & (df['cells'] != 'Input_test')]
df_input = df.loc[(df['cells'] == 'Input_control') | (df['cells'] == 'Input_test')]


export = pd.read_csv(file_path).fillna(0)
fc1, fc2, fc3, ix, = df_cells[::3], df_cells[1::3], df_cells[2::3], df_input

everything = [fc1, fc2, fc3, ix]
for replicate in everything:
    for cell_type in set(replicate['cells']):
        desired_column = replicate.loc[replicate['cells'] == cell_type]['fold']
        desired_column = desired_column.rename(cell_type).reset_index(drop=True)
        export = pd.concat([export,desired_column], axis=1)

export.to_excel(figure, freeze_panes=(1,4))


# 5 - Plot the graphs

sns.set(rc={'figure.figsize': (3.5, 3.5)}, style='ticks')

plot = sns.catplot(x='cells', y='fold', hue='phage',
                   kind='bar', data=df_cells,
                   palette=['#ffffff', '#c2c4c6'],
                   edgecolor='black',
                   ci='sd', errcolor='black', capsize=0.05, errwidth=1.5,
                   legend=False, aspect=6 / 9
                   )
plot = sns.stripplot(x='cells', y='fold', hue='phage',
                     jitter=0.2, dodge=True, data=df_cells,
                     color='black')

plt.yscale('log')
plt.xlabel("Phages")
plt.ylabel("Fold Change (FC)")
plt.ylim(0.001)


ticks = [0.001,0.01, 0.1, 1, 10]
labels = [0.001, 0.01, 0.1, 1, 10]
plt.yticks(ticks, labels)

plt.savefig('your_plot.pdf', transparent=True)
plt.show()
```
