## Supplementary material for "Multivalent DNA-encoded lectins on phage enable detecting compositional glycocalyx differences": GL_V_2_cell_panning_LiLA_v1_2.py

```
import pandas as pd
import os
import numpy as np
import matplotlib.pyplot as plt
import seaborn as sns
import math

plt.rcParams.update({'font.size': 7, 'font.family': 'arial', 'pdf.fonttype': 42})
pd.set_option('display.max_rows', None)

file_name = 'GL_V_2_cell_panning_LiLA_v1_2_master_list.csv'
file_path = os.path.join(os.getcwd(), file_name)
dictionary = 'GC.xlsx'
number_sdbs = pd.read_excel(os.path.join(os.getcwd(), dictionary)).shape[0]
replicates = 3
targets = 3
figure = 'Fig2c.xlsx'


# PART 1
if not os.path.exists(file_path):

    master_list = pd.read_excel(os.path.join(os.getcwd(), dictionary)).iloc[:, [0, 2, 5]].set_index('SDB')

    # 2 - Opens the folder containing all sequencing files
    folder = os.path.join(os.getcwd(), 'Sequencing Files', file_name[:-16])
    files = os.listdir(folder)
    for file in files:
        # 3 - Creates panda dataframe from illumina file
        data = pd.read_csv(f'{folder}/{file}', sep=' ', header=11)
        primers = list(data.columns.values)[6:]

        # 4 - Parses through each primer combination (one replicate)
        for reads in primers:

            total_reads = int(data[reads][0])

            # 5 - Combine reads with same mindex and delete all mindex != 0
            y = 0
            for x in data['mindex']:
                if x != 0:
                    data.loc[data.index == x, reads] += data[reads][y]
                y += 1
            mindex_0 = data[data['mindex'] == 0]

            # 6 - Creates a list of desirable SDBs and corresponding reads in ppm
            df = []
            for x in master_list['Sequence']:

                for y in mindex_0['Nuc']:
                    if x.upper() == y:
                        raw = (mindex_0.loc[mindex_0['Nuc'] == y][reads].tolist()[0])
                        ppm = ((mindex_0.loc[mindex_0['Nuc'] == y][reads].tolist()[0]) / total_reads) * 1000000
                        sequence = master_list.loc[master_list['Sequence'] == x].index[0]

                        df.append([raw, ppm, sequence])
            df = pd.DataFrame(df, columns=[reads[:-7] + '_Raw', reads[:-7] + '_CPM', 'SDB']).set_index('SDB')

            # 7 - Export data to master list

            master_list = pd.concat([master_list, df], axis=1)

            print(f'File {file}, read {reads} completed')
            print(master_list)

    # Separate column of raw reads from CPM and puts values in the correct order (i.e. input comes first followed
    # by the corresponding targets
    raw_cpm = [0, 1] + list(range(2, master_list.shape[1], 2)) + list(range(3, master_list.shape[1], 2))
    master_list.iloc[:, raw_cpm].to_csv(file_path)


# PART 2

# 1 - Import data from step 1, fill Nan with 0 and convert to numpy array
data = pd.read_csv(file_path).fillna(0).to_numpy()
print(data)
# 2 - take the mean of all mSDBs (Sc and Blank)
sc = data[4:13]
blank = data[13:]
mean_sc = np.mean(sc[:, 3:], axis=0)
mean_blank = np.mean(blank[:, 3:], axis=0)

new_sc = np.concatenate((['various_SDBs', 'various_sequences', 'Sc'], mean_sc)).reshape(1, 23)
new_blank = np.concatenate((['various_SDBs', 'various_sequences', 'blank'], mean_blank)).reshape(1, 23)
print(new_sc)

# 3 - Combine mSDB means with the rest of the library

new_data = np.concatenate((data[:4], new_sc, new_blank))
print(new_data)


# 4 - Obtain Fold changes
input, wt, cmas, chst1 = new_data[:, 13].reshape(6, 1), \
                         new_data[:, 14:17], \
                         new_data[:, 17:20], \
                         new_data[:, 20:]

fc = np.concatenate(((wt / input).reshape(18, 1),
                     (cmas / input).reshape(18, 1),
                     (chst1 / input).reshape(18, 1),
                     input.reshape(6,1)))

cells = np.array(['WT'] * 18 + ['CMAS'] * 18 + ['CHST1'] * 18 + ['Input']*6).reshape(60, 1)

lectin = np.array((['MBP'] * 3 + ['diCBM40-[290]'] * 3 + ['Siglec-7'] * 3 + ['Siglec-7R'] * 3 + ['Sc'] * 3 + ['Blank'] * 3) * 3 +
                  (['MBP'] + ['diCBM40-[290]'] + ['Siglec-7']+ ['Siglec-7R'] + ['Sc'] + ['Blank'])).reshape(
    60, 1)

df = pd.DataFrame(np.concatenate((cells, lectin, fc), axis=1), columns=['cells', 'lectin', 'recovery'])
df_cells = df.loc[(df['cells'] != 'Input')]
df_input = df.loc[(df['cells'] == 'Input')]


def mbp_norm(df_cell):
    mbp_values = df_cell.loc[df_cell['lectin'].isin(['MBP'])]
    means = [mbp_values['recovery'][0:3].mean(),
             mbp_values['recovery'][3:6].mean(),
             mbp_values['recovery'][6:9].mean()]

    mean = mbp_values['recovery'].mean()

    new_fc = []
    counter = 0
    index = 0
    for x in df_cell['recovery']:
        if counter == 18:
            counter = 0
            index +=1
            new_fc.append(x / means[index])
            counter += 1
        else:
            new_fc.append(x / means[index])
            counter += 1


    df_cell['recovery'] = new_fc

    return df_cell


test = mbp_norm(df_cells)

export = pd.read_csv(file_path).fillna(0)
fc1, fc2, fc3, ix, nfc1, nfc2, nfc3  = df_cells[::3], df_cells[1::3], df_cells[2::3], df_input, \
                                 test[::3], test[1::3], test[2::3]

everything = [fc1, fc2, fc3, ix, nfc1, nfc2, nfc3]

for replicate in everything:
    for cell_type in set(replicate['cells']):
        desired_column = replicate.loc[replicate['cells'] == cell_type]['recovery']
        desired_column = desired_column.rename(cell_type).reset_index(drop=True)
        export = pd.concat([export,desired_column], axis=1)

export.to_excel(figure, freeze_panes=(1,4))


if __name__ == '__main__':


    # 5 - Plot the graphs

    #sns.set(rc={'figure.figsize': (3.5, 3.5)}, style='ticks')
    fig, ax = plt.subplots(figsize=(3.5,1))
    plot = sns.barplot(x='cells', y='recovery', hue='lectin', order=['WT', 'CMAS', 'CHST1'],
                       data=test,
                       hue_order=['diCBM40-[290]', 'MBP', 'Siglec-7', 'Siglec-7R', 'Blank', 'Sc'],
                       palette=['#ec7e30', '#4268b1', '#71ad44', '#93c954', '#ffffff', '#b1adad'],
                       edgecolor='black',
                       ci='sd', errcolor='black', capsize=0.05, errwidth=1.5

                       )
    plot = sns.stripplot(x='cells', y='recovery', hue='lectin', order=['WT', 'CMAS', 'CHST1'],
                         jitter=0.2, dodge=True, data=test, hue_order=['diCBM40-[290]', 'MBP', 'Siglec-7', 'Siglec-7R', 'Blank', 'Sc'],
                         color='black', size=2)

    handles, labels = plot.get_legend_handles_labels()
    l = plt.legend(handles[6:12], labels[6:12])

    plt.yscale('log')
    plt.xlabel("Cell lines")
    plt.ylabel("Fold Change (FC)")
    plt.ylim(0.005)

    ticks = [0.005, 0.01, 0.1, 1, 10, 100, 1000, 10000]
    labels = [0.005,0.01, 0.1, 1, 10, 100, 1000, 10000]
    plt.yticks(ticks, labels)

    plt.savefig('your_plot.pdf', transparent=True)
    plt.show()
```
