## Supplementary material for "Multivalent DNA-encoded lectins on phage enable detecting compositional glycocalyx differences": GL_VI_148_lila_density_neuraminidase.py

```
import pandas as pd
import os
import numpy as np
import matplotlib.pyplot as plt
import seaborn as sns


plt.rcParams.update({'font.size': 7, 'font.family': 'arial', 'pdf.fonttype': 42})
pd.set_option('display.max_rows', None)

file_name = 'GL_VI_148_lila_density_neuraminidase_master_list.csv'
file_path = os.path.join(os.getcwd(), file_name)
dictionary = 'GD.xlsx'
number_sdbs = pd.read_excel(os.path.join(os.getcwd(), dictionary)).shape[0]
replicates = 3
targets = 4
figure = 'Fig2e.xlsx'


### STEP 1

# 1 - Creates a master list of all SDBs and associated lectins
if not os.path.exists(file_path):

    master_list = pd.read_excel(os.path.join(os.getcwd(), dictionary)).iloc[:,[0,2,5]].set_index('SDB')

    # 2 - Opens the folder containing all sequencing files
    folder = os.path.join(os.getcwd(), 'Sequencing Files', file_name[:-16])
    files = os.listdir(folder)
    for file in files:

        # 3 - Creates panda dataframe from illumina file
        data = pd.read_csv(f'{folder}/{file}', sep=' ', header=9)
        primers = list(data.columns.values)[6:]

        # 4 - Parses through each primer combination (one replicate)
        for reads in primers:

            total_reads = int(data[reads][0])

            # 5 - Combine reads with same mindex and delete all mindex != 0
            y = 0
            for x in data['mindex']:
                if x != 0:
                    data.loc[data.index == x, reads] += data[reads][y]
                y += 1
            mindex_0 = data[data['mindex'] == 0]


            # 6 - Creates a list of desirable SDBs and corresponding reads in ppm
            df = []
            for x in master_list['Sequence']:

                for y in mindex_0['Nuc']:
                    if x.upper() == y:
                        raw = (mindex_0.loc[mindex_0['Nuc'] == y][reads].tolist()[0])
                        ppm = ((mindex_0.loc[mindex_0['Nuc'] == y][reads].tolist()[0])/total_reads)*1000000
                        sequence = master_list.loc[master_list['Sequence'] == x].index[0]

                        df.append([raw, ppm, sequence])
            df = pd.DataFrame(df, columns=[reads[:-7]+'_Raw', reads[:-7]+'_CPM', 'SDB']).set_index('SDB')

            # 7 - Export data to master list


            master_list= pd.concat([master_list, df], axis=1)

            print(f'File {file}, read {reads} completed')
            print(master_list)

    # Separate column of raw reads from CPM and puts values in the correct order (i.e. input comes first followed
    # by the corresponding targets
    raw_cpm = [0,1] + list(range(2, master_list.shape[1], 2)) + list(range(3, master_list.shape[1], 2))
    master_list.iloc[:,raw_cpm].to_csv(file_path)

### PART 2

plt.rcParams.update({'font.size': 7, 'font.family': 'arial', 'pdf.fonttype': 42})

# 1 - Import data from step 1, fill Nan with 0 and convert to numpy array
data = pd.read_csv(file_path).fillna(0).to_numpy()

# 4 - Obtain Fold changes
input, wt, wt_n, cmas, cmas_n = data[:, 16].reshape(number_sdbs, 1), \
                                data[:, 17:20], \
                                data[:, 20:23], \
                                data[:, 23:26], \
                                data[:, 26:29]

fc = np.concatenate(((wt/input).reshape(number_sdbs*replicates, 1),
                     (wt_n/input).reshape(number_sdbs*replicates, 1),
                     (cmas/input).reshape(number_sdbs*replicates, 1),
                     (cmas_n/input).reshape(number_sdbs*replicates, 1),
                     (input).reshape(number_sdbs,1)))
cells = np.array(['WT']*number_sdbs*replicates + ['WT_n']*number_sdbs*replicates +
                 ['CMAS']*number_sdbs*replicates + ['CMAS_n']*number_sdbs*replicates + ['Input']*number_sdbs).\
    reshape((number_sdbs*replicates*(targets+1)-(number_sdbs*2), 1))

lectin = np.array([lectin for lectin in data[:,2] for _ in range(replicates)]*targets +
                  [lectin for lectin in data[:,2]]).\
    reshape((number_sdbs*replicates*(targets+1))-(number_sdbs*2), 1)

df = pd.DataFrame(np.concatenate((cells, lectin, fc), axis=1), columns=['cells', 'lectin', 'fold'])
df_cells = df.loc[(df['cells'] != 'Input')]
df_input = df.loc[(df['cells'] == 'Input')]


def mbp_norm(df_cell):
    df2_cell = df_cell.copy()
    mbp_values = df2_cell.loc[df2_cell['lectin'].isin(['MBP'])]
    interval = int(mbp_values.shape[0] / targets)
    means = [mbp_values[0:interval]['fold'].mean(), mbp_values[interval:interval * 2]['fold'].mean(),
             mbp_values[interval * 2:interval * 3]['fold'].mean(), mbp_values[interval * 3:interval * 4]['fold'].mean()]

    new_fc = []
    n = 0
    counter = 0
    for x in df2_cell['fold']:

        if counter < (number_sdbs * replicates - 1):
            new_fc.append(x / means[n])
            counter += 1
        else:
            new_fc.append(x / means[n])
            n += 1
            counter = 0

    df2_cell['fold'] = new_fc

    return df2_cell

def quadratic_scale(df_cell):
    new_fc = []
    for x in df_cell['fold']:
        new_fc.append(x**(1/2))

    df_cell['fold'] = new_fc

    return df_cell

test = mbp_norm(df_cells)

# Part 2 - Export data to an excel file

export = pd.read_csv(file_path).fillna(0)
fc1, fc2, fc3, ix, nfc1, nfc2, nfc3  = df_cells[::3], df_cells[1::3], df_cells[2::3], df_input, \
                                 test[::3], test[1::3], test[2::3]

everything = [fc1, fc2, fc3, ix, nfc1, nfc2, nfc3]

for replicate in everything:
    for cell_type in set(replicate['cells']):
        desired_column = replicate.loc[replicate['cells'] == cell_type]['fold']
        desired_column = desired_column.rename(cell_type).reset_index(drop=True)
        export = pd.concat([export,desired_column], axis=1)

export.to_excel(figure, freeze_panes=(1,4))


# PART 3 - Plot the graph

if __name__ == '__main__':

    r1, r2, r3 = test[::3], test[1::3], test[2::3]

    #sns.set(rc={'figure.figsize': (3.5, 3.5)}, style='ticks', )
    fig, ax = plt.subplots(figsize=(4.5,1.5))
    plot = sns.barplot(x='cells', y='fold',
                        hue='lectin',
                        data=test,
                       palette=['#a06236', '#c67942', '#dc9969' , '#e4af8a', '#f1d7c6',
                                '#506da5', '#ffffff'],
                       edgecolor='black',
                       ci=95, errcolor='black', capsize=0.01, errwidth=1

                       )
    sns.stripplot(x='cells', y='fold', hue='lectin',
                         jitter=0.2, dodge=True, data=r1,
                         palette=['#000000'
                                  ], edgecolor='black', linewidth=1, size=3)

    sns.stripplot(x='cells', y='fold', hue='lectin',
                         jitter=0.2, dodge=True, data=r2,
                         palette=['#808080'
                                  ], edgecolor='black', linewidth=1, size=3)

    sns.stripplot(x='cells', y='fold', hue='lectin',
                         jitter=0.2, dodge=True, data=r3,
                         palette=['#ffffff'
                                  ], edgecolor='black', linewidth=1, size=3)


    plt.yscale('log')
    plt.xlabel("WT")
    plt.ylabel("Fold Change (FC)")
    plt.ylim(0.05, 10000)
    labels = [0.05, 0.1, 1, 10, 100, 1000, 10000]
    plt.yticks(labels, labels)
    plt.legend('')
    plt.savefig('GL-VI_148-neuraminidase.pdf', transparent=True)
    plt.show()
```
