## Supplementary material for "Multivalent DNA-encoded lectins on phage enable detecting compositional glycocalyx differences": GL_VI_182_pbmc_illumina.py

```
import pandas as pd
import os
import numpy as np
import matplotlib.pyplot as plt
import seaborn as sns
import math

plt.rcParams.update({'font.size': 7, 'font.family': 'arial', 'pdf.fonttype': 42})
pd.set_option('display.max_rows', None)

file_name = 'GL_VI_182_pbmc_illumina_master_list.csv'
file_path = os.path.join(os.getcwd(), file_name)
dictionary = 'GE.xlsx'
number_sdbs = pd.read_excel(os.path.join(os.getcwd(), dictionary)).shape[0]
replicates = 3
targets = 4
figure = 'Fig3b_SFig4.xlsx'


### STEP 1

# 1 - Creates a master list of all SDBs and associated lectins
if not os.path.exists(file_path):

    master_list = pd.read_excel(os.path.join(os.getcwd(), dictionary)).iloc[:,[0,2,5]].set_index('SDB')

    # 2 - Opens the folder containing all sequencing files
    folder = os.path.join(os.getcwd(), 'Sequencing Files', file_name[:-16])
    files = os.listdir(folder)
    for file in files:

        # 3 - Creates panda dataframe from illumina file
        data = pd.read_csv(f'{folder}/{file}', sep=' ', header=9)
        primers = list(data.columns.values)[6:]

        # 4 - Parses through each primer combination (one replicate)
        for reads in primers:

            total_reads = int(data[reads][0])

            # 5 - Combine reads with same mindex and delete all mindex != 0
            y = 0
            for x in data['mindex']:
                if x != 0:
                    data.loc[data.index == x, reads] += data[reads][y]
                y += 1
            mindex_0 = data[data['mindex'] == 0]


            # 6 - Creates a list of desirable SDBs and corresponding reads in ppm
            df = []
            for x in master_list['Sequence']:

                for y in mindex_0['Nuc']:
                    if x.upper() == y:
                        raw = (mindex_0.loc[mindex_0['Nuc'] == y][reads].tolist()[0])
                        ppm = ((mindex_0.loc[mindex_0['Nuc'] == y][reads].tolist()[0])/total_reads)*1000000
                        sequence = master_list.loc[master_list['Sequence'] == x].index[0]

                        df.append([raw, ppm, sequence])
            df = pd.DataFrame(df, columns=[reads[:-7]+'_Raw', reads[:-7]+'_CPM', 'SDB']).set_index('SDB')

            # 7 - Export data to master list


            master_list= pd.concat([master_list, df], axis=1)

            print(f'File {file}, read {reads} completed')
            print(master_list)

    # Separate column of raw reads from CPM and puts values in the correct order (i.e. input comes first followed
    # by the corresponding targets
    raw_cpm = [0,1] + list(range(2, master_list.shape[1], 2)) + list(range(3, master_list.shape[1], 2))
    master_list.iloc[:,raw_cpm].to_csv(file_path)


### PART 2

plt.rcParams.update({'font.size': 7, 'font.family': 'arial', 'pdf.fonttype': 42})

# 1 - Import data from step 1, fill Nan with 0 and convert to numpy array
data = pd.read_csv(file_path).fillna(0).to_numpy()


# 4 - Obtain Fold changes
tcell, bcell, nk, monocyte, input = data[:, 16:19], data[:, 19:22], data[:, 22:25],data[:, 25:28], \
                                    data[:, 28].reshape(number_sdbs, 1)


fc = np.concatenate(((tcell / input).reshape(number_sdbs*replicates, 1),
                     (bcell / input).reshape(number_sdbs*replicates, 1),
                     (nk / input).reshape(number_sdbs*replicates, 1),
                     (monocyte / input).reshape(number_sdbs*replicates, 1),
                     input.reshape(number_sdbs,1)))

cells = np.array(['T-cells']*number_sdbs*replicates +
                 ['B-cells']*number_sdbs*replicates +
                 ['NK']*number_sdbs*replicates +
                 ['Monocytes']*number_sdbs*replicates +
                 ['Input']*number_sdbs).reshape((number_sdbs*replicates*(targets+1)-(number_sdbs*2), 1))

lectin = np.array([lectin for lectin in data[:,2] for _ in range(replicates)]*targets +
                  [lectin for lectin in data[:,2]]).\
    reshape((number_sdbs*replicates*(targets+1))-(number_sdbs*2), 1)

df = pd.DataFrame(np.concatenate((cells, lectin, fc), axis=1), columns=['cells', 'lectin', 'fold'])
df_cells = df.loc[(df['cells'] != 'Input')]
df_input = df.loc[(df['cells'] == 'Input')]

def mbp_norm(df_cell):
    df2_cell = df_cell.copy()
    mbp_values = df2_cell.loc[df2_cell['lectin'].isin(['MBP'])]
    interval = int(mbp_values.shape[0] / targets)
    means = [mbp_values[0:interval]['fold'].mean(), mbp_values[interval:interval * 2]['fold'].mean(),
             mbp_values[interval * 2:interval * 3]['fold'].mean(), mbp_values[interval * 3:interval * 4]['fold'].mean()]

    new_fc = []
    n = 0
    counter = 0
    for x in df2_cell['fold']:

        if counter < (number_sdbs * replicates - 1):
            new_fc.append(x / means[n])
            counter += 1
        else:
            new_fc.append(x / means[n])
            n += 1
            counter = 0

    df2_cell['fold'] = new_fc

    return df2_cell

def quadratic_scale(df_cell):
    new_fc = []
    for x in df_cell['fold']:
        new_fc.append(x**(1/2))

    df_cell['fold'] = new_fc

    return df_cell

test = mbp_norm(df_cells)

export = pd.read_csv(file_path).fillna(0)
fc1, fc2, fc3, ix, nfc1, nfc2, nfc3  = df_cells[::3], df_cells[1::3], df_cells[2::3], df_input, \
                                 test[::3], test[1::3], test[2::3]

everything = [fc1, fc2, fc3, ix, nfc1, nfc2, nfc3]

for replicate in everything:
    for cell_type in set(replicate['cells']):
        desired_column = replicate.loc[replicate['cells'] == cell_type]['fold']
        desired_column = desired_column.rename(cell_type).reset_index(drop=True)
        export = pd.concat([export,desired_column], axis=1)

export.to_excel(figure, freeze_panes=(1,4))

test = mbp_norm(df_cells)
test2 = quadratic_scale(mbp_norm(df_cells))


# Make graphs for one lectin and compare btw the 4 cell lines

if __name__ == '__main__':

    r1, r2, r3 = test2[::3], test2[1::3], test2[2::3]

    sns.set(rc={'figure.figsize': (3.5, 3.5)}, style='ticks', )
    order=['NK', 'Monocytes', 'T-cells', 'B-cells']

    plot = sns.catplot(x='cells', y='fold', order=order,
                        hue='lectin',
                       kind='bar', data=test2,
                       palette=['#a06236', '#c67942', '#dc9969', '#e4af8a', '#f1d7c6','#506da5',
                                '#71ad44', '#d6e2cd',
                                '#c2c4c6', '#ffffff'],
                       edgecolor='black',
                       ci=95, errcolor='black', capsize=0.01, errwidth=1,
                       legend=False, aspect=6 / 3
                       )
    plot = sns.stripplot(x='cells', y='fold', hue='lectin', order=order,
                         jitter=0.2, dodge=True, data=r1,
                         palette=['#000000'
                                  ], edgecolor='black', linewidth=1)

    plot = sns.stripplot(x='cells', y='fold', hue='lectin', order=order,
                         jitter=0.2, dodge=True, data=r2,
                         palette=['#808080'
                                  ], edgecolor='black', linewidth=1)

    plot = sns.stripplot(x='cells', y='fold', hue='lectin', order=order,
                         jitter=0.2, dodge=True, data=r3,
                         palette=['#ffffff'
                                  ], edgecolor='black', linewidth=1)


    labels = [0, 1, 5, 20, 50, 100, 200, 300]
    ticks = [x ** (1 / 2) for x in labels]
    plt.yticks(ticks, labels)
    plt.legend('')
    plt.savefig('your_plot.pdf', transparent=True)
    plt.show()
```
