## Supplementary material for "Multivalent DNA-encoded lectins on phage enable detecting compositional glycocalyx differences": GL_VII_105_illumina_pbmc_post_sorting.py

```
import pandas as pd
import os
import numpy as np
import matplotlib.pyplot as plt
import seaborn as sns

plt.rcParams.update({'font.size': 7, 'font.family': 'arial', 'pdf.fonttype': 42})
pd.set_option('display.max_rows', None)

file_name = 'GL_VII_105_illumina_pbmc_post_sorting_master_list.csv'
file_path = os.path.join(os.getcwd(), file_name)
dictionary = 'GF.xlsx'
number_sdbs = pd.read_excel(os.path.join(os.getcwd(), dictionary)).shape[0]
replicates = 4
targets = 4
figure = 'EFig3_SFig5.xlsx'

### PART 1

# 1 - Creates a master list of all SDBs and associated lectins
if not os.path.exists(file_path):

    master_list = pd.read_excel(os.path.join(os.getcwd(), dictionary)).iloc[:, [0, 2, 5]].set_index('SDB')

    # 2 - Opens the folder containing all sequencing files
    folder = os.path.join(os.getcwd(), 'Sequencing Files', file_name[:-16])
    files = os.listdir(folder)
    for file in files:

        # 3 - Creates panda dataframe from illumina file
        data = pd.read_csv(f'{folder}/{file}', sep=' ', header=9)
        primers = list(data.columns.values)[6:]

        # 4 - Parses through each primer combination (one replicate)
        for reads in primers:
            if file == '20230825-1728LJooBO-GL.txt' and reads not in ['R5F18_RN1RP1', 'R5F19_RN1RP2',
                                                                  'R5F20_RN1RP3', 'R6F1_RN1RP4']:
                pass
            else:
                total_reads = int(data[reads][0])

                # 5 - Combine reads with same mindex and delete all mindex != 0
                y = 0
                for x in data['mindex']:
                    if x != 0:
                        data.loc[data.index == x, reads] += data[reads][y]
                    y += 1
                mindex_0 = data[data['mindex'] == 0]

                # 6 - Creates a list of desirable SDBs and corresponding reads in ppm
                df = []
                for x in master_list['Sequence']:

                    for y in mindex_0['Nuc']:
                        if x.upper() == y:
                            raw = (mindex_0.loc[mindex_0['Nuc'] == y][reads].tolist()[0])
                            ppm = ((mindex_0.loc[mindex_0['Nuc'] == y][reads].tolist()[0]) / total_reads) * 1000000
                            sequence = master_list.loc[master_list['Sequence'] == x].index[0]

                            df.append([raw, ppm, sequence])
                df = pd.DataFrame(df, columns=[reads[:-7] + '_Raw', reads[:-7] + '_CPM', 'SDB']).set_index('SDB')

                # 7 - Export data to master list

                master_list = pd.concat([master_list, df], axis=1)

                print(f'File {file}, read {reads} completed')
                print(master_list)

    # Separate column of raw reads from CPM and puts values in the correct order (i.e. input comes first followed
    # by the corresponding targets
    raw_cpm = [0, 1] + list(range(2, master_list.shape[1], 2)) + list(range(3, master_list.shape[1], 2))
    master_list.iloc[:, raw_cpm].to_csv(file_path)

### PART 2


# 1 - Import data from step 1, fill Nan with 0 and convert to numpy array
data = pd.read_csv(file_path).fillna(0).to_numpy()

# 4 - Obtain Fold changes
input, tcell, bcell, nkcell, monocyte = np.mean(data[:,39:43], axis=1).reshape(number_sdbs, 1), \
                                        data[:, 23:27], data[:, 27:31], data[:, 31:35], data[:, 35:39]


fc = np.concatenate((np.divide(tcell, input, out=np.zeros_like(tcell), where=input!=0).reshape(number_sdbs*replicates, 1),
                     np.divide(bcell, input, out=np.zeros_like(bcell), where=input!=0).reshape(number_sdbs*replicates, 1),
                     np.divide(nkcell, input, out=np.zeros_like(nkcell), where=input!=0).reshape(number_sdbs*replicates,1),
                     np.divide(monocyte, input, out=np.zeros_like(monocyte), where=input!=0).reshape(number_sdbs*replicates,1),
                     input.reshape(number_sdbs,1)))

cells = np.array(['T-cells']*number_sdbs*replicates +
                 ['B-cells']*number_sdbs*replicates +
                 ['NK']*number_sdbs*replicates +
                 ['Monocytes']*number_sdbs*replicates +
                 ['Input']*number_sdbs).\
    reshape((number_sdbs*replicates*(targets+1)-(number_sdbs*3), 1))

lectin = np.array([lectin for lectin in data[:,2] for _ in range(replicates)]*targets +
                  [lectin for lectin in data[:,2]]).\
    reshape((number_sdbs*replicates*(targets+1))-(number_sdbs*3), 1)

df = pd.DataFrame(np.concatenate((cells, lectin, fc), axis=1), columns=['cells', 'lectin', 'fold'])
df_cells = df.loc[(df['cells'] != 'Input')]
df_input = df.loc[(df['cells'] == 'Input')]


def mbp_norm(df_cell):
    mbp_values = df_cell.loc[df_cell['lectin'].isin(['MBP'])]
    interval = int(mbp_values.shape[0]/targets)
    means = [mbp_values[0:interval]['fold'].mean(), mbp_values[interval:interval*2]['fold'].mean(),
             mbp_values[interval*2:interval*3]['fold'].mean(), mbp_values[interval*3:interval*4]['fold'].mean()]

    new_fc = []
    n = 0
    counter = 0
    for x in df_cell['fold']:

        if counter < (number_sdbs*replicates-1):
            new_fc.append(x/means[n])
            counter += 1
        else:
            new_fc.append(x / means[n])
            n += 1
            counter = 0

    df_cell['fold'] = new_fc

    return df_cell


def quadratic_scale(df_cell):
    new_fc = []
    for x in df_cell['fold']:
        new_fc.append(x**(1/2))

    df_cell['fold'] = new_fc

    return df_cell

test = mbp_norm(df_cells)

export = pd.read_csv(file_path).fillna(0)
fc1, fc2, fc3, ix, nfc1, nfc2, nfc3  = df_cells[::3], df_cells[1::3], df_cells[2::3], df_input, \
                                 test[::3], test[1::3], test[2::3]

everything = [fc1, fc2, fc3, ix, nfc1, nfc2, nfc3]

for replicate in everything:
    for cell_type in set(replicate['cells']):
        desired_column = replicate.loc[replicate['cells'] == cell_type]['fold']
        desired_column = desired_column.rename(cell_type).reset_index(drop=True)
        export = pd.concat([export,desired_column], axis=1)

export.to_excel(figure, freeze_panes=(1,4))


test = quadratic_scale(test)

if __name__ == '__main__':

    r1, r2, r3, r4 = test[::4], test[1::4], test[2::4], test[3::4]
    # 5 - Plot the graphs
    order = ['T-cells', 'B-cells', 'NK', 'Monocytes']
    lectin_order = [#'diCBM40-[290]', 'diCBM40-[145]', 'diCBM40-[73]', 'diCBM40-[26]', 'diCBM40-[3]',
                   # 'MBP'
                      '(Siglec-7)2',  'Siglec-7', 'Siglec-7R'
                    ]

    # sns.set(rc={'figure.figsize': (10, 2.31)}, style='ticks')
    fig, ax = plt.subplots(figsize=(8,3))
    plot = sns.barplot(x='cells', y='fold', hue='lectin',
                       hue_order=lectin_order,
                       order = order,
                       data=test,
                       palette=[#'#a06236', '#c67942', '#dc9969', '#e4af8a', '#f1d7c6','#506da5',

                                '#44AD56','#71ad44', '#d6e2cd'
                           #,
                            #    '#c2c4c6', '#ffffff'
                                ],
                       edgecolor='black',
                       ci=95, errcolor='black', capsize=0.01, errwidth=1.5

                       )


    sns.stripplot(x='cells', y='fold', hue='lectin', hue_order=lectin_order,
                  order=order,
                         jitter=0.2, dodge=True, data=r1,
                         palette=['#000000'
                                  ], edgecolor='black', linewidth=1, size=3)

    sns.stripplot(x='cells', y='fold', hue='lectin', hue_order=lectin_order,
                  order=order,

                         jitter=0.2, dodge=True, data=r2,
                         palette=['#808080'
                                  ], edgecolor='black', linewidth=1, size=3)

    sns.stripplot(x='cells', y='fold', hue='lectin',
                  order=order,
                         jitter=0.2, dodge=True, data=r3, hue_order=lectin_order,
                         palette=['#ABABAB'
                                  ], edgecolor='black', linewidth=1, size=3)

    sns.stripplot(x='cells', y='fold', hue='lectin',
                  order=order,
                         jitter=0.2, dodge=True, data=r4, hue_order=lectin_order,
                         palette=['#ffffff'
                                  ], edgecolor='black', linewidth=1, size=3)


    plt.xlabel("Target")
    plt.ylabel("Fold Change (FC)")
    plt.legend('')
    plt.ylim(0, 1600 ** (1 / 2))
    #plt.ylim(1000 ** (1/2), 1600 ** (1/2))
    #labels = [1000, 1200, 1600]
    labels = [0, 1, 5, 20, 50, 100, 200, 300, 500, 1000, 1200, 1600]
    ticks = [x ** (1 / 2) for x in labels]
    plt.yticks(ticks, labels)

    plt.savefig('GL-VII-105 pbmcs_EF3b_both.pdf', transparent=True)

    plt.show()
```
