## Supplementary material for "Multivalent DNA-encoded lectins on phage enable detecting compositional glycocalyx differences": GL_VII_108_drug_treatments.py

```
import pandas as pd
import os
import numpy as np
import matplotlib.pyplot as plt
import seaborn as sns

pd.set_option('display.max_rows', None)


file_name = 'GL_VII_108_drug_treatments_master_list.csv'
file_path = os.path.join(os.getcwd(), file_name)
dictionary = 'GF.xlsx'
number_sdbs = pd.read_excel(os.path.join(os.getcwd(), dictionary)).shape[0]
replicates = 3
targets = 5
figure = 'SFig7.xlsx'

### PART 1

### PART 1

# 1 - Creates a master list of all SDBs and associated lectins
if not os.path.exists(file_path):

    master_list = pd.read_excel(os.path.join(os.getcwd(), dictionary)).iloc[:,[0,2,5]].set_index('SDB')

    # 2 - Opens the folder containing all sequencing files
    folder = os.path.join(os.getcwd(), 'Sequencing Files', file_name[:-16])
    files = os.listdir(folder)
    for file in files:
        # 3 - Creates panda dataframe from illumina file
        data = pd.read_csv(f'{folder}/{file}', sep=' ', header=9)
        primers = list(data.columns.values)[6:]

        # 4 - Parses through each primer combination (one replicate)
        for reads in primers:
            if file == '20230825-1728LJooBO-GL.txt' and reads not in ['R6F15_RN1RP1', 'R6F16_RN1RP2', 'R6F17_RN1RP3']:
                pass
            else:
                total_reads = int(data[reads][0])

                # 5 - Combine reads with same mindex and delete all mindex != 0
                y = 0
                for x in data['mindex']:
                    if x != 0:
                        data.loc[data.index == x, reads] += data[reads][y]
                    y += 1
                mindex_0 = data[data['mindex'] == 0]


                # 6 - Creates a list of desirable SDBs and corresponding reads in ppm
                df = []
                for x in master_list['Sequence']:

                    for y in mindex_0['Nuc']:
                        if x.upper() == y:
                            raw = (mindex_0.loc[mindex_0['Nuc'] == y][reads].tolist()[0])
                            ppm = ((mindex_0.loc[mindex_0['Nuc'] == y][reads].tolist()[0])/total_reads)*1000000
                            sequence = master_list.loc[master_list['Sequence'] == x].index[0]

                            df.append([raw, ppm, sequence])
                df = pd.DataFrame(df, columns=[reads[:-7]+'_Raw', reads[:-7]+'_CPM', 'SDB']).set_index('SDB')

                # 7 - Export data to master list


                master_list= pd.concat([master_list, df], axis=1)

                print(f'File {file}, read {reads} completed')
                print(master_list)

    # Separate column of raw reads from CPM and puts values in the correct order (i.e. input comes first followed
    # by the corresponding targets
    raw_cpm = [0,1] + list(range(2, master_list.shape[1], 2)) + list(range(3, master_list.shape[1], 2))
    master_list.iloc[:,raw_cpm].to_csv(file_path)

### PART 2

# 1 - Import data from step 1, fill Nan with 0 and convert to numpy array
data = pd.read_csv(file_path).fillna(0).to_numpy()

# 4 - Obtain Fold changes
input, dmso, ara, aza, ven, azaven = np.mean(data[:, 36:39], axis=1).reshape(number_sdbs, 1), \
                                     data[:, 21:24], data[:, 24:27], data[:, 27:30], \
                                     data[:, 30:33], data[:, 33:36]


fc = np.concatenate((
                     np.divide(dmso, input, out=np.zeros_like(dmso), where=input!=0).reshape(number_sdbs*replicates,1),
                     np.divide(ara, input, out=np.zeros_like(ara), where=input!=0).reshape(number_sdbs*replicates,1),
                     np.divide(aza, input, out=np.zeros_like(aza), where=input!=0).reshape(number_sdbs*replicates, 1),
                     np.divide(ven, input, out=np.zeros_like(ven), where=input!=0).reshape(number_sdbs*replicates,1),
                     np.divide(azaven, input, out=np.zeros_like(azaven), where=input!=0).reshape(number_sdbs*replicates,1),
                     input.reshape(number_sdbs, 1)
                     ))


cells = np.array(
                 ['+DMSO']*number_sdbs*replicates + ['+750 nM Cytarabine (Ara-C)']*number_sdbs*replicates +
                 ['+15 uM Azacitidine']*number_sdbs*replicates + ['+25 nM Venetoclax']*number_sdbs*replicates +
                 ['+5 uM Azacitidine +15 nM Venetoclax']*number_sdbs*replicates + ['Input']*number_sdbs).\
    reshape((number_sdbs*replicates*(targets+1)-(number_sdbs*2), 1))

lectin = np.array([lectin for lectin in data[:,2] for _ in range(replicates)]*targets +
                  [lectin for lectin in data[:,2]]).\
    reshape((number_sdbs*replicates*(targets+1))-(number_sdbs*2), 1)


df = pd.DataFrame(np.concatenate((cells, lectin, fc), axis=1), columns=['cells', 'lectin', 'fold'])
df_cells = df.loc[(df['cells'] != 'Input')]
df_input = df.loc[(df['cells'] == 'Input')]


def mbp_norm(df_cell):
    mbp_values = df_cell.loc[df_cell['lectin'].isin(['MBP'])]
    interval = int(mbp_values.shape[0]/targets)
    means = [mbp_values[0:interval]['fold'].mean(), mbp_values[interval:interval*2]['fold'].mean(),
             mbp_values[interval*2:interval*3]['fold'].mean(), mbp_values[interval*3:interval*4]['fold'].mean(),
             mbp_values[interval*4:interval*5]['fold'].mean()]

    new_fc = []
    n = 0
    counter = 0
    for x in df_cell['fold']:

        if counter < (number_sdbs*replicates-1):
            new_fc.append(x/means[n])
            counter += 1
        else:
            new_fc.append(x / means[n])
            n += 1
            counter = 0

    df_cell['fold'] = new_fc

    return df_cell


test = mbp_norm(df_cells)

export = pd.read_csv(file_path).fillna(0)
fc1, fc2, fc3, ix, nfc1, nfc2, nfc3  = df_cells[::3], df_cells[1::3], df_cells[2::3], df_input, \
                                 test[::3], test[1::3], test[2::3]

everything = [fc1, fc2, fc3, ix, nfc1, nfc2, nfc3]

for replicate in everything:
    for cell_type in set(replicate['cells']):
        desired_column = replicate.loc[replicate['cells'] == cell_type]['fold']
        desired_column = desired_column.rename(cell_type).reset_index(drop=True)
        export = pd.concat([export,desired_column], axis=1)

export.to_excel(figure, freeze_panes=(1,4))


if __name__ == '__main__':


    r1, r2, r3= test[::3], test[1::3], test[2::3]
    fig, ax = plt.subplots(figsize=(12,3))
    order = ['+DMSO', '+25 nM Venetoclax', '+15 uM Azacitidine',
             '+750 nM Cytarabine (Ara-C)', '+5 uM Azacitidine +15 nM Venetoclax']
    plot = sns.barplot(x='cells', y='fold', hue='lectin', order=order,
                          data=test,
                       palette=['#a06236', '#c67942', '#dc9969', '#e4af8a', '#f1d7c6','#506da5',
                                '#71ad44', '#d6e2cd', '#44AD56',
                                '#c2c4c6', '#ffffff'
                                ],
                       edgecolor='black',
                       ci=95, errcolor='black', capsize=0.01, errwidth=1.5
                       )

    sns.stripplot(x='cells', y='fold', hue='lectin',
                         order=order,
                         jitter=0.2, dodge=True, data=r1,
                         palette=['#000000'
                                  ], edgecolor='black', linewidth=1)

    sns.stripplot(x='cells', y='fold', hue='lectin',
                         order=order,
                         jitter=0.2, dodge=True, data=r2,
                         palette=['#5C5C5C'
                                  ], edgecolor='black', linewidth=1)

    sns.stripplot(x='cells', y='fold', hue='lectin',
                         order=order,
                         jitter=0.2, dodge=True, data=r3,
                         palette=['#ffffff'
                                  ], edgecolor='black', linewidth=1)


    plt.xlabel("Target")
    plt.ylabel("Fold Change (FC)")
    plt.yscale('log')
    plt.legend('')
    labels = [0.01, 0.1, 1, 10, 100, 1000, 10000]
    plt.savefig('GL-VII-108 cscs treatment_double_drugs.pdf', transparent=True)

    plt.show()
```
