## Supplementary material for "Multivalent DNA-encoded lectins on phage enable detecting compositional glycocalyx differences": GL_VII_139_lila_invivo.py

```
import pandas as pd
import os
import numpy as np
import matplotlib.pyplot as plt
import seaborn as sns


plt.rcParams.update({'font.size': 7, 'font.family': 'arial', 'pdf.fonttype': 42})
pd.set_option('display.max_rows', None)

file_name = 'GL_VII_139_lila_invivo_master_list.csv'
file_path = os.path.join(os.getcwd(), file_name)
dictionary = 'GF.xlsx'
number_sdbs = pd.read_excel(os.path.join(os.getcwd(), dictionary)).shape[0]
replicates = 3
number_lectins = 11
targets = 10
figure = 'Fig5c-d.xlsx'

### STEP 1

# 1 - Creates a master list of all SDBs and associated lectins
if not os.path.exists(file_path):
    master_list = pd.read_excel(os.path.join(os.getcwd(), dictionary)).iloc[:,[0,2,5]].set_index('SDB')

    # 2 - Opens the folder containing all sequencing files
    folder = os.path.join(os.getcwd(), 'Sequencing Files', file_name[:-16])
    files = os.listdir(folder)
    for file in files:

        # 3 - Creates panda dataframe from illumina file
        data = pd.read_csv(f'{folder}/{file}', sep=' ', header=9)
        primers = list(data.columns.values)[6:]

        # 4 - Parses through each primer combination (one replicate)
        for reads in primers:

            total_reads = int(data[reads][0])

            # 5 - Combine reads with same mindex and delete all mindex != 0
            y = 0
            for x in data['mindex']:
                if x != 0:
                    data.loc[data.index == x, reads] += data[reads][y]
                y += 1
            mindex_0 = data[data['mindex'] == 0]


            # 6 - Creates a list of desirable SDBs and corresponding reads in ppm
            df = []
            for x in master_list['Sequence']:

                for y in mindex_0['Nuc']:
                    if x.upper() == y:
                        raw = (mindex_0.loc[mindex_0['Nuc'] == y][reads].tolist()[0])
                        ppm = ((mindex_0.loc[mindex_0['Nuc'] == y][reads].tolist()[0])/total_reads)*1000000
                        sequence = master_list.loc[master_list['Sequence'] == x].index[0]

                        df.append([raw, ppm, sequence])
            df = pd.DataFrame(df, columns=[reads[:-7]+'_Raw', reads[:-7]+'_CPM', 'SDB']).set_index('SDB')

            # 7 - Export data to master list
            master_list= pd.concat([master_list, df], axis=1)

            print(f'File {file}, read {reads} completed')
            print(master_list)

    # Separate column of raw reads from CPM
    raw_cpm = [0,1] + list(range(2, master_list.shape[1], 2)) + list(range(3, master_list.shape[1], 2))
    master_list.iloc[:,raw_cpm].to_csv(file_path)


### PART 2

plt.rcParams.update({'font.size': 7, 'font.family': 'arial', 'pdf.fonttype': 42})

# 1 - Import data from step 1, fill Nan with 0 and convert to numpy array
data = pd.read_csv(file_path).fillna(0).to_numpy()

export = pd.read_csv(file_path).fillna(0)
# 4 - Obtain Fold changes

# 4.1 Get CPM counts
input, lungs, heart, spleen, kidneys, liver, plasma, tcell, bcell, rbc, hf = np.mean(data[:,62:65], axis=1).reshape(number_sdbs, 1), \
                                                                        data[:, 59:62], data[:, 44:47], data[:, 47:50], \
                                                                        data[:, 50:53], data[:, 35:38], data[:, 38:41], \
                                                                        data[:, 53:56], data[:, 56:59], data[:, 41:44], data[:,34].reshape(number_sdbs,1)
x = number_sdbs*replicates
cpm = np.concatenate((lungs.reshape(x, 1), heart.reshape(x, 1), spleen.reshape(x, 1),
                        kidneys.reshape(x, 1), liver.reshape(x, 1), plasma.reshape(x, 1),
                        tcell.reshape(x, 1), bcell.reshape(x, 1), rbc.reshape(x, 1), input,
                        hf.reshape(number_sdbs,1)))

# 4.2 Create array with the organs and respective constructs, include input
organs = np.array(['Lungs']*number_sdbs*replicates +
                  ['Heart']*number_sdbs*replicates +
                 ['Spleen']*number_sdbs*replicates + ['Kidneys']*number_sdbs*replicates +
                 ['Liver']*number_sdbs*replicates + ['Plasma']*number_sdbs*replicates +
                 ['T-cells']*number_sdbs*replicates + ['B-cells']*number_sdbs*replicates +
                 ['RBCs']*number_sdbs*replicates + ['Input']*number_sdbs + ['HF']*number_sdbs).\
    reshape((number_sdbs*replicates*targets)-(number_sdbs), 1)

lectin = np.array([lectin for lectin in data[:,2] for _ in range(replicates)]*(targets-1) +
                  [lectin for lectin in data[:,2]]*2).\
    reshape((number_sdbs*replicates*targets)-(number_sdbs*1), 1)

df = pd.DataFrame(np.concatenate((organs, lectin, cpm), axis=1), columns=['organs', 'lectin', 'cpm'])

def indexing(df):
    index_list = []
    for x in df['Axis name']:
        if x not in index_list:
            index_list.append(x)
    indices = []
    for y in index_list:
        indices.append(df.index[df['Axis name'] == y].tolist()[0])
    return indices

# 4.3 Replace misbehaving SDBs with 'NaN'
cpm_values = df['cpm'].values

clean_df = df.copy()
for i in range(0, len(cpm_values), 3):
    if np.all(cpm_values[i:i+3] == 0):
        clean_df.iloc[i:i+3, -1] = np.nan


# 4.3 Average all SDBs from a given replicate and assign indices

organs_df = clean_df.loc[(clean_df['organs'] != 'Input') & (clean_df['organs'] != 'HF')]

input_df = clean_df.loc[df['organs'] == 'Input']

mouse1, mouse2, mouse3 = organs_df[0::3].groupby(['organs', 'lectin'], sort=False)['cpm'].mean().reset_index(),\
                         organs_df[1::3].groupby(['organs', 'lectin'], sort=False)['cpm'].mean().reset_index(),\
                         organs_df[2::3].groupby(['organs', 'lectin'], sort=False)['cpm'].mean().reset_index()
inputx = input_df.groupby(['organs', 'lectin'], sort=False)['cpm'].mean().reset_index()


# 4.4 Normalize everything to MBP

def mbp_norm(df_cell):
    mbp_values = df_cell.loc[df_cell['lectin'].isin(['MBP'])]['cpm'].reset_index(drop=True)
    new_fc = []
    n = 0
    counter = 0
    for x in df_cell['cpm']:
        if counter < number_lectins-1:
            new_fc.append(x/mbp_values[n])
            counter += 1
        else:
            new_fc.append(x / mbp_values[n])
            n += 1
            counter = 0

    df_cell['cpm'] = new_fc

    return df_cell

m1, m2, m3, i = mbp_norm(mouse1), mbp_norm(mouse2), mbp_norm(mouse3), mbp_norm(inputx)

def concatenation(df, y):
    for replicate in df:
        for x in set(replicate['organs']):
            a = replicate.loc[replicate['organs'] == x]
            cpm_column = a.drop('organs', axis=1).set_index(pd.Index(indexing(y))).iloc[:,-1:]
            cpm_column = cpm_column.rename(columns={cpm_column.columns[0]: x})

            y = pd.concat([y,cpm_column
                                ], axis=1)
    return y

[]


input_values = i['cpm'].to_numpy().reshape(1,-1)

fc1, fc2, fc3 = m1.copy(), m2.copy(), m3.copy()
samples = [fc1, fc2, fc3]

# 4.5 Divide each mouse to input

for x in samples:
    values = x['cpm'].to_numpy().reshape(targets-1, number_lectins)
    x['final'] = (values/input_values).reshape(-1,1)

everything = [m1, m2, m3, i, fc1, fc2, fc3]
export = concatenation(everything, export).to_excel(figure, freeze_panes=(1,4))

# PART 3

df = pd.concat([fc1, fc2, fc3])

def quadratic_scale(df_cell):
    new_fc = []
    for x in df_cell['final']:
        new_fc.append(x**(1/2))

    df_cell['final'] = new_fc

    return df_cell

test = df
if __name__ == '__main__':


    # 5 - Plot the graphs

    #sns.set(rc={'figure.figsize': (3.5, 3.5)}, style='ticks', )
    figure, ax = plt.subplots(figsize=(9,3))
    plot = sns.barplot(x='organs', y='final',
                        hue='lectin',

                       data=test,

                       palette=['#a06236', '#c67942', '#dc9969', '#e4af8a', '#f1d7c6','#506da5',
                                '#71ad44', '#d6e2cd', '#44AD56',
                               '#c2c4c6', '#ffffff'
                                ],
                       edgecolor='black', ci=False,
                        errcolor='black', capsize=0.01, errwidth=1

                       )
    plot = sns.stripplot(x='organs', y='final', hue='lectin',

                         jitter=0.2, dodge=True, data=test,

                         palette=['#000000'
                                  ], edgecolor='black', linewidth=1)
"""
    plot = sns.stripplot(x='cells', y='fold', hue='lectin',
                         order=['rbc', 'tcell', 'bcell', 'plasma'],
                         hue_order=['Sig7di', 'MBP'],
                         jitter=0.2, dodge=True, data=r2,
                         palette=['#808080'
                                  ], edgecolor='black', linewidth=1)

    plot = sns.stripplot(x='cells', y='fold', hue='lectin',
                         order=['rbc', 'tcell', 'bcell', 'plasma'],
                         hue_order=['Sig7di', 'MBP'],
                         jitter=0.2, dodge=True, data=r3,
                         palette=['#ffffff'
                                  ], edgecolor='black', linewidth=1)
"""
plt.xlabel("Organs")


plt.yscale('log')
plt.ylabel("Fold Change (FC)")


plt.legend('')
plt.savefig('GL-VII-139 invivo-cells-sig-upper.pdf', transparent=True)


plt.show()
```
