## Supplementary material for "Multivalent DNA-encoded lectins on phage enable detecting compositional glycocalyx differences": GL_VII_147_toxicity_targeting_curve.py

```
import matplotlib.pyplot as plt
import pandas as pd
import seaborn as sns
# Obtain FC values from the cells

import GL_VI_148_lila_density_neuraminidase, GL_VI_182_pbmc_illumina, \
    GL_VI_148_lila_v_1_4_csc, GL_VII_108_illumina_CHST1_and_CHST2


plt.rcParams.update({'font.size': 7, 'font.family': 'arial', 'pdf.fonttype': 42})

density = GL_VI_148_lila_density_neuraminidase.test
pbmcs = GL_VI_182_pbmc_illumina.test
mv4 = GL_VI_148_lila_v_1_4_csc.test
chsts = GL_VII_108_illumina_CHST1_and_CHST2.test

df = pd.concat([density, pbmcs, mv4, chsts])
lectin = 'diCBM40-[73]'
order = ['CMAS_n', 'CMAS', 'WT_n', 'B-cells', 'NK', 'Monocytes', 'T-cells', 'SORE+', 'Wt', 'CHST1', 'CHST2']
data = df.loc[df['lectin'] ==lectin].set_index('cells').loc[order].reset_index()
# x, y = data['cells'], data['fold']

#sns.set(rc={'figure.figsize': (2, 3.5)}, style='ticks')
fig, ax = plt.subplots(figsize=(8,2))
r1, r2, r3 = data[::3],data[1::3],data[2::3]

plot = sns.barplot(x='cells', y='fold', hue='lectin',

                    data=data,
                   palette=[#'#c67942'#,
                        '#dc9969'
                             ],

                   edgecolor='black',
                   ci=95, errcolor='black', capsize=0.01, errwidth=1.5

                   )
sns.stripplot(x='cells', y='fold', hue='lectin',
                         jitter=.2, dodge=True, data=r1,
                         palette=['#000000'
                                  ], edgecolor='black', linewidth=1, size=4)

sns.stripplot(x='cells', y='fold', hue='lectin',

                         jitter=0.2, dodge=True, data=r2,
                         palette=['#808080'
                                  ], edgecolor='black', linewidth=1, size=4)

sns.stripplot(x='cells', y='fold', hue='lectin',
                         jitter=0.2, dodge=True, data=r3,
                         palette=['#ffffff'
                                  ], edgecolor='black', linewidth=1, size=4)

plt.yscale('log')
plt.legend('')
plt.ylim(0.08, 100, 200)
labels = [0.1, 1, 10, 100, 200]
plt.yticks(labels, labels)
plt.ylabel("Fold Change (FC)")
plt.savefig(f'{lectin}.pdf', transparent=True)

plt.show()


# Create the plot


"""
This is the xy graph
plt.figure(figsize=(15, 6))
plt.plot(x, y, marker='o', linestyle='-', color='b', label='fold')
plt.xlabel('Cell Types')
plt.ylabel('Fold Change (FC)')
plt.yscale('log')
plt.legend('')
plt.xticks(rotation=45)  # Rotate the labels for better visibility
labels = [1, 10, 100, 1000]
ticks = labels
plt.yticks(ticks, labels)

plt.savefig('cbm50.pdf', transparent=True)
plt.show()


"""
```
