## Supplementary material for "Multivalent DNA-encoded lectins on phage enable detecting compositional glycocalyx differences": GL_VII_152 phage density modeling EC.py

```
#!/usr/bin/env python3

import collections
import gzip
import math
import random
import re
import sys

import matplotlib.pyplot as plt
import numpy as np

density = 26  #290
repeats = int(10000/density)
seed = 263
random.seed(seed)
output_file = gzip.open('phage-{}.cif.gz'.format(seed), 'w')


def read_PDB(filename):
    # read in an atom list from a compressed PDB file
    # ignore anything after the first model
    atom_list = []
    end_of_model = False
    # the field-structure of PDB ATOM and HETATM lines
    coord_line = re.compile(r'(.{6})(.{5}) (....)(.)(...).(.)(....)(.)...(.{8})(.{8})(.{8})(.{6})(.{6}).{10}(..)(..)')

    for line in gzip.open(filename, 'r'):
        line = line.decode()
        if not end_of_model and (line[0:6] == 'ATOM  ' or line[0:6] == 'HETATM'):
            # parse the line into a tuple of character (fixed-width) fields
            atom_list.append(coord_line.match(line).groups())
        elif line[0:6] == 'ENDMDL':
            end_of_model = True

    return atom_list

atom_list = read_PDB('2MJZ.pdb.gz')

### numbers found by fitting to N-terminal amine nitrogens in PDB 2MJZ MODEL 1

protein_radius = 24
rise = 16.6637857142857127  # AAngstroms
twist = +0.639339243817 # radians = -35.368 degrees
symmetry = 2*math.pi / 5 # radians - angle between 5-fold units
radius = 35.080302062362 + protein_radius  # 35.0803 AAngstroms
twist_offset = 1.0532032408269
x_offset = -0.1439143
y_offset = -0.2107714
z_offset = 9.749443


# compute the N-terminal positions across several crystal repeats
position_list = []
for k in range(0, 2700//5):   # 2700 copies for a typical viral capsid
    for theta in range(5):
        angle = twist * k + symmetry * theta + twist_offset
        position_list.append( (k, theta, angle, math.cos(angle), math.sin(angle), rise * k + z_offset) )

def pair_distance(i, j):
    (k_i, th_i, angle_i, sin_i, cos_i, z_i) = i
    (k_j, th_j, angle_j, sin_j, cos_j, z_j) = j

    dot_product = (sin_i * sin_j + cos_i * cos_j)
    """Understanding dot product
    
    Given two vectors A and B which is the radius of the phage. Angles of A and B are
    i and j respectively. 
    By definition dot product is the sum of product of x and y coordinates of two vectors
    Dot product =  yA*yB + xA*xB
    To find the x and y coordinates we have to imagine drawing a triangle, where x and y 
    are the bases and the height (respectively) of the hypotenuse A or B. 
    Therefore yA, xA = Asini, A.cosi
    yB, xB = B.sinj, B.cosj
    
    We replace x and y coordinates with the sin and cos functions: 
    dot product = AB (sini*sinj + cosi*cosj)
    We can remove AB because all we care is whether is positive or negative. 
    dot product = (sini*sinj + cosi*cosj)
    Given two angles where i is 0 and j is 0 + x, the product of sin will always be 0, so:
    dot product = (0 + 1*cosj). 
    For cosj, everything within 180° (i.e. same side of the phage) will be >= 0. 
    
    Now, if we want to know the exact angle between A and B:
    dot product = AB (sini*sinj + cosi*cosj)
    = AB * cos (i-j)  - see proof here https://www.themathpage.com/aTrig/sum-proof.htm
    
    Since A*B is just a proportionatly factor, we can just treat is as 1 to find the
    angle between these 2 vectors:
    (i-j) = cos-1 (dot product)
       
    """

    if dot_product >= 0 and i != j: # not self-neighbour nor far side
        # for acos() clamp dot-products to max of 1.0 (may round to more)
        delta_angle = math.acos(min(dot_product, 1.0))
        distance = math.sqrt((z_i - z_j)**2 + (radius * delta_angle)**2)
        return distance
    else:
        return 9e99


all_distances = []

for r in range(1):
    randomized_position = position_list.copy()

    random.shuffle(randomized_position)
    counter = 0

    glycan_list = []
    for p in randomized_position:
        if counter == 0:
            glycan_list.append(p)
            counter +=1
        else:
            # check if p conflicts with any already in glycan_list
            distances = [pair_distance(p, x) for x in glycan_list]
            if min(distances) >= protein_radius*2:
                glycan_list.append(p)

            if len(glycan_list) >= density:  # exit if enough already selected
                break

# build dictionary indexed by layers of sets of angles
chosen = collections.defaultdict(set)
for g in glycan_list:
    chosen[g[0]].add(g[1])


# write a minimal CIF header
output_file.write("data_virion\n# \nloop_\n_atom_site.group_PDB\n_atom_site.id\n_atom_site.type_symbol\n_atom_site.label_atom_id\n_atom_site.label_comp_id\n_atom_site.label_asym_id\n_atom_site.label_entity_id\n_atom_site.label_seq_id\n_atom_site.Cartn_x\n_atom_site.Cartn_y\n_atom_site.Cartn_z\n_atom_site.occupancy\n_atom_site.B_iso_or_equiv\n_atom_site.pdbx_PDB_model_num\n".encode())


# write out the atom list transformed into each sucessive position
N = 420  #2700
amine_list = []
atom_number = 1
chain_number = 1
old_chain = ''
for ring in range(0, (N-1)//5, 7):

    # work out a transform matrix
    angle = twist * ring #+ twist_offset
    cosine = math.cos(angle)
    sine = math.sin(angle)
    (xx, xy, xz, xo) = (cosine, -sine, 0, x_offset)
    (yx, yy, yz, yo) = (sine, cosine, 0, y_offset)
    (zx, zy, zz, zo) = (0, 0, 1, rise*ring - 668.476)  # offset for CoM

    # run through the atom list and output transformed coordinates
    for (record, serial, name, altLoc, resName, chainID, resSeq, iCode,
         x, y, z, occupancy, tempFactor, element, charge) in atom_list:


        x = float(x) - x_offset
        y = float(y) - y_offset
        z = float(z)

        xt = xx * x + xy * y + xz * z + xo
        yt = yx * x + yy * y + yz * z + yo
        zt = zx * x + zy * y + zz * z + zo
        # detect the first residues and remember the N-terminal amine sites
        if (name == ' N  ' and resSeq == '   1'):
            amine_list.append( (xt, yt, zt) )

        # update the chain id - if changed
        if chainID != old_chain:
            chain = 'P{}'.format(chain_number)
            chain_number += 1
        # TODO: some of the hard-coded widths may be too small
        output_file.write('ATOM   {:<6d} {:2s} {:4s} {:3s} {:5s} {:<1d} {:<4d} {:<8.3f} {:<8.3f} {:<8.3f} {:<6.2f} {:<6.2f} {:<1d} \n'.format(atom_number,
            element.strip(), name.strip(), resName, chain, 1, int(resSeq),
            xt, yt, zt, float(occupancy), float(tempFactor), 1).encode())
        atom_number += 1
        old_chain = chainID

###print("# ")			# don't end the coordinate table here

#######################################################################
### now write out some amine locations

offset = 10.0  # 1 nm radial offset of the chain S atoms
residue_number = 1
for x, y, z in amine_list:

    # determine if this amine is one of the chosen ones
    kk = ring + int(math.floor(z/rise-0.58 + 0.5))
    theta = 5 / (2 * math.pi) * (math.atan2(y, x) - kk * twist) - 0.83
    theta = int(math.floor(theta + 0.5)) % 5
    if theta in chosen[kk]:

        # compute the offset location
        radius = math.sqrt(x*x + y*y)
        x0 = x * (radius + offset) / radius
        y0 = y * (radius + offset) / radius

        # emit a nitrogen atom there
        output_file.write('ATOM   {:<6d} {:2s} {:4s} {:3s} {:5s} {:<1d} {:<4d} {:<8.3f} {:<8.3f} {:<8.3f} {:<6.2f} {:<6.2f} {:<1d} \n'.format(atom_number,
            'N', 'N', 'ALA', 'S', 1, residue_number, x0, y0, z, 1.0, 0.0, 1).encode())
        atom_number += 1
        residue_number += 1

# end the output file
output_file.write("# \n".encode())
output_file.close()

sys.exit(10)


all_distances = []

for r in range(repeats):
    randomized_position = position_list.copy()
    random.shuffle(randomized_position)
    counter = 0

    glycan_list = []
    for p in randomized_position:
        if counter == 0:
            glycan_list.append(p)
            counter +=1
        else:
            # check if p conflicts with any already in glycan_list
            distances = [pair_distance(p, x) for x in glycan_list]
            if min(distances) >= protein_radius*2:
                glycan_list.append(p)

            if len(glycan_list) >= density:  # exit if enough already selected
                break

    randomized_position = 0
    # for each site find the minimum distances to any other
    neighbours_list = []
    for i, (k_i, th_i, angle_i, sin_i, cos_i, z_i) in enumerate(glycan_list):
        # find distances to all others
        distance_list = []
        for j, (k_j, th_j, angle_j, sin_j, cos_j, z_j) in enumerate(glycan_list):

            # over-kill but this simple algorithm works
            dot_product = (sin_i * sin_j + cos_i * cos_j)
            if dot_product >= 0 and i != j: # not self-neighbour nor far side
                # for acos() clamp dot-products to max 1.0 (rounding noise may
                delta_angle = math.acos(min(dot_product, 1.0)) # be higher)
                distance = math.sqrt((z_i - z_j)**2 + (radius * delta_angle)**2)
                distance_list.append((distance, j))
        if len(distance_list) >= 1 :

            min_distance, j = sorted(distance_list)[0]
            neighbours_list.append((j, min_distance))
    for x in neighbours_list:
        all_distances.append(x[1])
    print(f'{round(r / repeats * 100, 1)}% completed')


# plot the graph

plt.rcParams.update({'font.size': 7, 'font.family': 'arial', 'pdf.fonttype': 42})

distances = [x for x in all_distances if x <=250]
bin_width = 10 # angstroms
min_value = min(distances)
max_value = max(distances)
num_bins = int((max_value - min_value) / bin_width)
"""
print(distances)
a = 5
boxes = list(range(a,100000, a))
numbers = []
for x in boxes:
    counter = 0
    for y in distances:
        if y <=x and y >=(x-a):
            counter+=1
    numbers.append(counter)
non_zero_count = len([x for x in numbers if x != 0])
print(non_zero_count)
"""
fig, ax = plt.subplots(figsize=(5, 1.5))
plt.hist(all_distances, color='#f1d7c6', bins = num_bins, range=(min_value, max_value), edgecolor='black',
         weights=np.ones(len(all_distances)) / len(all_distances)
         )  # You can adjust the number of bins as needed
plt.xlabel('Distance (nm)')
plt.ylabel("")

ticksy = [0.2, 0.4, 0.6, 0.8]
labelsy = [20, 40, 60, 80]
plt.yticks(ticksy, labelsy)

ticksx = [x for x in range(0,250,20)]
labelsx = [x/10 for x in ticksx]
plt.xticks(ticksx, labelsx)


plt.xlim(0,250)

plt.savefig(f'{density}_histogram.pdf', transparent=True)

plt.show()
```
