## Supplementary figures and images for "Multivalent DNA-encoded lectins on phage enable detecting compositional glycocalyx differences"

### EFig. 3a.tif

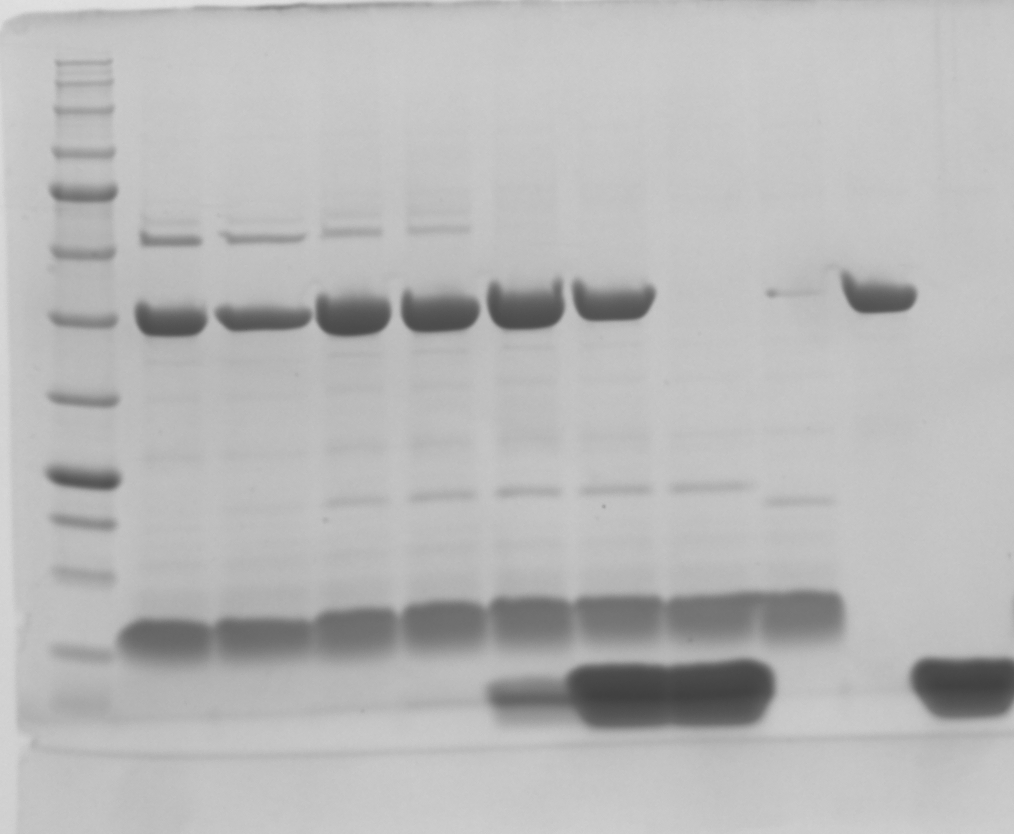

### EFig. 3b.tif

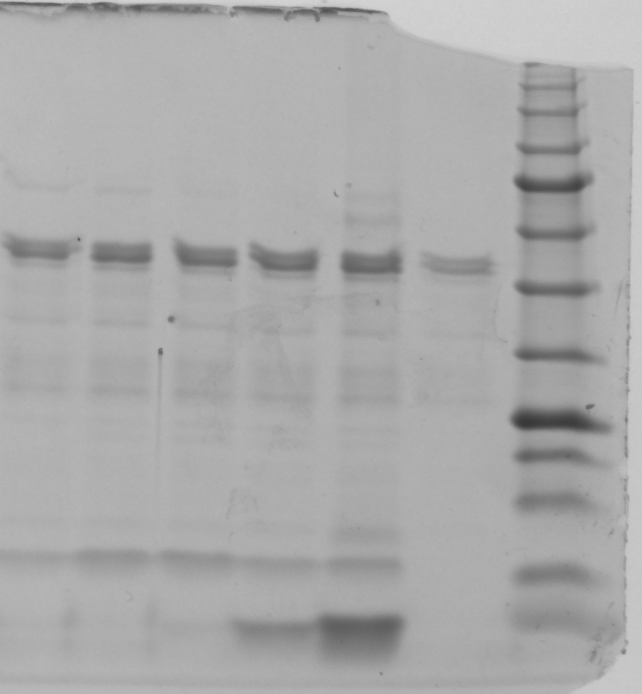

### Fig. 1e and EFig. 2a,c.tif

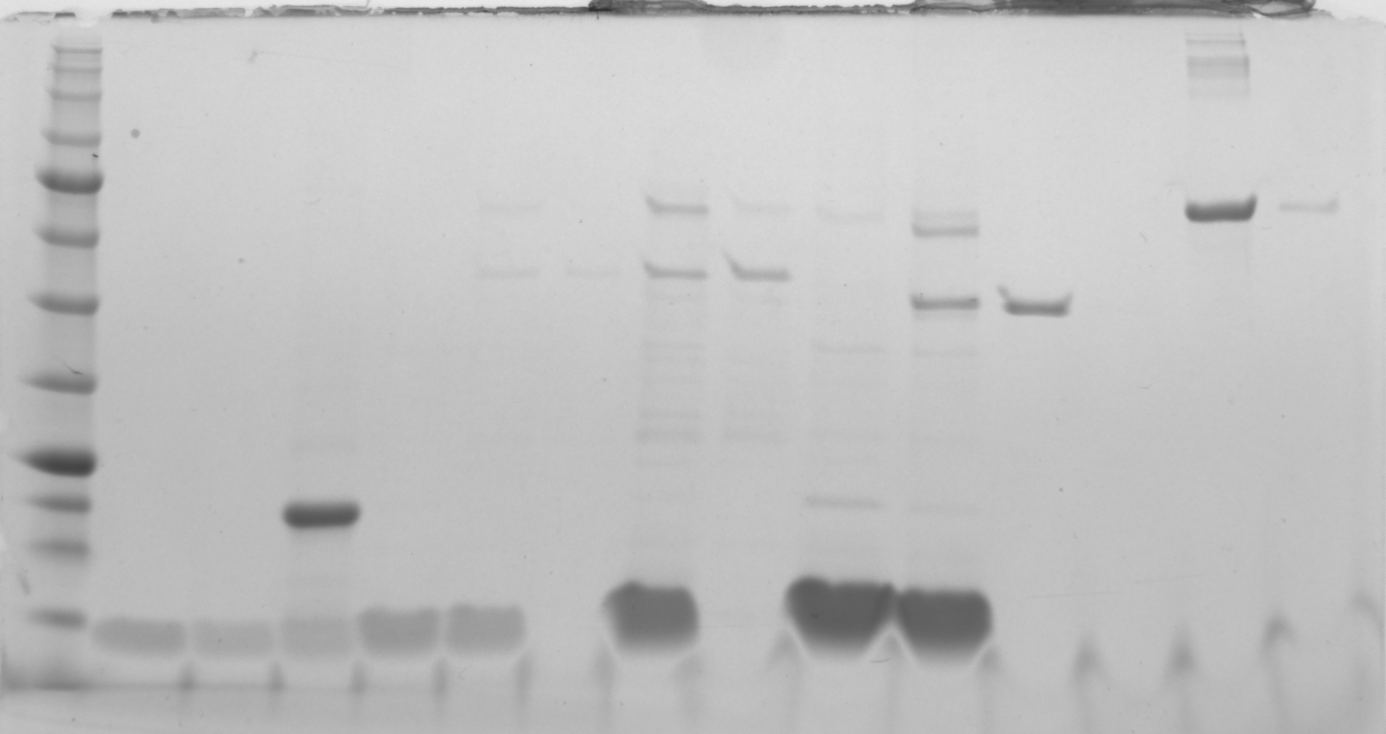

### SFig. 4.tif

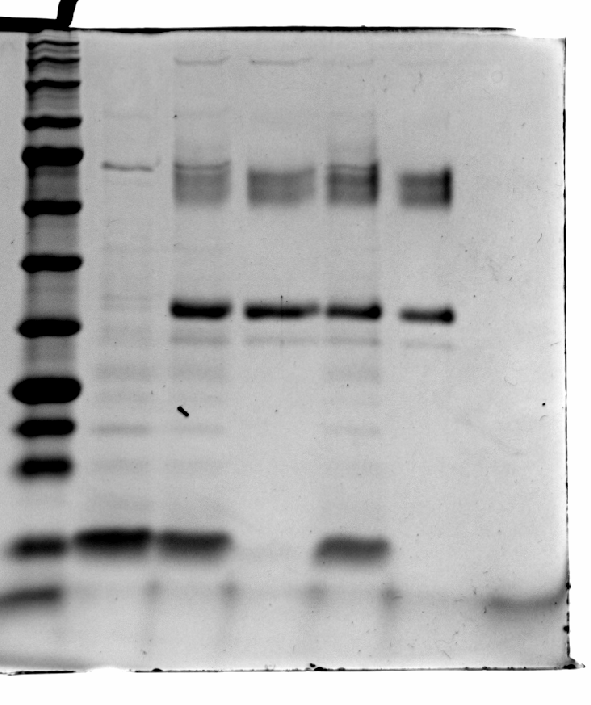

### SFig. 9b.tif

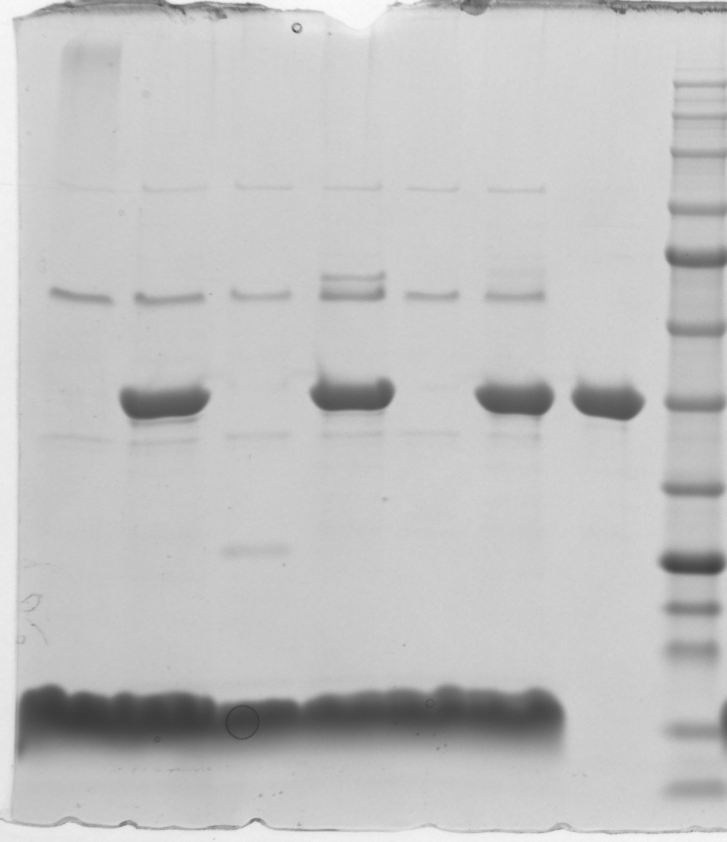
