## Supplementary Information for "Multivalent DNA-encoded lectins on phage enable detecting compositional glycocalyx differences"

**Abbreviation**

| | |
|---|---|
| bp | Base pair |
| dsDNA | Double-stranded DNA |
| DCM | Dichloromethane |
| DIPEA | N,N-Diisopropylethylamine |
| DMSO | Dimethyl sulfoxide |
| DMF | N,N-Dimethylformamide |
| DNA | Deoxyribonucleic acid |
| EDT | 1,2-ethanedithiol |
| EDTA | Ethylenediaminetetraacetic acid |
| dNTPs | Deoxyribonucleotide triphosphates |
| FACS | Fluorescence-activated cell sorting |
| IPTG | Isopropyl $\beta$-D-1-thiogalactopyranoside |
| LB | Lysogeny broth |
| MALDI-TOF | Matrix-assisted laser desorption/ionization – Time of flight |
| SIA | *N*-succinimidyl iodoacetate |
| PBS | Phosphate buffered saline |
| PCR | Polymerase chain reaction |
| PEG | Poly (ethylene glycol) |
| TCEP | Tris(2-carboxyethyl) phosphine |
| TIPS | Triisopropyl silane |

### 1. Synthetic Methods

#### 1.1 *General synthetic methods*

All reagents were purchased from commercial sources and used without further purification. Reagents for peptide synthesis were purchased from ChemPep. Ovendried glassware was used for all reactions. Flash column chromatography was performed using C18 reversed phase 15.5g column (RediSep Gold). A gradient of solvent A (MQ water/TFA 99.9/0.1) and solvent B (MECN) was run at a flow rate of 30 mL/min (0-1.0 min 0% B; 1.0-6.0 min 0 → 35% B; 6.0-7.0 min 35% B; 7.0-12.0 min 35 → 70% B; 12-16 min 70 – 100% B). LCMS analysis of peptide was obtained on Agilent Technologies 6130 LCMS. A gradient of solvent A (MQ water) and solvent B (MeCN/H2O 95/5) was run at a flow rate of 0.5 mL/min (0-4.0 min 5% B; 4.0-5.0 min 5%→60% B; 5.0-5.5 min 60%→100% B; 5.5-7.5 100% B, 7.5-11 min 100%→5% B). Organic solutions were concentrated under vacuum below 40 °C.

#### 1.2 *Synthesis of TAMRA-SpyTag peptide*

The SpyTag peptide were synthesized on an automated peptide synthesizer (Prelude®X; GYROS PROTEIN Technology) using standard solid phase amide coupling. After synthesis the resin was transferred to a Poly-Prep column (BIORAD) and washed with DCM (10 mL) and dried in vacuum. 20 mg resin containing approximately 6 mg of SpyTag peptide was mixed with 3.2 eq. of TAMRA NHS ester, 5-isomer (Lumiprobe, # 57120) in 500 μL DMF. Mixture was incubated for 2h at room temperature with agitation in the dark. *N*-terminally labelled SpyTag-resin was filtered, washed with DCM until the eluate became clear and colorless, and dried in vacuum. The resin was then treated with a cleavage cocktail (0.7 mL) containing TFA/$H_2O$/TIPS/EDT, 90/2.5/5/2.5 (v/v/v/v) for the global deprotection and cleavage of the peptide from the resin for 5h at 200 rpm in the dark. The flow through from the column was collected and the resin was rinsed with TFA (1 mL). The combined cleavage mixture reduced in volume by means of gently bubbling nitrogen through it and 30ml of cold diethyl ether was added in the solution to precipitate the peptide. The precipitate formed was separated by centrifugation (10 min, 3000 rpm). Supernatant was decanted and the precipitated was washed with cold diethyl ether (30 mL). The centrifugation and washing steps were repeated for two more cycles. Residual diethyl ether was removed by nitrogen blow. Then the peptide was dissolved in water for combi flash purification. The fractions corresponding to the main peak were collected (wavelength = 214nm) and submitted for LCMS. Solvent was removed by bubbling nitrogen through it and lyophilized to get the pure TAMRA-SpyTag peptide.

**Supplementary Table S1. List of LiLA mixtures used in this manuscript.**

To access the data concatenate URL as 48hd.cloud/file/20220408-1024OOooBO-GL

| LiLA | Composition details | Naïve composition | Experimental data | Used in |
|---|---|---|---|---|
| GA.xlsx | Blank phages | 20220408-1024OOooBO-GL | 20220408-1024OOuaaRC-GL | S. Fig. 3 |
| GB.xlsx | diCBM40 and blank phages | 20220408-1024CBMCooBO-GL | 20220408-1024CBMCuaaRC-GL | Fig. 1h |
| GC.xlsx | 4 lectins, Sc and blank phage | 20220527-1163LILAAooBO-GL<br>20220527-1164MBPCooBO-GL<br>20220527-1165CBMCooBO-GL<br>20220527-1166SIGCooBO-GL<br>20220527-1167SGRCooBO-GL<br>20220527-1168OOooBO-GL<br>20220527-1169SPYCooBO-GL | 20220527-1163LILAAuaaYX-GL<br>20220527-1163LILAAubaYX-GL<br>20220527-1163LILAAucaYX-GL | Fig. 2e |
| GD.xlsx | diCBM40 displayed at 5 densities, MBP and blank phage | 20230203-1591CBMMBPCooBO-GL | 20230203-1591CBMMBPCuaaYX-GL<br>20230203-1591CBMMBPCuabYX-GL<br>20230203-1591CBMMBPCubaYX-GL<br>20230203-1591CBMMBPCubbYX-GL | Fig. 2 i |
| GE.xlsx | GD + Siglec-7, Siglec-7R and Sc phage | 20230203-1590LILAAooBO-GL<br>20230317-1618LILAAooBO-GL<br>20230515-1618LILAAooBO-GL<br>20230712-1618LILAAooBO-GL | 20230203-1590LILAAsoreaYX-GL<br>20230203-1590LILAAsorebYX-GL<br>20230317-1618LILAAhaaFWA-GL<br>20230317-1618LILAAhbaFWA-GL<br>20230317-1618LILAAhcaFWA-GL<br>20230317-1618LILAAhdaFWA-GL<br>20230515-1618LILAAsupmaYX-GL<br>20230515-1618LILAAsupmbYX-GL<br>20230515-1618LILAAuconnaYX-GL<br>20230515-1618LILAAuconnbYX-GL<br>20230712-1618LILAAmbaFWB-GL | Fig. 3b<br>Fig. 3d<br>Fig. 4c<br>Fig. 4d<br>Fig. 4e |

| | | | | |
|---|---|---|---|---|
| | | | 20230712-1618LILAAmaaFWB-GL | |
| GF.xlsx | GE + diSiglec-7-Fc | 20230825-1728LJooBO-GL | 20230825-1728LJjaaYX-GL | Fig. 2h |
| | | | 20230825-1728LJjbaYX-GL | Fig. 5 |
| | | | 20230825-1728LJmdaYX-GL | S. Fig. 6b |
| | | | 20230825-1728LJmeaYX-GL | S. Fig. 8 |
| | | | 20230825-1728LJmfaYX-GL | Extended Data |
| | | | 20230825-1728LJmgaYX-GL | Fig. 4 |
| | | | 20230825-1728LJmhaYX-GL | |
| | | | 20230825-1728LJhaaFWB-GL | |
| | | | 20230825-1728LJhbaFWB-GL | |
| | | | 20230825-1728LJhcaFWB-GL | |
| | | | 20230825-1728LJhdaFWB-GL | |
| | | | 20230825-1728LJuaaYX-GL | |
| | | | 20230825-1728LJucaYX-GL | |
| | | | 20230825-1728LJudaYX-GL | |
| | | | 20231005-0OOooYXB-GL | |
| | | | 20231005-1728LLbiEYA-GL | |
| | | | 20231005-1728LLcoBLD-GL | |
| | | | 20231005-1728LLcwYXB-GL | |
| | | | 20231005-1728LLhiEYA-GL | |
| | | | 20231005-1728LLikEYA-GL | |
| | | | 20231005-1728LLkoEYA-GL | |
| | | | 20231005-1728LLmaaFWA-GL | |
| | | | 20231005-1728LLmbaFWA-GL | |
| | | | 20231005-1728LLnkEYA-GL | |
| | | | 20231005-1728LLooBO-GL | |
| GF.xlsx | Siglec-7 displayed at various densities | 20230203-1523LILAAooBO-GL | 20230203-1523LILAAuaaYX-GL | Extended Data Fig. 3d |

**Supplementary Table S2.** Sequence of oligos used in this study.

| Name | Sequence |
|---|---|
| P1 | GGAGTTGCGTGATTGTTCTGGTAAAACTATTAG |
| P2 | CTAATAGTTTTACCAGAACAATCACGCAACTCC |
| P3 | GTTTCTTGGTACCCGTGGCGTGCCTCATATC |
| P4 | GTTTCTTGGATCCGGACCACTTTCACCACTACCC |
| P5 | GCCTACAAACGCTATAAGGAGAACCTGTACTTCCAGG |
| P6 | AGCAGCCCCGGGTCACTTCTCGAACTGGGGGTGGGA |
| P7 | AGCAGCACCGGTCGTGGGGTTCCACACATAGTAATGGTGGATGCCTACAAACGCTATAAG |
| P8 | TTTTGGAGATTTTCAACGTG |
| P9 | CCCTCATAGTTAGCGTAACG |
| P10 | CAAGCAGAAGACGGCATACGAGATCGGTCTCGGCATTCCTGCTGAACCGCTCTTCCGATCTNNNNTTGGAGATTTTCAACGTG |
| P11 | AATGATACGGCGACCACCGAGATCTACACTCTTTCCCTACACGACGCTCTTCCGATCTNNNNACAGTTTCGGCCGA |
| P12 | GTTTCTTGGCCAACGTGGCCGGGCCATGGTAACCACCTTATC |
| P13 | GTTTCTTGGCCCCAGAGGCCAAGCTACCACTGGATCCAG |
| P14 | AGGTCTCATGCGGCCATGGTAACCACCTTATC |
| P15 | AGGTCTCACCGCGCTGGAGCTCCCGGATCCTCCACCGCTCTGAGTATGAGCGTCACCTTCAG |
| P16 | AGGTCTCAGCGGAAGGCGATGACCCTGCGAAGGC |

| | |
|---|---|
| P17 | AGGTCTCACGCAAAGCTTAACATTGGG |

**Supplementary Table S3.** Summary of LiLA-FACS campaign.

$n$ = 3 technical replicates for human pre-sorting and murine post-sorting incubation and $n$ = 4 technical replicates for post-sorting incubation. PBMCs were collected from a single volunteer.

| Target Cell | Marker | Fluorophore | # Sorted Cells |
|---|---|---|---|
| Human T-cells | CD3+ | BV650 | 146,641 ± 1004 (Pre-sorting incubation)<br>264,106 ± 37986 (Post-sorting incubation) |
| Human B-cells | CD19+ | APC-Cy7 | 20,652 ± 171 (Pre-sorting incubation)<br>137,750 ± 12659 (Post-sorting incubation) |
| Human NK-cells | CD56+ | BV510 | 32,733 ± 172 (Pre-sorting incubation)<br>13,357 ± 1599 (Post-sorting incubation) |
| Human Monocytes | CD14+ | PE-Cy7 | 50,639 ± 1312 (Pre-sorting incubation)<br>10,523 ±4647 (Post-sorting incubation) |
| Murine T-cells | CD3+ | FITC | 286,602 ± 72189 (Post-sorting incubation) |
| Murine B-Cells | CD19+ | APC | 507,046 ± 100,014 (Post-sorting incubation) |

#### Supplementary Figures

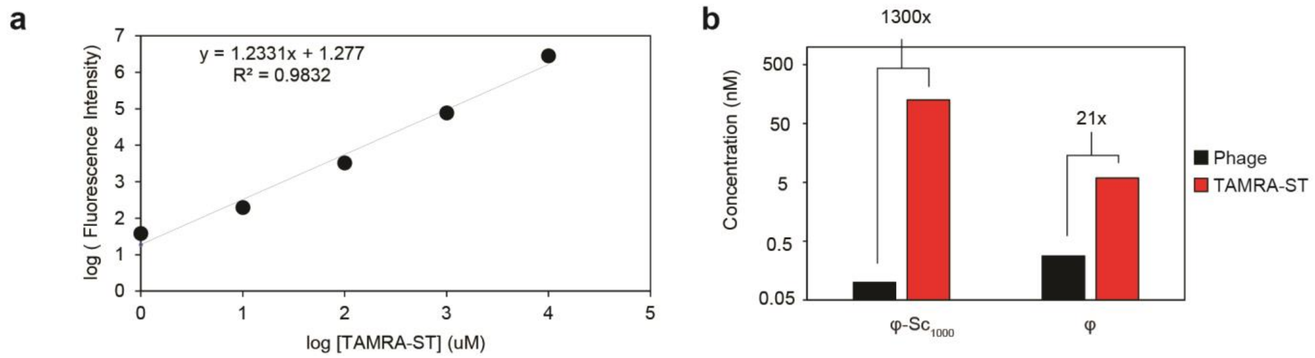

**Supplementary Fig. 1. Synthesis and characterization of TAMRA-ST. a,** TAMRA-ST fluorescence calibration curve. **b,** Fluorescence of φ-$Sc_{1000}$ and blank φ phages after reaction with TAMRA-ST. *n* = 1 experiment.

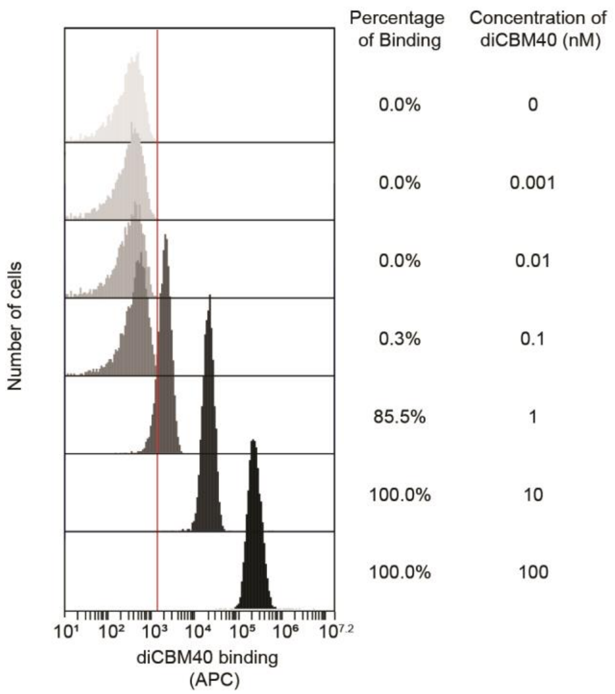

**Supplementary Fig. 2. Probing WT U937 cells with different concentrations of diCBM40.** Flow cytometry results of diCBM40-APC binding to U937 cells. Concentrations ranged from 0 to 100 nM. Binding of diCBM40-APC to the cells were not detectable at the picomolar range.

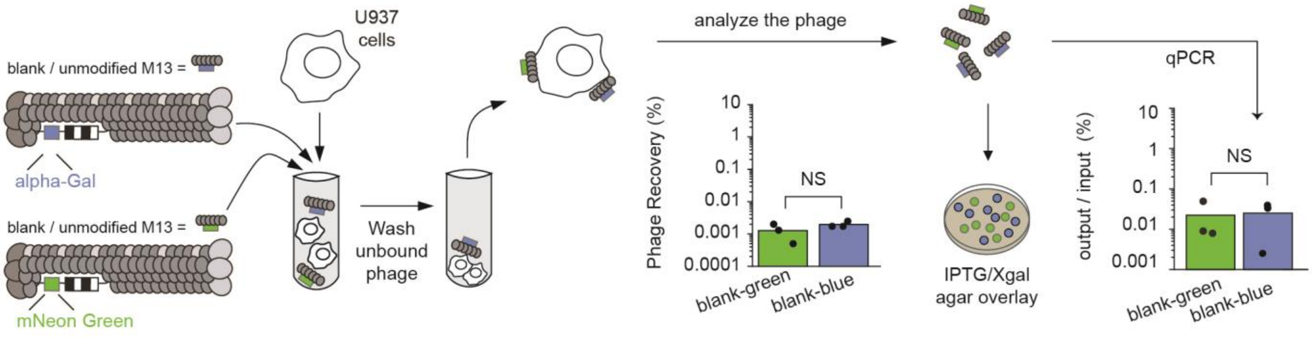

**Supplementary Fig. 3. Binding profile of unmodified reporter phages to WT U937 cells. a,** Recovery of blank phages as measured by plaque assay. Values represent mean of *n* = 3, Mann-Whitney U Test. **b,** Recovery of blank phages as measured by the ratio of DNA reads after (output) and before (input) cell panning. Values represent mean of *n* = 3, Mann Whitney U Test. All respective *n* values are technical replicates. NS, *P* > 0.05.

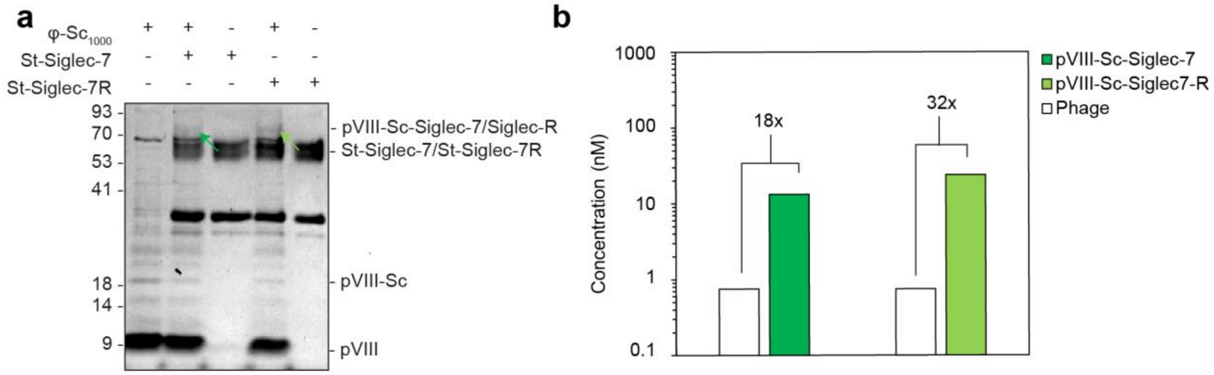

**Supplementary Fig. 4. Copy number estimation of Siglecs on phage. a,** SDS-PAGE of φ-Sc1000 phages conjugated to St-Siglec-7 and St-Siglec-7R. Dark and light green arrows indicate pVIII-Sc protein fused with St-Siglec-7 and St-Siglec-7R, respectively. **b,** Concentration of phages and pVIII-Sc-lectins (pVIII-Sc-Siglec-7 and pVIII-Sc-Siglec-7R) as measured by qPCR and SDS-PAGE densitometry, respectively. *n* = 1 experiment.

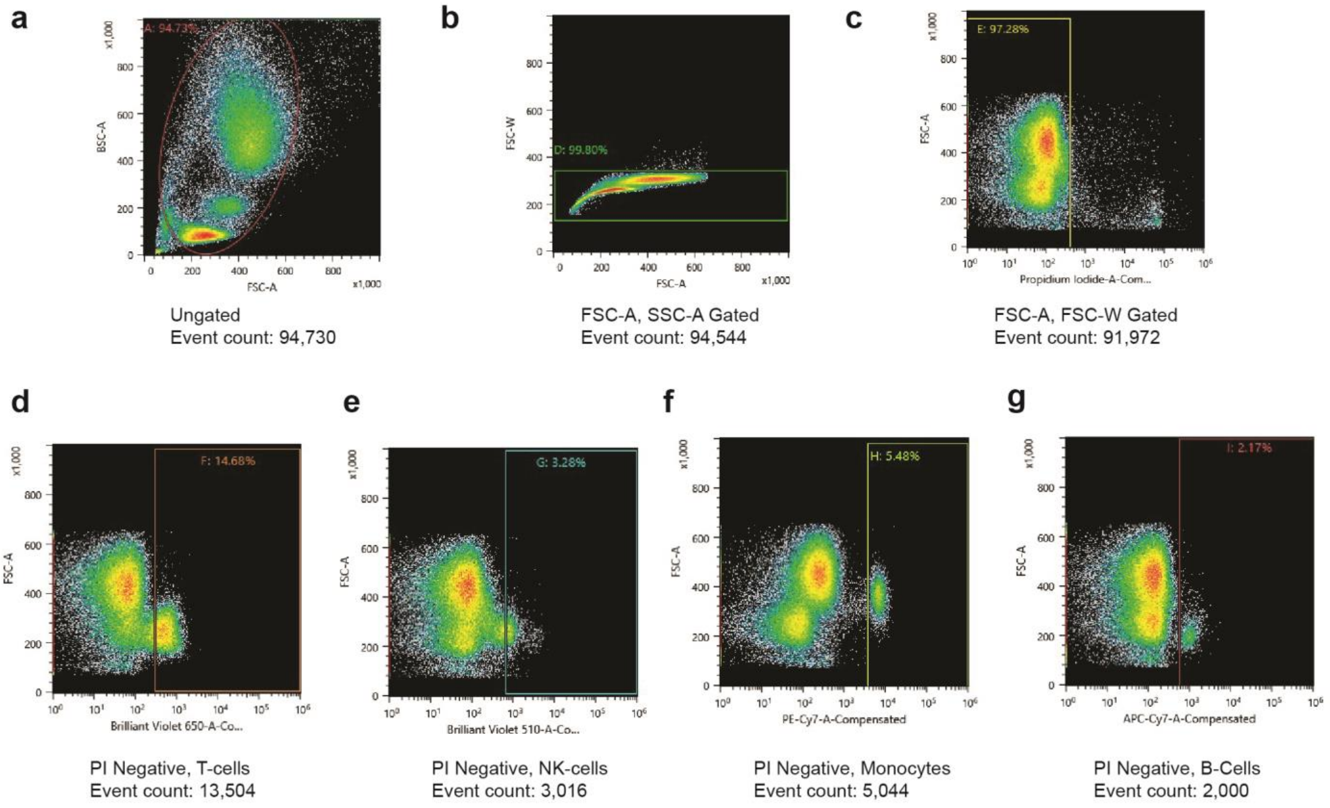

**Supplementary Fig. 5. FACS Gating Strategy.** Gating strategy used in FACS sorting to isolate four different PBMCs from human blood for results depicted in Fig. 3b. **a,b,** Forward and side scatter parameters followed by area and width parameters used to isolate singlets and exclude doublets. **c,** Cell viability dye propidium iodide was used to select negatively stained live cells. **d-g,** Next, PBMCs were gated on each of the desired cell type, based on their respective markers.

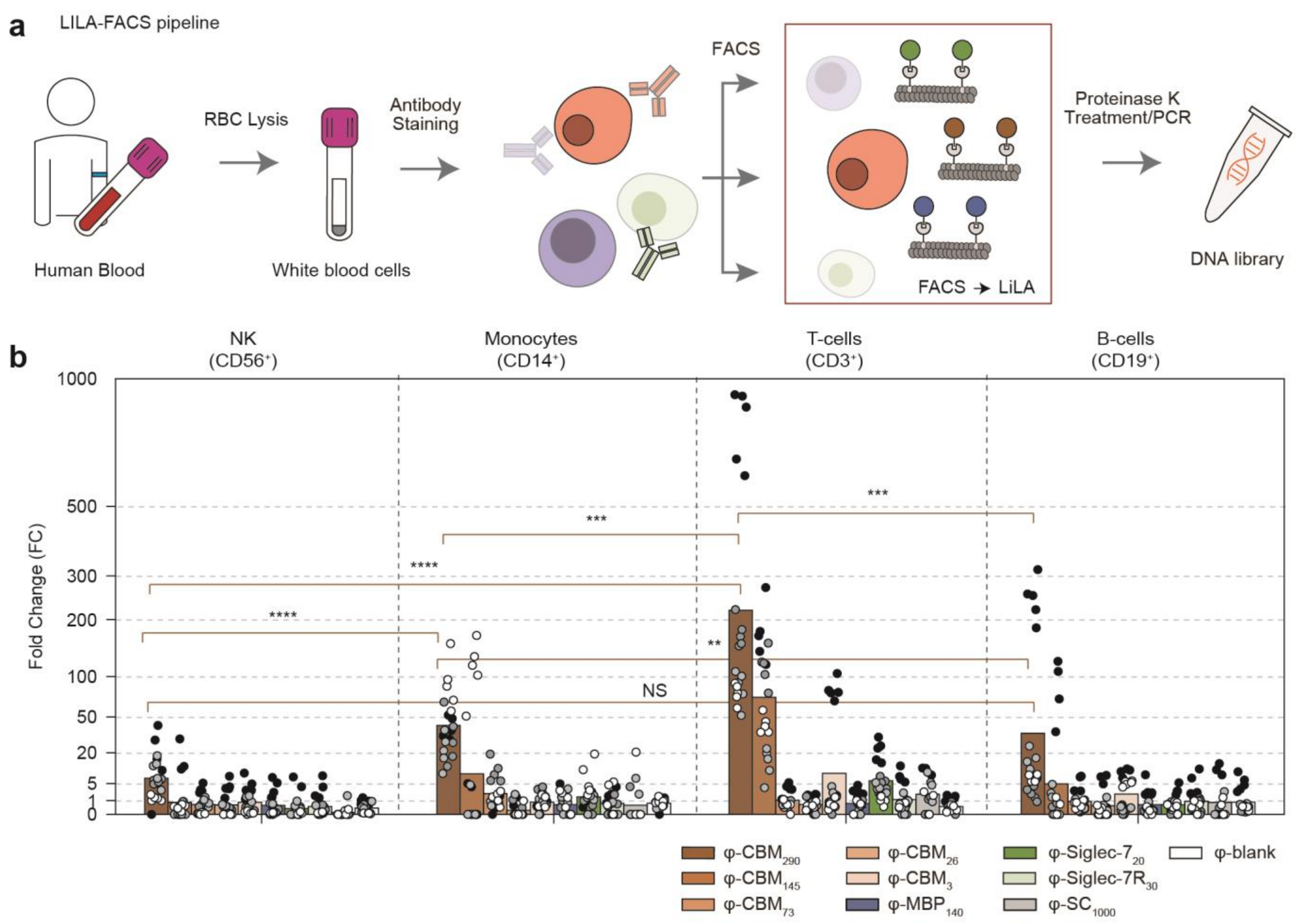

**Supplementary Fig. 6. Glycome profiling of human PBCS by LiLA. a,** Scheme of the FACS→LiLA approach using human PBMCs. **b,** Binding profile of human NK-cells, monocytes, T-cells and B-cells to LiLA. FC of φ-$MBP_{140}$ clones was set to one. Values represent mean of $n = 4$ (technical replicates) x 5 (DNA barcodes per construct). PBMCs were collected from a single volunteer. NS, $P > 0.05$; ** $P < 0.01$; *** $P < 0.001$; **** $P < 0.0001$.

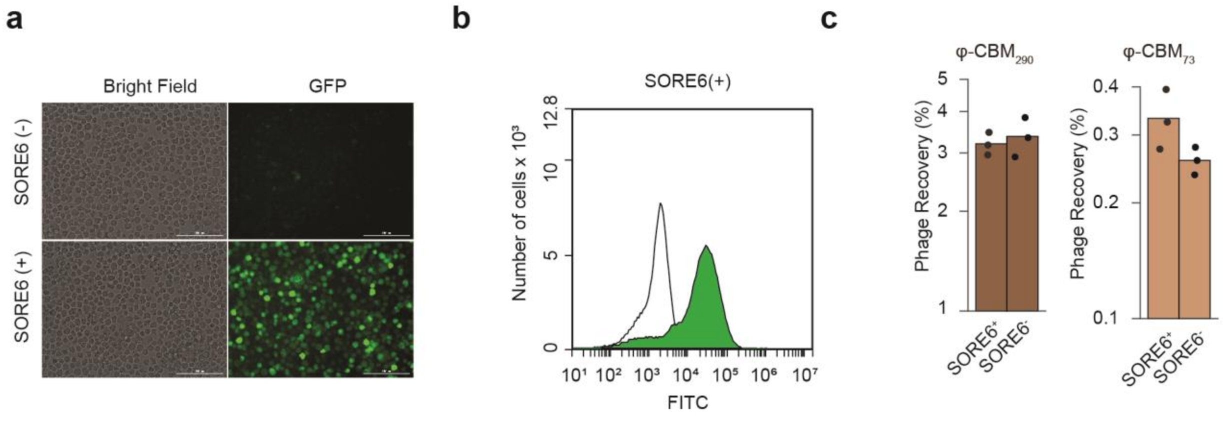

**Supplementary Fig. 7. Characterization of CSC by fluorescence and Lila. a,** Fluorescence cell imaging and **b,** flow cytometry results of MV4-11 cells. **c,** φ-$CBM_{290}$ and φ-$diCBM40_{73}$ enrichment comparisons in MV4-11 cells by titering. Values represent mean of $n$ = 3 technical experiments, Mann-Whitney U test

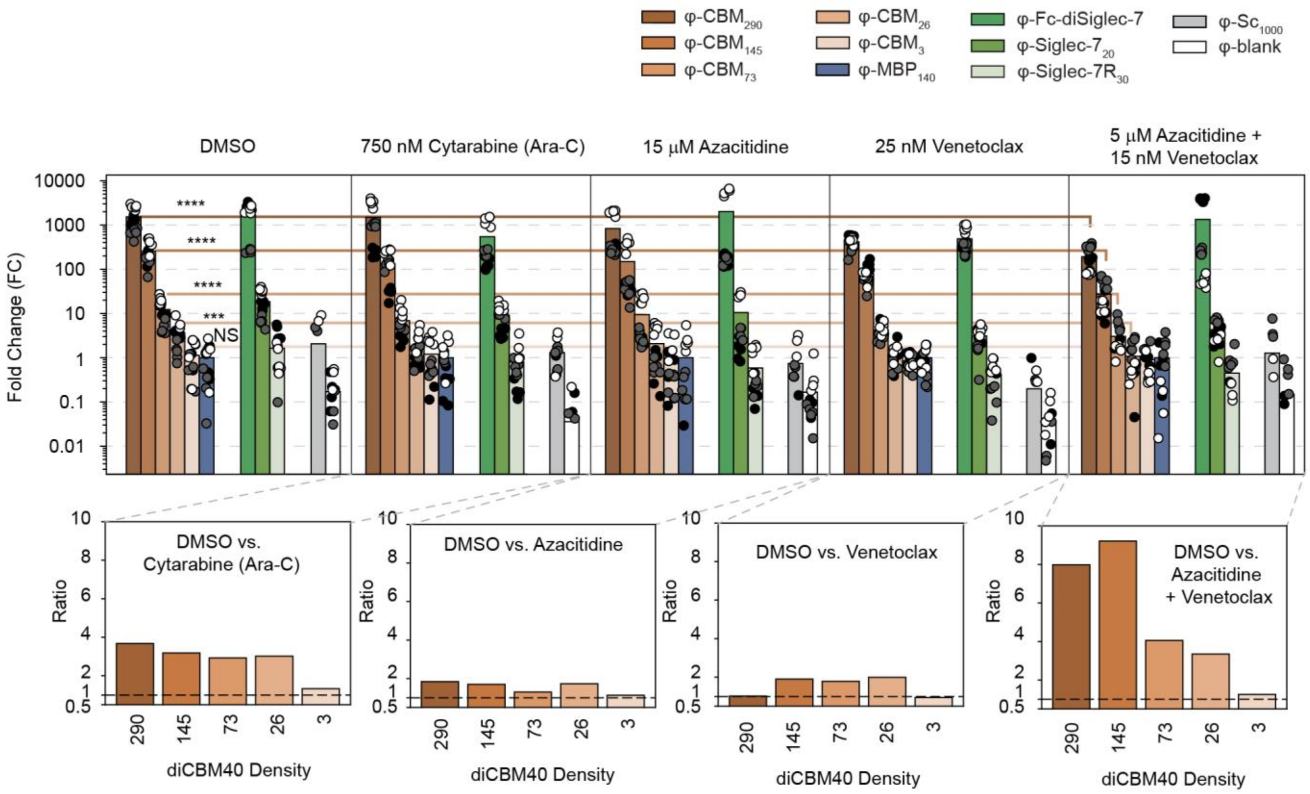

**Supplementary Fig. 8. Binding profile of LiLA on MV4-11 cells submitter to various treatments.** Data is grouped by the type of treatment. Individual doses of Venetoclax, Azacytidine and cytarabine were used or a combination of Venetoclax and Azacitidine. Bottom plot represents ratio of FC values between the different drug treaments. FC of φ-$MBP_{140}$ clones was set to one. Values represent mean of $n$ = 3 technical replicates) × 5 (DNA barcodes per construct). Bottom plots represent ratio of FC values of treatment compared to DMSO control. NS, $P > 0.05$; *** $P < 0.001$, **** $P < 0.0001$.

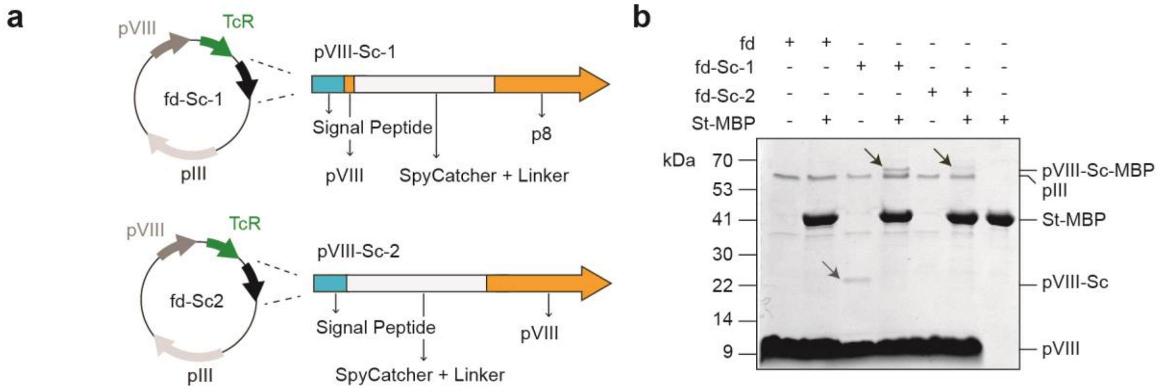

**Supplementary Fig. 9. Cloning and expression of SpyCatcher003 on pVIII. a,** Cloning scheme of the *SpyCatcher003* gene. Digestion of fth1 vector with *SfiI* restriction enzyme yielded fd phages with *SpyCatcher003* gene downstream the first three pVIII amino acids (fd-Sc-1). Digestion of PCR amplified fth1 vector with *BsaI* yielded scarless constructs with *SpyCatcher003* gene downstream the pVIII signal sequence (fd-Sc-2) **b,** SDS-PAGE of fd-Sc constructs conjugated to SpyTag-Maltose-binding protein (MBP). Grey arrow indicates pVIII protein fused with SpyCatcher (pVIII-Sc) while black arrows indicate pVIII-Sc conjugated to MBP (pVIII-Sc-MBP). Band intensities of both pVIII-Sc-MBP products are lower than the pIII protein band intensity. 3-5 copies of the latter are commonly displayed by phages.

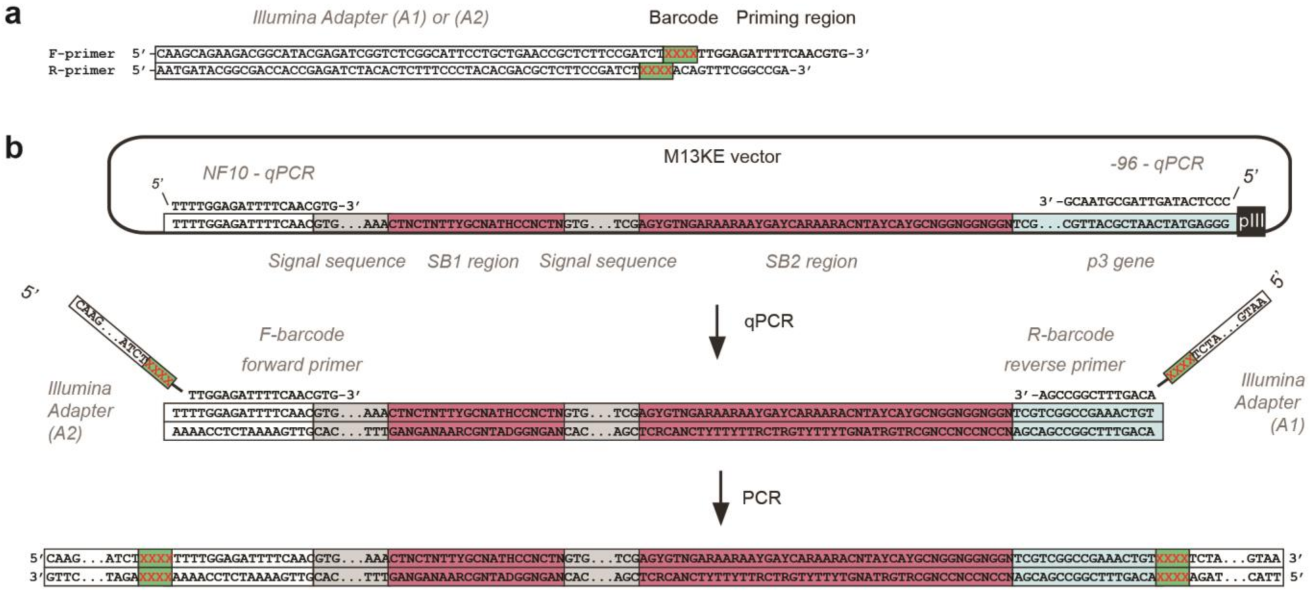

**Supplementary Fig. 10. DNA sequences of 2-step PCR amplification protocol for Illumina deep sequencing. a,** Primers used for amplifying qPCR product. XXXX denotes 4-nucleotide-long barcodes used to trace multiples samples in an Illumina sequencing experiment. **b,** Generation of PCR product. Alignment of forward and reverse qPCR and Illumina primers to sequences flanking the SDB region in M13KE vector.

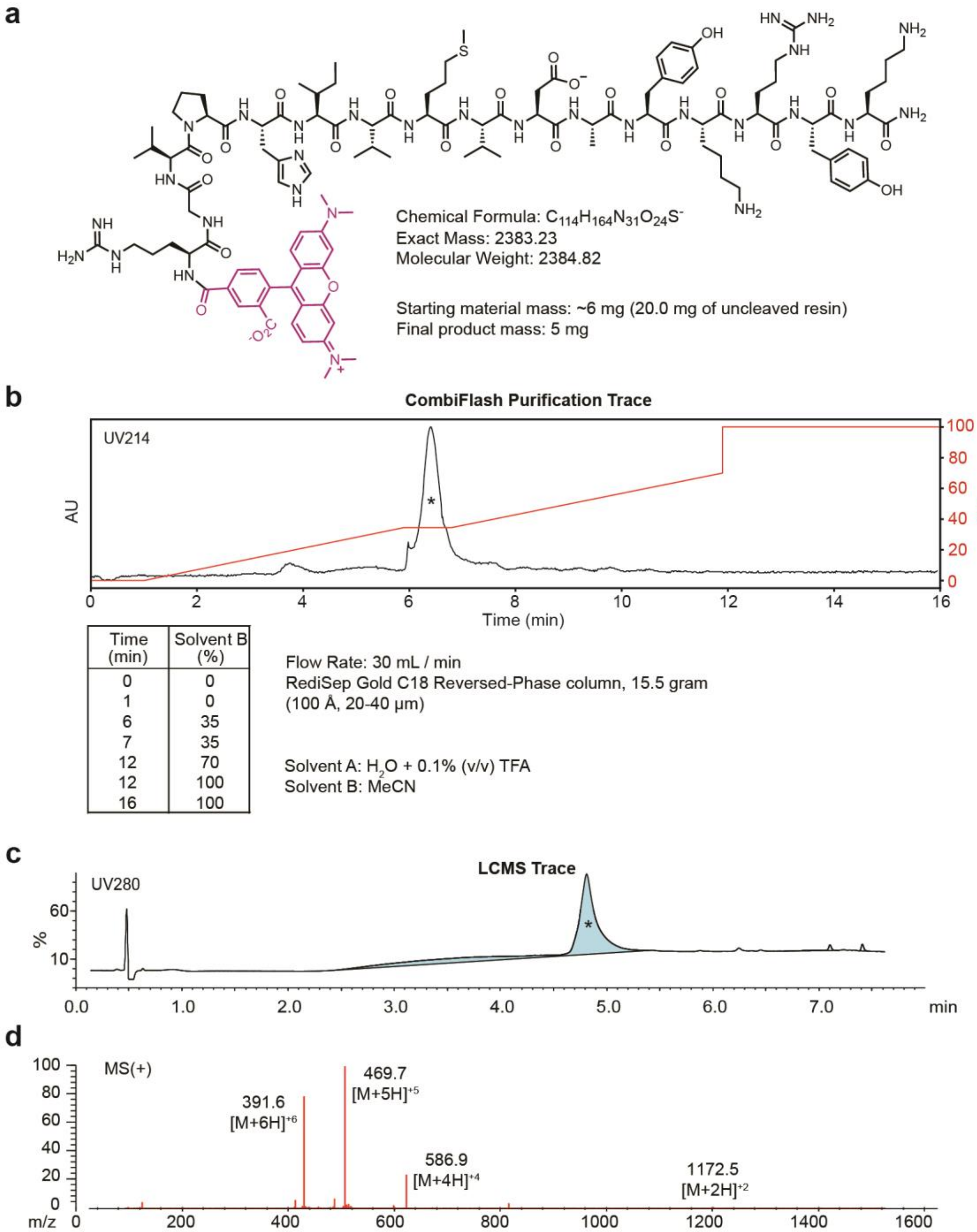

| Time (min) | Solvent B (%) |
|---|---|
| 0 | 0 |
| 1 | 0 |
| 6 | 35 |
| 7 | 35 |
| 12 | 70 |
| 12 | 100 |
| 16 | 100 |

**Supplementary Fig. 11. Synthesis and characterization of TAMRA-ST. a,** Chemical structure of TAMRA-SpyTag **b-d,** Summary of TAMRA-ST purification.
